# Lemonite: identification of regulatory metabolites through data-driven, interpretable integration of transcriptomics and metabolomics data

**DOI:** 10.64898/2026.03.27.714373

**Authors:** Vandemoortele Boris, Devlies Hilde, Tom Michoel, Vanhaecke Lynn, Vandenbroucke Roosmarijn E, Laukens Debby, Vermeirssen Vanessa

## Abstract

Current transcriptomics–metabolomics integration approaches are limited by poor interpretability or incomplete prior knowledge, preventing systematic identification of regulatory metabolites. Here, we present Lemonite, a data-driven, interpretable framework integrating bulk transcriptomics and metabolomics data to uncover regulatory metabolites acting on gene modules. Lemonite extends module network inference to jointly associate transcription factors and metabolites with gene programs, without requiring prior differential analysis or complete metabolome annotation. To contextualize predictions, we built a knowledge graph integrating over 370,000 metabolite–gene/protein and 2.1 million protein–protein interactions. Applied to glioblastoma (n=99) and inflammatory bowel disease (n=75) cohorts, Lemonite identified over 50 functionally coherent gene modules per disease, revealing known and novel metabolite–gene regulatory relationships. In glioblastoma, myo-inositol and phosphatidylcholines, together with IRF6, regulate mesenchymal-like immune programs primarily expressed, per single-cell transcriptomics, in tumor-associated macrophages and monocytes. In inflammatory bowel disease, prioritized regulatory metabolites altered expression of their predicted target genes in colonic epithelial cells in vitro. Overall, Lemonite provides a principled framework to explore the genome-wide regulatory potential of the metabolome and generate interpretable, experimentally testable hypotheses from multi-omics data.

## Introduction

Life on Earth is shaped by the dynamic interplay between different biomolecules that enables cells to maintain homeostasis and respond to changing environments. These molecular networks are tightly interwoven such that signals propagate across multiple molecular layers, thus maintaining homeostasis or orchestrating coordinated responses to perturbation. In this regard, the cellular metabolome has traditionally been considered the endpoint of regulatory cascades, acting as a phenotypic readout and implicitly integrating upstream genomic, epigenomic, transcriptomic and proteomic regulatory phenomena (Cavill *et al*., 2016; Vandemoortele and Vermeirssen, 2023). Today, a paradigm shift emerges as researchers increasingly recognize the regulatory potential of the metabolome, with studies showing that metabolites affect protein stability (Verma *et al*., 2020), protein-protein interaction networks (Schlossarek *et al*., 2022), transcription factor activity (TFA) and specificity (Lopez-Pajares *et al*., 2025), and act as substrates for histone modifications and chromatin reorganization (Sahu and Lu, 2025; Ziogas *et al*., 2025). For example, the presence of citric acid cycle enzymes in the nucleus has been shown to directly influence nuclear acetyl-CoA and histone acetylation levels, having a downstream regulatory effect on several pluripotency genes (Li *et al*., 2022). Such observations call for a multi-modal systems-oriented approach to study gene regulation, where not only nucleic acids and proteins, but also small molecule metabolites are considered.

For this purpose, we define ‘regulatory metabolites’ as small biomolecules originating from metabolic processes within the diet-host-microbiota axis, which have the potential to influence the expression levels of a specific set of target genes or proteins, either directly or indirectly. In contrast to transcriptional regulation of cellular metabolism, limited efforts have focused on uncovering the regulatory potential of the metabolome, especially in high-throughput contexts (Lempp *et al*., 2019; Ortmayr, Dubuis and Zampieri, 2019). While many computational methods exist to model metabolism using transcriptomics or proteomics data (Xiao, Dai and Locasale, 2019; Alghamdi *et al*., 2021; Wagner *et al*., 2021; DeTemmerman, Vandemoortele and Vermeirssen, 2026), many of these tools do not consider metabolomics data itself but leverage transcriptomics or proteomics data as a proxy for metabolic enzyme activity. Consequently, there is currently a lack of both data and data integration approaches that allow studying the genome-wide regulatory potential of the metabolome. This requires the construction of integrated metabolite-gene regulatory networks (GRNs), with nodes representing genes, metabolites and transcription factors (TFs), connected by edges representing putative regulatory interactions between both TFs and genes, and metabolites and genes. These edges are typically inferred in a data-dependent manner, with methods ranging from naïve correlation measures to advanced machine learning (ML) approaches that consider prior knowledge (Loers and Vermeirssen, 2024; Rodriguez-Mier *et al*., 2025).

In practice, existing frameworks that facilitate the integration of transcriptomics and untargeted metabolomics (incl. lipidomics) data can be broadly divided into fully data-driven and prior knowledge-based methods (Cavill *et al*., 2016; Morabito *et al*., 2025). Fully data-driven approaches encompass statistical/correlation-based approaches, multivariate methods, and ML/artificial intelligence (AI) techniques. Dimensionality reduction mitigates the noisy and highly redundant nature of multi-omics data and project features from different modalities into a common subspace. Weighted Gene Correlation Network Analysis (WGCNA) identifies gene-coexpression modules that are characterized by an ‘eigengene’ expression vector, which can be correlated with sample metadata, metabolite abundances or metabolite co-expression modules to identify putative metabolite-gene interactions (Langfelder and Horvath, 2007; Morabito *et al*., 2023). The R package MixOmics contains a suite of statistical and multivariate methods for data integration including the deterministic ‘Data Integration Analysis for Biomarker Discovery using Latent Variable Approaches for Omics Studies’ (DIABLO). DIABLO integrates multi-omics data profiled on the same samples by supervised canonical correlation analysis and partial least squares discriminant analysis, identifying latent components i.e. linear combinations of variables from each omics dataset that maximize covariance across each pair of datasets, including the discretized sample groupings as a dataset to discriminate between them (Rohart *et al*., 2017; Singh *et al*., 2019). MOFA(+) achieves multi-omics integration by inferring a set of latent factors that jointly explain the shared and omics-specific sources of variation across all omics layers in a unified probabilistic framework that disentangles the axes of variability across multi-omics data (Argelaguet *et al*., 2018, 2020). However, latent factors and their associated feature weights are not directly interpretable, and therefore these methods are typically followed by pathway enrichment or clinical covariate association analysis. Lastly, ML/AI- based methods identify hidden underlying patterns in complex datasets to automatically make informed predictions and decisions, supervised or unsupervised, but they require vast amounts of data to build robust models and often provide limited interpretability (Lobentanzer *et al*., 2024; Morabito *et al*., 2025).

In contrast, prior knowledge-based methods integrate data within a curated database, directly generating context-specific, interpretable and testable hypotheses. This database can take the form of either pathways that consist of specific sets of genes and/or metabolites, or integrated metabolite-gene networks that provide topological information. Existing ‘single-omics’ methodologies are often first applied to identify a subset of interesting features from each omics dataset, e.g. differentially abundant transcripts or metabolites, after which these are mapped onto known biochemical pathways and networks, facilitating overrepresentation and integrated network analyses (Dugourd, 2021; Braisted *et al*., 2023; Maan *et al*., 2023; Pang *et al*., 2024). Within the MetaboAnalyst 6.0 framework, joint pathway analysis performs an overrepresentation analysis on an integrated set of genes and metabolites obtained from combined metabolomics and gene expression studies (Pang *et al*., 2024). An important note here is that only a minor proportion of the complete metabolic space can be identified in current metabolomics data analysis, and therefore classical-overrepresentation analysis is unfeasable (Wieder *et al*., 2021). The method COSMOS requires the user to identify a set of genes, proteins and/or metabolites that are deregulated in a given context through classical differential abundance or activity analyses. Using footprinting, the activity of a TF, kinase or phosphatase is estimated from the expression/abundance levels of known target genes and proteins respectively, as such accounting for post-translational modifications of the TF, kinase or phosphatase under study (Badia-i-Mompel *et al*., 2022). Selected features are then mapped onto a meta-prior knowledge network (PKN) which combines multiple sources of experimentally curated causal interactions. COSMOS then detects shortest paths in this meta-network that connect deregulated nodes, thus providing mechanistic hypotheses supported by literature-curated interactions (Dugourd, 2021). In COSMOS+, MOFA(+) is first applied and resulting feature weights are leveraged to select input features for COSMOS+, effectively circumventing the naïve use of single-omics data-analysis methods prior to integration (Dugourd *et al*., 2024). However, these approaches cannot handle biomolecules that are not present in the prior knowledge network, which is a major limitation for (untargeted) metabolomics data since often the majority of metabolites have an unknown or ambiguous identity, for example stereoisomers (Gauglitz *et al*., 2022). Also, the use of prior knowledge-based methods is complicated by discrepancies regarding molecular identifiers for metabolites, which include HMDB, KEGG, ChEBI, InChiKey, SMILES and PubChem IDs among others (Pham *et al*., 2019). Finally, the aforementioned prior knowledge-based methods are limited to detecting known interactions in a context specific manner but cannot identify novel putative interactions from the data.

To circumvent the aforementioned issues and integrate transcriptomics and metabolomics data at population-level in both a data-driven and interpretable way, and put forward “regulatory metabolites” next to other regulatory factors, we developed Lemonite. First, using transcriptomics and metabolomics data of glioblastoma (GBM, n=99) and inflammatory bowel disease (IBD, n=75) patient cohorts, we explored several data-driven and prior knowledge-based data integration methods: MixOmics (Rohart *et al*., 2017), MOFA+ (Argelaguet *et al*., 2020), and COSMOS+ (Dugourd *et al*., 2024). Next, to counteract the limitations of previous methods, we introduced Lemonite (LemonTree for metabolites), an extended adaptation to the open source and modular framework LemonTree that leverages ensemble methods for module network inference (Joshi, Van de Peer and Michoel, 2008; Bonnet, Calzone and Michoel, 2015). Lemonite associates regulatory metabolites and transcription factors to gene modules by integrating bulk-level transcriptomics and metabolomics data. Our methodology not only optimized multi-omics network inference for metabolites, but added biological interpretation through integration with an extensive metabolite- gene knowledge graph (KG) comprising 370 270 unique metabolite-gene and 2 186 811 protein- protein interactions (PPI), respectively. In both GBM and IBD datasets, we prioritized over 50 functional gene modules, with their metabolic and transcriptional regulators, reflecting established and putatively novel disease mechanisms. In glioblastoma, modules showed cell-type specific expression patterns in matched single cell transcriptomics data, and we identified creatinine, 2- hydroxyglutaric acid, myo-inositol, homocysteine, pyrophosphate, phosphatidylcholines and phosphatidylserine/omega-3 fatty acids as metabolic hubs for GBM. In IDB, plasmalogens, carnitine species, sphingomyelins, phosphatidylcholines, gabapentin and putrescine acted as metabolic hubs, targeting multiple gene modules. For IBD, we experimentally validated novel predicted regulatory metabolite-gene interactions, for example the upregulation of *PER3* expression, which is involved in regulation of circadian rhythm, in response to trigonelline, a bioactive compound found in coffee. The Lemonite KG together with IBD and GBM networks presented in this study can be interactively explored at www.lemonite.ugent.be and a NextFlow pipeline is available at www.github.com/CBIGR/lemonite.

## Results

### Exploring existing methods for transcriptomics-metabolomics data integration

To explore the state-of-the-art, we carefully selected two multi-omics cohorts to re-analyze by popular data-driven, linear dimensionality reduction (MixOmics DIABLO, MOFA+) (Rohart *et al*., 2017; Argelaguet *et al*., 2020) and prior-knowledge-based (COSMOS+) (Dugourd *et al*., 2024) data integration methods. GBM is the most common primary malignant brain tumor and metabolic reprogramming is known to fuel tumor progression and therapy resistance (Singh *et al*., 2025). We turned to an extensively multi-omics profiled dataset of 99 treatment-naive GBMs, including tissue transcriptomics, metabolomics and lipidomics, with 4 multi-omics subtypes reported: nmf1 (proneural-like) being enriched for synaptic vesicle cycle, neurotransmitter transport and amino acid metabolism; nmf2 (mesenchymal-like) for innate immune response and glycolysis; nmf3 (classical-like) for mRNA splicing, RNA metabolism, DNA repair and histone acetylation; and finally isocitrate dehydrogenase (IDH) mutated gliomas (L.-B. Wang *et al*., 2021). IBD on its turn, including Crohn’s disease and ulcerative colitis, is a prevalent disease characterized by chronic inflammation of the gastrointestinal tract, accompanied by widespread metabolic dysregulation (Noh *et al*., 2024). Here, we reanalyzed an IBD multi-omics dataset, focusing on 25 ulcerative colitis (UC) and 22 control patients, with 75 colon/rectum biopsies and corresponding stool samples analyzed for transcriptomics and metabolomics respectively (Lloyd-Price *et al*., 2019) (Methods). In both cohorts, we followed the data analysis strategy for metabolomics from the original publications; metabolomics and lipidomics data were treated as separate omics in GBM, and hence metabolites and lipids were kept separate, while in IBD the abundances of both polar and non-polar metabolites were merged into a single metabolomics data file, hence metabolites include lipids here.

Applying DIABLO to GBM bulk transcriptomics, metabolomics and lipidomics data, we identified 6 components with cross-omics correlations > 0.6, with the first 3 components displaying some discriminative behavior between GBM multi-omics subtypes (Supplementary Fig. 1A, Supplementary Data 1). While component 1 differentiated somewhat between nmf1, nmf2 and nmf3 subtypes, component 2 clearly distinguished IDH mutant samples from GBM subtypes and component 3 discriminated to some extent the nmf3 subtype from the other GBM subtypes. Investigation of absolute feature loadings higher than 0.5, revealed a high importance in component 1 for *PACSIN1* and creatinine. *PACSIN1* was previously described as mesenchymal molecular subtype marker with decreased expression in high grade gliomas and being predictive of overall survival (Zimu *et al*., 2021). Creatinine is a degradation product of creatine, which is produced by tumor-associated myeloid cells in hypoxic environments to fuel GBM cells, enhance their stemness and protect them from hypoxia-induced stress (Rashidi *et al*., 2024). Component 2 was primarily driven by 2-hydroxyglutaric acid (2-HG), *FBXO17*, PC(P_18:0/22:6)_B and SM(d18:1/24:0) (Fig. 1A, Supplementary Data 1). 2-HG is an oncometabolite produced as a result of gain-of-function mutations in isocitrate dehydrogenase IDH1/2, which converts α-ketoglutarate to 2-HG, leading to increased methylation and tumor growth, reflecting the IDH-mutant glioma phenotype with increased aerobic glycolysis and deregulated tricarboxylic acid cycle (TCA) metabolism (Han *et al*., 2020). *FBXO17* is highly expressed in GBM, acting as an oncogene that promotes tumor progression, proliferation, plasticity and migration and is one of the genes being downregulated in IDH mutant glioma compared to GBM (N. Wang *et al*., 2021). Component 3 was primarily driven by *FOLR2*, *MGST1* and a metabolite labeled ‘Unknown_017’. *FOLR2* encodes a high-affinity folate receptor primarily highly expressed by myeloid cells within the immunosuppressive tumor microenvironment of GBM (Jiang *et al*., 2025). *MGST1* (Microsomal Glutathione S-Transferase 1) is upregulated in glioma tissues and functions as an oncogene that promotes tumor progression, migration, and chemoresistance, inhibiting oxidative stress and ferroptosis (Yan *et al*., 2024). Although gene set enrichment analysis (GSEA) using MixOmics DIABLO feature weights can functionally characterize components, no significantly enriched pathways were identified in any of these 3 components (Supplementary Data 1).

**Figure 1.**
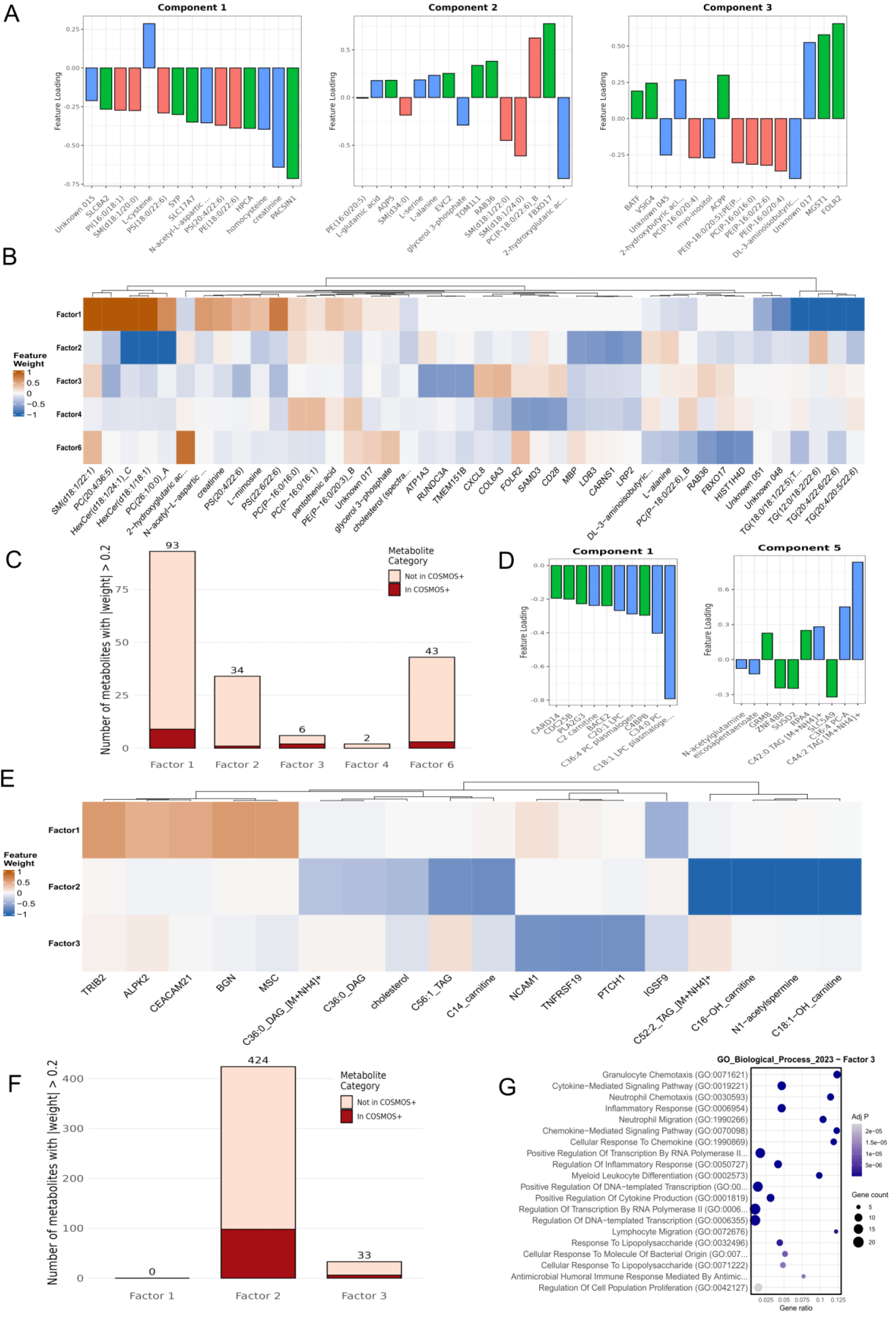
Exploratory analysis of GBM and IBD patient cohorts through integration of metabolomics, lipidomics (GBM-only) and transcriptomics data. **A.** Feature weights for the GBM MixOmics DIABLO model showing the top 5 features in lipidomics (red), metabolomics (green) and transcriptomics (blue) data in components 1-3 with cross-omics correlations > 0.7. **B**. Feature weights of the GBM MOFA+ model showing top 3 features in lipidomics, metabolomics and transcriptomics data for factors 1,2,3,4 and 6. **C.** COSMOS+ analysis on GBM MOFA factors. Bar height (light red) represents the number of metabolites with absolute feature weight > 0.2 per MOFA+ factor, colored part (dark red) represents the number of those metabolites that can be mapped onto the COSMOS+ meta-network. **D.** Feature weights for the IBD MixOmics DIABLO model showing the top 5 features in metabolomics (blue) and transcriptomics (green) data in components 1 and5. **E**. Feature weights of the IBD MOFA+ model showing top 3 features in metabolomics and transcriptomics data for factors 1-6. **F.** COSMOS+ analysis on IBD MOFA factors. Bar height (light red) represents the number of metabolites with absolute feature weight > 0.2 per MOFA+ factor, dark red bar represents the number of those metabolites that can be mapped onto the COSMOS+ meta-network. **G.** Overrepresentation analysis on nodes in the COSMOS+ network for MOFA+ factor 3 in IBD.. DIABLO: Analysis for Biomarker Discovery using Latent Variable Approaches for Omics Studies; MOFA+: Multi-Omics Factor Analysis+. Meta- PKN: COSMOS+ meta prior knowledge network. Feature labels show the first 20 characters to enhance readability (listed in Supplementary Table 1).

On the same dataset, MOFA+ identified 15 latent factors. Factors 1, 2, 3, 4 and 6 displayed differential sample loadings between the GBM multi-omics subtypes (p-adj < 0.05, Kruskal-Wallis test, Supplementary Fig 1B), although only factors 5 and 6 captured > 1% of the total variance across all three omics (Supplementary Table 2). Factor 1 was primarily driven by lipids and capturing 19.78% of the total variance in lipidomics data, with several triacylglycerols (TG(18:0/18:1/22:5);TG(18:0/18:0/22:6), TG(12:0/18:2/22:6), TG(20:4/22:6/22:6) and TG(20:4/20:5/22:6)) having a strong negative contribution , while also displaying lower sample loadings in IDH mutant glioma compared to other GBM subtypes (Supplementary Data 1). Factor 2 displayed higher sample loadings in the IDH mutant subtype and strong negative loadings for HexCer(d18:1/24:1)_C, PC(26:1/0:0)_A and PC(26:1/0:0)_A, next to *MBP*, *LDB3*, *CARNS1* and *LRP2*. While *CARNS1* does not have an established connection with lipid metabolism, MBP is a major structural component of myelin and LRP2 acts as an endocytic receptor involved in the uptake of extracellular lipoproteins (Christensen and Birn, 2002). Factor 3 was primarily driven by *ATP1A3*, *RUNDC3A* and *TREM151B*, next to *CXCL8* and *COL6A3* that have established roles in GBM as a microenvironmental cytokine or extracellular matrix remodeling respectively (Long *et al*., 2025). *ATP1A3* is emerging as a therapeutic target due to its tumor suppressive role next to modulating sensitivity to chemotherapy (Zou *et al*., 2025). Classical-like subtype samples displayed higher weights in factor 4, to which *FOLR2*, *SAMD3* and *CD28* contributed negatively and PC(P-16:0/16:0), PC(P-16:0/16:1) and PE(P-16:0/20:3)_B positively. Finally, factor 6 clearly displayed lower sample loadings in IDH-mutant glioma, and was primarily driven by 2-HG, *RAB36*, *FBXO17* and *HIST1H4D*. IDH mutant gliomas are characterized by severe epigenetic rewiring, such as changes to histone acetylation and methylation including HIST1H4D (Turcan *et al*., 2018), while *RAB36* is overexpressed in GBM compared to healthy brain tissue (Han *et al*., 2024).

MOFA+ factors can be functionally characterized by ranking genes based on feature weights in the respective factors and then applying GSEA (Supplementary Data 1). Genes in factor 1 were involved in myelination and axon ensheathment, in line with the factor primarily being driven by triacylglycerols. Factors 3 and 6 displayed a positive enrichment for inflammation-related pathways, while factor 5 was downregulated for ‘T cell selection’ and IL12 production (Supplementary Data 1).

In addition to these purely data-driven methods, we integrated MOFA factor feature weights with the COSMOS+ prior knowledge meta-network, which comprises 117,065 directed interactions between 3,495 metabolites and 19,530 genes (Dugourd et al., 2024). After filtering for metabolites with absolute weight > 0.2 in each factor, less than half could be mapped onto the COSMOS+ meta-network (Fig. 1E, 9/93 in factor 1, 1/43 in factor 2, 2/6 in factor 3, 0/2 in factor 4 and 3/40 in factor 6). COSMOS+ subsequently constructed a subgraph that can provide mechanistic hypotheses by connecting the selected nodes in the meta-network through causal signaling interactions, however, the final networks were limited to TFs and target genes, and did not contain any metabolites (Supplementary Data 1).

Similarly, on the IBD cohort DIABLO inferred 5 latent components, with components 1 and 5 capturing a cross-omics correlation > 0.6, and only component 1 displayed discriminative power (Supplementary Fig. 1C). Component 1 was primarily driven by C18:1 LPC plasmalogen, but GSEA did not reveal any significantly enriched pathways (p-adj < 0.05, GO biological process) (Supplementary Data 1). On the same dataset, MOFA+ uncovered 15 latent factors, with factors 1, 2 and 3 displaying differential sample loadings between IBD and controls (p-adj < 0.05, Kruskal- Wallis test, Supplementary Fig. 1D, Supplementary Data 2). Factor 1 was primarily driven by *TRIB2*, *ALPK2*, *CEACAM21*, *BGN*, *MCS* and *IGSF9*. TRIB2 is a negative regulator of TLR5- mediated NF-kB signaling and is decreased in actively inflamed IBD lesions (Wei *et al*., 2012), while BGN contributes to intestinal fibrosis through BMP-7 mediated signaling (M. Yu *et al*., 2025).

Factor 2 was primarily driven by C52:2_TAG_[m+nh4]+, C16-OH_carnitine, N1-acetylspermine and C18:1-OH_carnitine, with N1-acetylspermine being part of the polyamine pathway which is altered in IBD (Lloyd-Price *et al*., 2019). Factor 3 finally was primarily driven by *NCAM1*, *TNFRSF19* and *PTCH1*, the latter being a TF upstream of hedgehog signaling that controls intestinal development and homeostasis (Walton and Gumucio, 2021) (Supplementary Data 2). COSMOS+ networks were next constructed (identical approach as in GBM analysis), although limited numbers of metabolites passed weight-based selection and mapping onto the meta- network (Fig. 1F). Subsequently, the network for factor 3 was the only network in which at least 1 metabolite was retained (11,14-eicosadienoyl-CoA), comprised of 45 nodes in total, and was enriched for several inflammation-related pathways such as granulocyte chemotaxis and cytokine-mediated signaling pathway(Fig. 1G, Supplementary Data 2).

Data-driven methods MixOmics and MOFA+ provided valuable multi-omics insights. Projecting samples into an integrated low-dimensional space enabled clustering of GBM multi-omics subtypes and separation of UC patients from controls using transcriptomics and metabolomics data. Both methods consistently prioritized similar metabolites and genes, such as 2-HG, FOLR2 and FBXO17 in GBM and C18:1 LPC plasmalogen in IBD. Although these approaches identified important biomolecules associated with specific components or factors related to the disease, they did not provide mechanistic insights or enable the construction of an integrated network of metabolism and gene regulation The prior-knowledge-based method COSMOS+ on the other hand was limited in the number of metabolites that were retained after filtering based on MOFA+ feature weights, and that could subsequently be mapped onto the meta-network, and were retained in the final network solutions.

### Lemonite is an interpretable, data-driven multi-omics integration methodology that identifies regulatory metabolites and target genes

To overcome the aforementioned limitations, we developed a multi-omics data integration strategy tailored towards, but not limited to, bulk transcriptomics and metabolomics datasets by building on the existing module network inference software suite LemonTree (Bonnet, Calzone and Michoel, 2015). First, transcription factor activities (TFAs) were calculated on normalized, highly variable gene selected, scaled transcriptomics data using DecouplR and the CollecTri TF-target gene network encompassing 43 175 signed interactions for 1 186 TFs (Methods) (Badia-i-Mompel *et al*., 2022; Müller-Dott *et al*., 2023). By default 100 independent model-based Gibbs samplers simultaneously inferred co-expression clusters from transcriptomics data, after which the results were combined into consensus modules through spectral edge clustering (Bonnet, Calzone and Michoel, 2015). Per regulator type (TF, metabolite, lipid, other), an ensemble of regulatory programs was inferred non-linearly for each module based on decision trees leveraging TFA (or TF expression levels) and normalized and scaled metabolite and lipid abundances (or other omics if provided). For each module, regulator consensus scores were put forward, together with a random regulator score, calculated based on an empirical score distribution. As prioritization of computational predictions is a major challenge, we made module and regulator prioritization explicitly available in the Lemonite pipeline. First, modules that displayed weak coexpression patterns were removed from the analysis and modules were tested for enrichment of known PPIs involving module genes. Second, modules were prioritized based on sample group discrimination by differential expression. Third, modules were selected based on functional overrepresentation or gene set enrichment analysis for specific genes or pathways of interest and/or the presence of PPI or other functional gene-gene links. Finally, modules were selected based on the assignment of high-scoring and highly-connected regulators (with many target modules) and/or regulators for which the biological role corresponded to the module function. In this way, Lemonite builds a data- driven, integrated metabolite-TF-module network, in which regulator-module pairs represent functional gene modules that are regulated by a ranked list of scored metabolites and TFs. The complete Lemonite (LemonTree for metabolites) pipeline, including data preprocessing, network inference, downstream analyses and visualizations, is available as a NextFlow pipeline on GitHub (https://github.com/CBIGR/Lemonite/nextflow). We note that although Lemonite was primarily developed as a tool for integrating transcriptomics and metabolomics data, it can be applied to other omics data types.

### Construction of an integrated gene/protein-metabolite knowledge graph

Metabolites can interact with genes and proteins in several ways facilitating the crosstalk between metabolism and gene expression (Hornisch and Piazza, 2025). To facilitate the interpretation of, and to in silico validate, our predicted integrated metabolite-TF module network, we gathered known gene/protein-metabolite interactions from different resources. We will term them metabolite-gene interactions from now on to enhance readability. Despite the fact that substantial efforts have been devoted to the curation of metabolite-gene interactions (Braisted *et al*., 2023; Farr *et al*., 2024; X. Wang *et al*., 2025), prior knowledge on metabolite-gene interactions is scattered across multiple databases and platforms, obstructing ease-of-access and high-throughput usability by researchers. Moreover, databases often use a different type of ID for metabolites, limiting comparison and integration of information across databases. We therefore parsed the human metabolic database (HMDB) and retrieved 217 920 unique metabolites, of which 13 562 were annotated with a ChEBI ID, 5 908 with a KEGG ID, 103 682 with a PubChem ID, 217 895 with an InchIKey and 217 460 with a SMILES ID (Table 1, Fig. 2A). Leveraging these identifiers, we first queried the BioGRID (Oughtred *et al*., 2021), UniProt (Bansal *et al*., 2022; The UniProt Consortium, 2023), IntAct (Orchard *et al*., 2014), chEMBL (Zdrazil *et al*., 2024), LINCS (Keenan *et al*., 2018) and STITCHdb (Szklarczyk *et al*., 2016) databases to retrieve curated metabolite-gene interactions (Table 1). Second, we queried the Human1 genome-scale metabolic model (GEM) for metabolite-enzyme interactions, which we mapped to metabolite-gene interactions using the GEM’s gene-protein-reaction rules (Robinson *et al*., 2020). Since metabolic regulation frequently encompasses feed-back and feed-forward loops involving substrate or product metabolites and enzymes, pairs of neighboring reactions in the metabolic network were collapsed into a single reaction (Berg *et al*., 2023). As such, we considered Human1 metabolite-gene interactions to comprise all genes that can be reached within two reactions in the metabolic network from a given metabolite, after removing hyper-connected metabolites (Methods). Third, we retrieved all metabolite-gene interactions from MetalinksDB, a recently published intercellular metabolite-gene interaction database focused on signaling interactions (Farr *et al*., 2024). We found surprisingly limited overlap between databases, with each database contributing a set of mostly unique interactions (Fig. 3B). Together, these data were combined into a single metabolite-gene PKN network comprising 8 849 unique metabolites identified by an HMDB ID, 16 719 unique genes and 371 270 unique metabolite-gene interactions (Fig. 3B). We next extended this metabolite- gene-gene network with PPI data from STRINGdb (Szklarczyk *et al*., 2015), the Human Reference Interactome (HuRI) (Luck *et al*., 2020), BIOGRID (Oughtred *et al*., 2021) and HumanNet (Kim *et al*., 2022), by retrieving all experimental, physical PPIs involving at least one of 16 719 unique genes (i.e. the encoded protein) in our metabolite-gene network, resulting in a total of 2 488 214 unique PPIs (Fig. 3C). The Lemonite knowledge graph (KG) encompasses 2 859 484 unique molecular interactions between 8 849 and 32 549 unique metabolites and genes/proteins respectively. The KG was designed to be comprehensive, integrating experimentally supported molecular interactions from multiple curated resources, with metabolite–gene/protein interactions classified as causal (LINCS, ChEMBL), metabolic pathway–based (Human1-GEM), or ‘other’ interaction types (IntAct, UniProt, BioGRID, STITCHdb, MetalinksDB) (Table 1).

**Figure 2.**
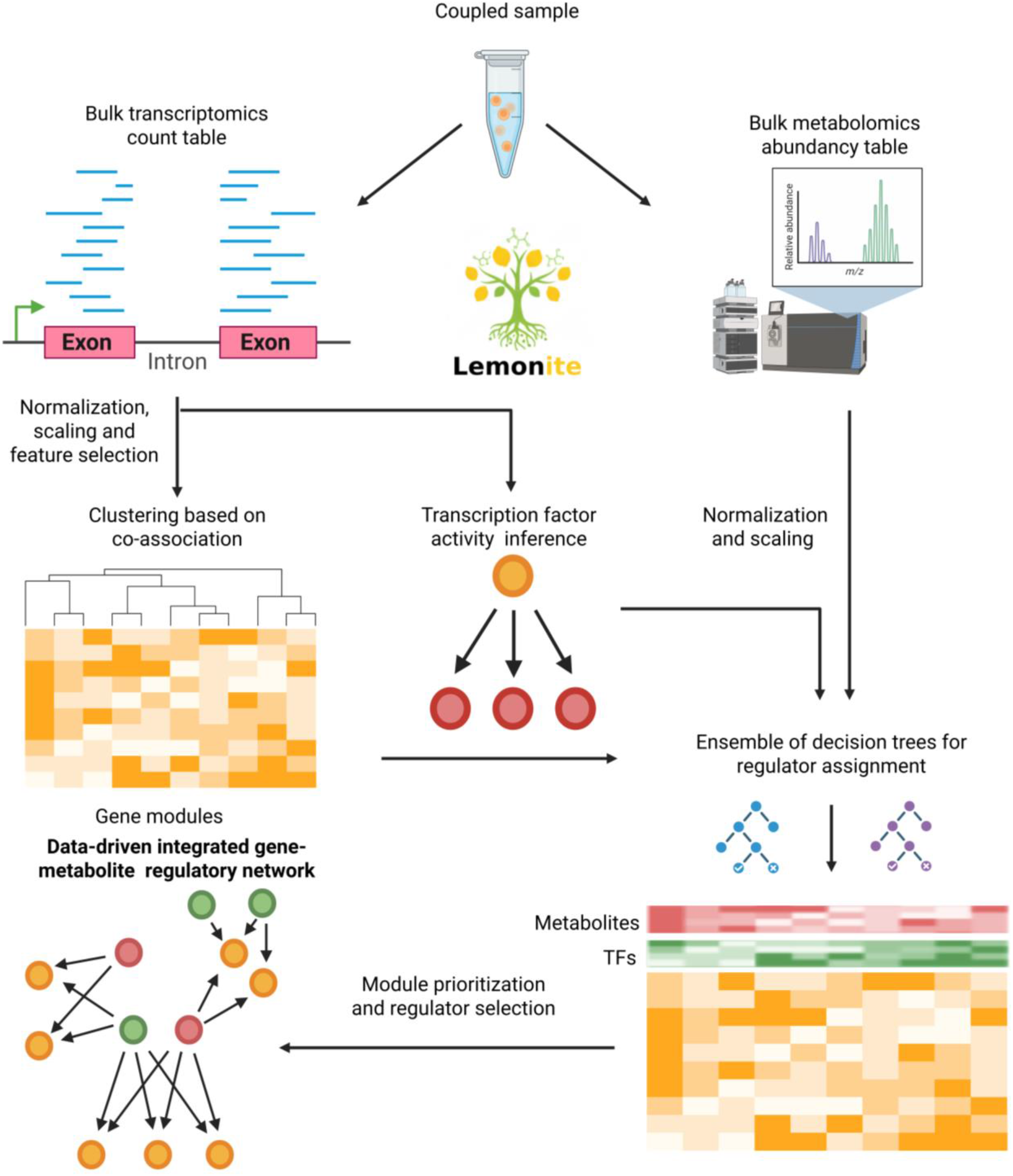
Lemonite performs data-driven, interpretable integration of bulk transcriptomics and metabolomics data. Lemonite requires coupled data as input and was developed for the integration of bulk transcriptomics and metabolomics data, but also accepts other omics data types (proteomics, phosphoproteomics, copy number variation, microbiomics, epigenomics…). Data are preprocessed, genes are clustered based on co-association into gene modules and transcription factor activities (TFA) are inferred. Regulatory programs are then predicted for each gene module by an ensemble of decision trees leveraging TFA scores (of TF expression if TFA not available) and metabolite abundances (and abundances of additional omics if provided). Lemonite then selects the most relevant regulators based on rank and score and prioritizes gene modules with their respective regulators based on module coherence (i.e. coexpression), module differential expression, PPIs within gene modules, functional annotation analysis on module genes and regulators, regulator-to-module scores and known interactions involving regulators and module genes (see next section and Methods).

**Figure 3.**
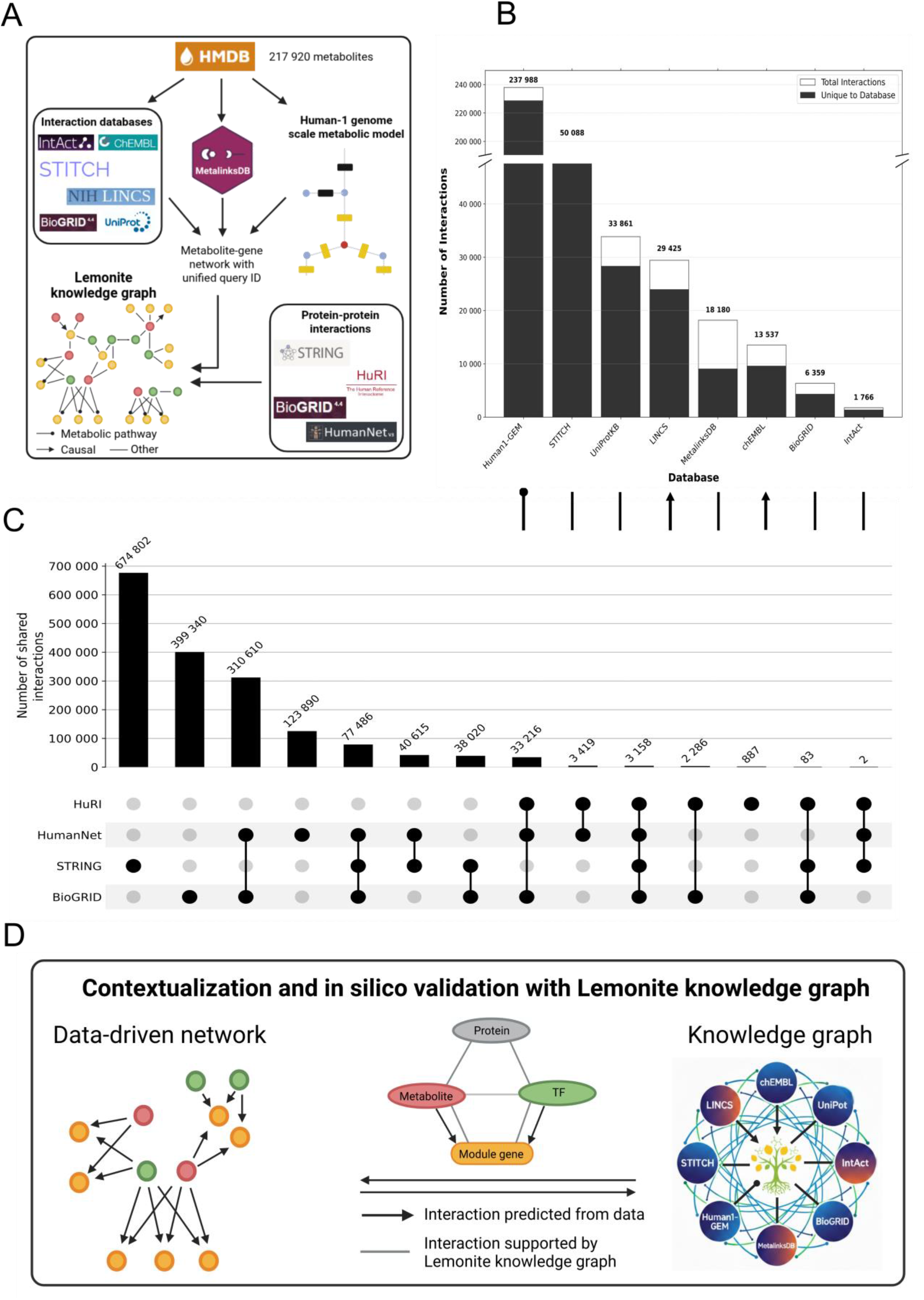
The Lemonite knowledge graph. **A.** The Lemonite knowledge graph (KG) contains metabolite-protein and metabolite-gene interactions queried from UniProt, IntAct, LINCS, chEMBL, BioGRID, STITCHdb, Human1-GEM and metalinksDB for metabolites from the Human Metabolome Database (HMDB). Interactions are annotated as causal (LINCS & chEMBL, arrow), metabolic pathway (Human1-GEM, dot) or other (straight line). These were complemented with protein-protein interactions retrieved from STITCH, BioGRID, the Human Reference Interactome (HuRI) and HumanNet. **B.** Number of metabolite-gene interactions retrieved from IntAct, BioGRID, chEMBL, UniProt, STITCH, LINCS, Human1-GEM and MetalinksDB. Black bar represents the number of interactions that are unique to each database, white bar represents to total number retrieved from each database. **C.** Intersection between protein-protein (PPI) interactions from STRING, BioGRID, HuRI and HumanNet. **D.** Lemonite performs contextualization and in silico validation of its data-driven metabolite-TF-module network trough comparison with the Lemonite knowledge graph. True metabolite-module gene and PPI interactions are displayed on module figures and can be considered for prioritization of modules and predictions.

**Table 1.** Databases with metabolite-gene-gene or protein-metabolite interactions included in the Lemonite KG. Databases have varying scopes and were queried using different metabolite identifiers. ‘Metabolite identifier’ indicates which type of identifier must be used to query a given database. ‘HMDB metabolites in database’ shows how many metabolites from HMDB are present in a given database, after parsing the database with the identifier provided by HMDB. ‘Interactions involving HMDB metabolites’ represents the number of unique interactions that can be retrieved for all metabolites present in HMDB with their identifiers. Interactions were classified into three distinct categories based on database content: causal (LINCS and chEMBL), metabolic pathway (Human1-GEM) and other (BioGRID, UniProt, STITCHdb, IntAct, MetalinksDB). HMDB: Human Metabolome Database.

| Database | Metabolite identifier | HMDB metabolites in database | Interactions involving HMDB metabolites | Database content | Interaction type |
|---|---|---|---|---|---|
| **BioGRID (Oughtred *et al.*, 2021)** | InChIKey | 1 672 | 6 359 | Biomedical interaction repository – interactions curated from scientific literature | Other |
| **UniProt (The UniProt Consortium, 2023)** | InChIKey | 2 517 | 32 708 | Protein structure database – physical protein-metabolite interactions | Other |
| **STITCHdb (Szklarczyk *et al.*, 2016)** | PubChem ID | 1 946 | 50 126 | Chemical-protein interaction database – mostly metabolite-receptor interactions | Other |
| **IntAct (Orchard *et al.*, 2014)** | ChEBI ID | 340 | 1 758 | Experimentally observed physical interactions | Other |
| **chEMBL(Zdrazil *et al.*, 2024)** | Canonical SMILES | 2 749 | 13 135 | Experimental assays with measured effect on protein target – causal interactions | Causal |
| **LINCS (Keenan *et al.*, 2018)** | Canonical SMILES | 2 884 | 29 425 | Changes in gene expression and other cellular processes that occur when cells are exposed to a variety of perturbing agents – causal interactions | Causal |
| **Human1 (Robinson *et al.*, 2020)** | ChEBI ID | 984 | 1 704 364 | Metabolite-enzyme interactions and gene-protein-reaction rules | Metabolic pathway |
| **MetalinksDB (Farr *et al.*, 2024)** | HMDB ID | 968 | 18 180 | Allosteric metabolite-protein interactions, metabolite enzyme sets | Other |

In the Lemonite pipeline, the Lemonite KG serves a dual role. First, it performs *in silico* validation of predicted metabolite-module interactions by evaluating the presence of KG interactions in the data-driven Lemonite network. Second, it enhances biological interpretation by providing molecular context for predicted TFs, metabolite regulators and module genes. The annotation of edges in the Lemonite knowledge graph as ‘causal’, ‘metabolic pathway’ or ‘other’ enhances the interpretability of predicted interactions and allows prioritization of regulator-module pairs. PPIs in the Lemonite KG further allow to identify potential co-regulatory mechanisms between pairs of TFs and metabolites that are associated to the same target genes, potentially reflecting allosteric regulation of TFs through interaction with a metabolite. Also, the PPIs present in the KG allow to directly test if gene modules are enriched for PPIs, as such identifying functionally coherent modules. The Lemonite KG is available as part of the Lemonite Nextflow pipeline and as an online searchable web application at www.lemonite.ugent.be, which also connects to the original databases where the interaction was retrieved to enhance interpretability, information content and contextualization for the end user. Users can also upload their own clustering and regulator data and perform *in silico* validation with the Lemonite KG. The construction of the KG has been implemented in a computational pipeline that allows regular updates.

### Lemonite infers an integrated metabolite-TF-module network in GBM

We integrated transcriptomics, metabolomics and lipidomics data from primary GBM tumor tissues using Lemonite and identified 63 modules with metabolite, TF and lipid regulators, of which 46 were retained after filtering out modules with weak coexpression, after which also the most relevant regulators were selected as described in the Methods section (Supplementary Fig. 2A) The resulting integrated metabolite-TF-module network comprised of 1930 genes, 48 metabolites, 157 lipids and 111 TFs together spanning 5018 metabolite-gene, 9 969 lipid-gene and 12 866 TF-gene interactions (Supplementary Data 1. Analysis of the regulator hubs, nodes with high outgoing-degree in the network, revealed high regulatory relevance for several TFs known to be relevant in GBM. IRF6 functions as a tumor suppressor in GBM and gliomas, while overexpression induces apoptosis, inhibits glioma cell proliferation and impairs glycolysis (Subbalakshmi *et al*., 2026). *SOHLH1* is an essential TF in germline differentiation, but has recently been identified as a tumor suppressor gene by inhibiting Wnt signaling in glioma stem- like cells (Zhi *et al*., 2025). Also *MYT1L* functions as a tumor suppressor in GBM cells by repressing *YAP1* and activating neuronal differentiation gene programs (Melhuish *et al*., 2018). *NKX6-2* is involved in nervous system development and oligodendrocyte maturation, but currently has no known role in GBM. *EMX1* finally functions as a tumor inhibitor in spinal cord glioma, but its role in GBM remains to be established (Han *et al*., 2022) (Fig 4A, Table 2). The top two lipid hub regulators were phosphatidylcholines, suggesting high relevance for this group of lipids in GBM as was also described in the original publication (Fig. 4A, Table 2) (L.-B. Wang *et al*., 2021). PS(22:6/22:6) ranked third, a highly unsaturated phosphatidylserine with 2 docosahexaenoic acid acyl chains involved in membrane remodeling that may affect sensitivity of GBM cells to ferroptosis (Mitre *et al*., 2022; Qiu *et al*., 2024). Moreover, oncometabolite 2-HG ranked second among highly connected hub metabolites, preceded by creatinine which has been implicated in GBM through its closely related product creatine (which was absent in this dataset) (Rashidi *et al*., 2024) (Fig. 4A, Table 2). Further, both myo-inositol and homocysteine, which were ranked third and fourth respectively, have been reported as potential serum biomarkers in GBM (Djurovic *et al*., 2020; El-Abtah *et al*., 2022), and myo-inositol has further been documented to decrease oxygen binding by hemoglobin, causing an increase in oxygen release in hypoxic tumor with high myo- inositol levels (Grgic *et al*., 2021). Interestingly, several modules were targeted by the same set of regulators, such as myo-inositol and IRF6 targeting modules 5, 24 and 50, while each of these modules was characterized by an enrichment of immune-related pathways (Fig. 4A, p-adj < 0.05, Fisher exact test, Supplementary Data 1). We next prioritized modules based on differential expression between subtypes (42 out of 46 with p-adj < 0.05, Kruskal-Wallis test), enrichment for PPIs (30 out of 46 with p-adj < 0.05, hypergeometric test) and functional enrichment analyses (Fig. 4B, Supplementary Data 1).

**Figure 4.**
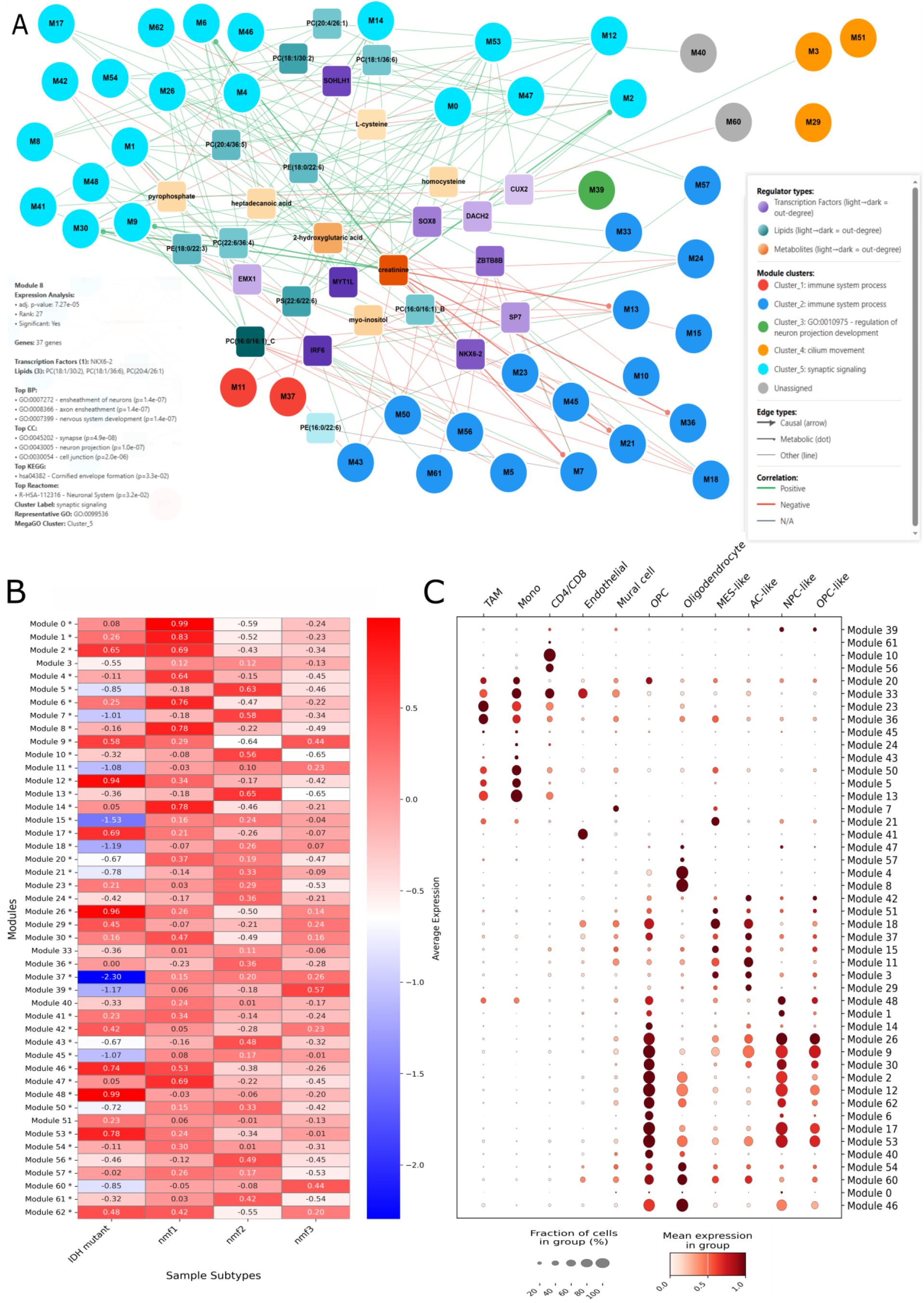
Lemonite builds an integrated metabolite-lipid-TF-module network in GBM. **A.** Lemonite GBM network containing 46 gene modules and most important metabolite, lipid and TF regulators (Methods). Gene modules are colored by and labeled GO enrichment group (Methods), regulators are colored by type and out-degree. Edges end in a dot or arrow if the regulator-module connection captures known metabolic pathway or causal interactions resp. Edge color represents the sign of correlation between regulator expression and module expression. The complete network can be explored and searched interactively at www.lemonite.ugent.be. Mouse hover (bottom left) shows regulators for each module, number of genes in a module, differential expression statistics, and a summary (top 3 pathways) of functional enrichment results. **B.** Heatmap colored by average gene expression per module per multi-omics subtype. * indicates significantly differentially expressed modules (Kruskall-Wallis test, p-adj < 0.05). **C.** Expression of Lemonite modules in matched single cell data (n=45 144 cells). Dot color indicates the mean scaled expression of module genes in a given group of cells (grouped per cell type), dot size represents the proportion of cells within a group that has non-zero expression of module genes.

**Table 2.** Hub regulators in the integrated metabolite-TF-module GBM network. Top 5 regulators for TFs, metabolites and lipids, ranked by ‘Combined score’. Combined score represents the sum of all regulator-module scores for each regulator, ‘# Target modules’ is the number of modules each regulators targets, and ‘# Target genes’ is the total combined number of genes across all modules targeted by a given regulator.

| **Regulator** | **Combined score** | **# Target modules** | **# Target genes** |
|---|---|---|---|
| **Creatinine** | 195.39 | 16 | 971 |
| **2-hydroxyglutaric acid** | 138.56 | 11 | 600 |
| **Myo-inositol** | 53.31 | 6 | 219 |
| **Homocysteine** | 48.50 | 4 | 164 |
| **Pyrophosphate** | 31.21 | 5 | 169 |
| **PC(16:0/16:1)_C** | 60.35 | 11 | 514 |
| **PC(20:4/36:5)** | 58.92 | 6 | 508 |
| **PS(22:6/22:6)** | 33.16 | 8 | 383 |
| **PE(16:0/18:1)** | 28.50 | 3 | 112 |
| **PE(18:0/22:6)** | 26.92 | 6 | 479 |
| **SOHLH1** | 76.83 | 7 | 481 |
| **IRF6** | 68.90 | 10 | 451 |
| **NKX6-2** | 68.57 | 6 | 149 |
| **MYT1L** | 56.42 | 9 | 595 |
| **EMX1** | 31.27 | 4 | 362 |

Module 0 was significantly upregulated in proneural-like nmf1 samples compared to other multi- omics subtypes (p-adj 1.23e-7, Kruskal Wallis H test, Supplementary Fig 3A, Supplementary Data 1). The module was biologically functionally coherent as indicated by a significant enrichment of several neurotransmitter-related pathways, similar to what has been described for the nmf1 multi- omics subtype characterization in the original paper (p-adj < 0.05, GSEA, Supplementary Fig. 3B, Supplementary Data 1) (L.-B. Wang *et al*., 2021) and a significant enrichment of known PPIs (p-adj 2.72e-188, hypergeometric test) (Supplementary Fig. 3A, Supplementary Data 1). In addition to the first regulator creatinine, the second regulator L-lysine coincides with dysregulated amino acid metabolism for the proneural subtype (L.-B. Wang *et al*., 2021), and has been associated with increased histone crotonylation and reprogrammed immunity in GBM (Yuan *et al*., 2023). The majority (4/6) of lipid regulators were phosphatidylcholines (PCs), with PS and PE also being member of the broader phospholipid class. These lipid regulators showed lower abundance in nmf2 samples, and accordingly nmf2 samples displayed lower expression of genes in module 0 (Supplementary Fig. 3). TF regulators for module 0 included EMX1, SOHLH1 and MYT1L, and which displayed higher activities in nmf1samples. MYT1L actively represses non-neuronal cell fates and induces neuronal differentiation in neural stem cells (Mall *et al*., 2017), and also SOHLH1 and EMX1 have been implicated in GBM (see above). Interestingly, each of these TFs could be connected to the regulator metabolites through a single protein-protein interaction in the Lemonite KG. For example, L-lysine could be connected to MYT1L through a PPI with PXDN, an extracellular matrix-associated protein that has been implicated in poor prognosis for GBM (Shi *et al*., 2022) (Supplementary Fig. 3C).

Module 5 on the other hand was significantly upregulated in mesenchymal-like nmf2 samples compared to other multi-omics subtypes (p-adj 3.95e-6, Kruskall Wallis H test, Supplementary Fig 4A, Supplementary Data 1) and enriched for known PPIs (p-adj 1.53e-240, hypergeometric test). This functional module was involved in ‘inflammatory response’, and ‘’Neutrophil chemotaxis’ (p- adj < 0.05, GSEA, Supplementary Fig. 4B, Supplementary Data 1), in line with the initial study’s description of nmf2 samples being enriched for in the ‘innate immune response’ and ‘neutrophil degranulation’ (L.-B. Wang *et al*., 2021). Lemonite identified myo-inositol and several phosphatidylcholines as ‘negative’ metabolic and lipid regulators, next to IRF6 as ‘positive’ top transcriptional regulator. All top 6 lipid regulators of module 5 were phosphatidylcholines, suggesting high relevance for this class of lipids in mesenchymal-like GBM, as was also described in the original study (L.-B. Wang *et al*., 2021). IRF6 is involved in immune response and metabolic stress regulation and has been implicated in epithelial-mesenchymal transition (EMT) as well (Li *et al*., 2019, p. 6; Kim *et al*., 2024, p. 6). Interestingly, myo-inositol has anti-inflammatory effects and can reverse the transformation of epithelial cells into a mesenchymal-like phenotype, and causes an increase of cell motility in GBM cells, together suggesting a regulatory effect of myo-inositol during the process of EMT (Brosch *et al*., 2022; Rashidi *et al*., 2024). Thus, myo-inositol and phosphatidylcholine abundances negatively correlated with module gene expression, whereas the activity of IRF6 positively correlated with the gene module. Furthermore, *TGF-β1* is member of module 5, and it is known that myo-inositol can revert *TGF-β1* EMT in a model of non- tumorigenic breast cells (Monti *et al*., 2023). Furthermore, module 5 recovered a known interaction between myo-inositol and ANPEP, a membrane-bound Zn metalloproteinase involved in cell migration and thus further supporting the involvement of this module and its regulators in EMT (Ghosh and Shapiro, 2019).

### Lemonite recapitulates a connection between 2-HG, methionine metabolism and DNA methylation through branched-chain amino acid metabolism in GBM

Module 18 comprised genes that were collectively downregulated in IDH-mutant and upregulated in mesenchymal-like samples (p-adj 3.74 e-4, Kruskal–Wallis H test), and was significantly enriched for PPIs (p-adj 3.74 e-4, hypergeometric test) (Fig. 5A, Supplementary Data 1). Top metabolite regulators included 2-HG and L-methionine. Consistent with the biology of IDH mutations, 2-HG abundance showed a strong positive correlation with module gene expression, whereas L-methionine abundance was anticorrelated. Module 18 retrieved known interactions between metabolite regulators and module genes, involving BCAT1 and INMT (Fig. 5B). BCAT1 catalyzes reversible reactions between branched-chain amino acids and their corresponding keto acids using α-ketoglutarate (α-KG) as a cofactor. Methionine metabolism contributes to the trans- sulfuration pathway via α-KG or enters the methionine cycle to generate SAM, the major methyl- group donor for chromatin modifications, whereas α-KG-dependent demethylases are inhibited by 2-HG (Sahu and Lu, 2025). Together, these connections suggest that altered α-KG availability, oncometabolite accumulation, and methionine metabolism may jointly influence module 18 regulation and contribute to broader changes in DNA and histone methylation states in glioma.

**Figure 5.**
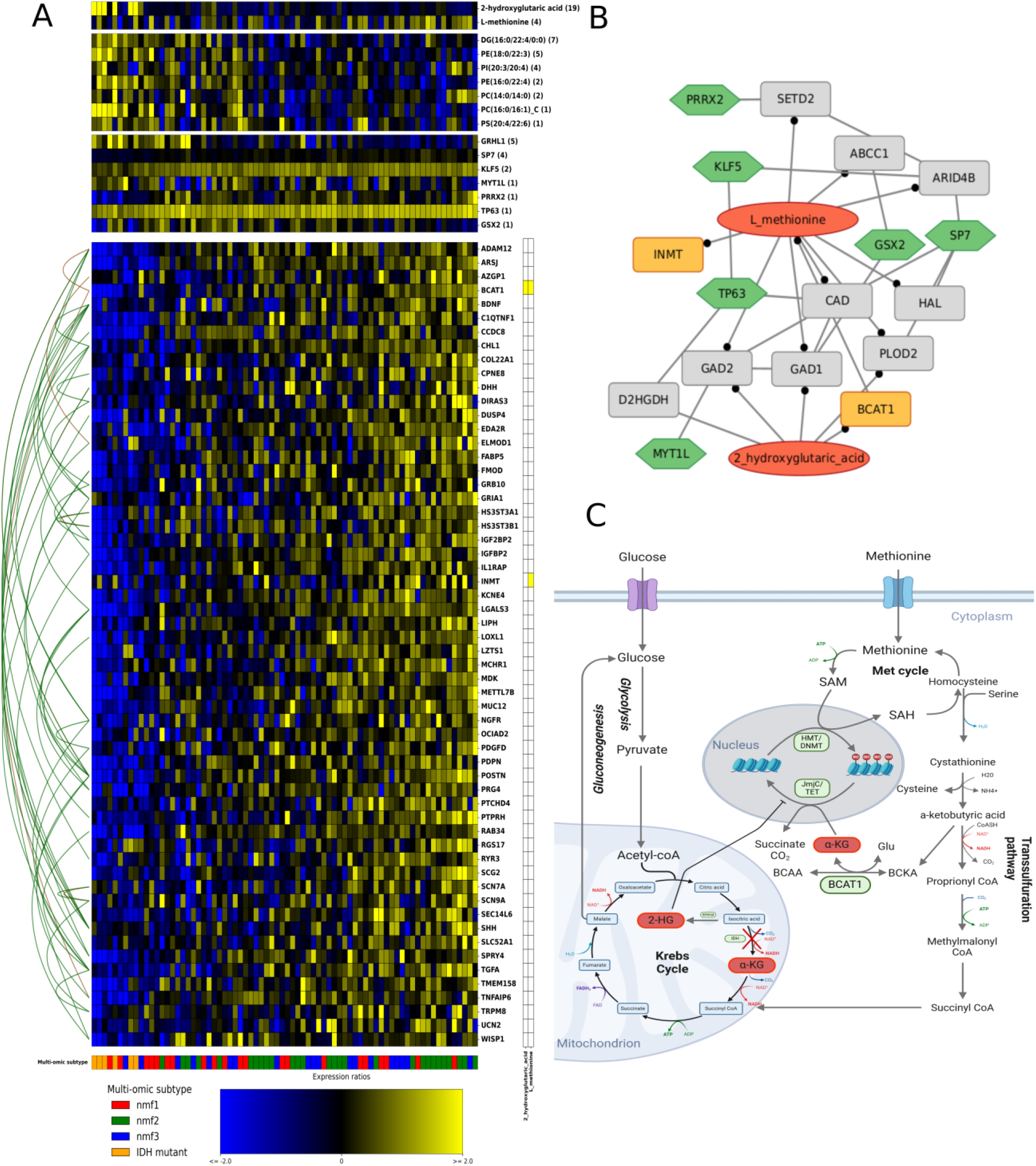
Module 18 connects 2-HG, L-methionine and DNA/histone methylation through BCAT1 in IDH mutant samples. **A.** Module 18 is colored by Z-scored expression (blue: -2; yellow: +2). TF, lipid and metabolite regulators with respective regulator scores are shown above the module. Annotations on the bottom indicate multi-omics subtypes (proneural/nmf1: red; mesenchymal/nmf2: green; classical/nmf3: blue; IDH mutant: orange), yellow boxes on the right represent metabolite-gene interactions that are present in the Lemonite KG. Lines on the left represent physical PPIs (green) or HumanNet functional interactions (brown) between module genes. **B.** Module 18 in the Lemonite KG. Metabolite regulators colored in red, TF regulators in green, module genes in orange. Nodes that connect metabolite regulators to TF regulators through a single PPI in the LemonIte KG colored in grey. Grey boxes represent genes involved in a protein-protein interaction that connects metabolite regulators to TF regulators in the Lemonite KG. Edges ending in a dot represent metabolic pathway interactions, normal lines represent PPIs and ambiguous interactions. **C.** The Lemonite KG provides molecular biological context around module genes and predicted regulators.

TF regulators of module 18 included GRHL1, SP7, KLF5, MYT1L, PRRX2, TP63 and GSX2. Several of these transcription factors have been implicated in cancer-related processes, including cell proliferation, differentiation, stemness, and stress-response pathways across multiple tumor types. Among them, MYT1L has been shown to regulate GBM cell proliferation through repression of YAP1 expression (Melhuish *et al*., 2018), while PRRX2 has been reported to promote glioma stem cell survival by inhibiting ferroptosis through upregulation of GCH1 (Liu *et al*., 2018). Consistent with these findings, Wang et al. reported increased H2B acetylation in genes associated with ferroptosis in mesenchymal-like GBMs (L.-B. Wang *et al*., 2021), while our analysis identified significant upregulation of module 18, regulated by PRRX2, in mesenchymal- like samples. Notably, ferroptosis activity depends not only on iron metabolism but also on mitochondrial function, lipid peroxidation, and glutathione metabolism, processes closely linked to aberrant TCA cycle activity and IDH mutation status.

Lemonite’s knowledge graph further contextualized these findings by connecting module genes, transcription factors, and metabolites into interpretable molecular networks. PRRX2 connected to L-methionine through a single PPI intermediary, SETD2, a histone methyltransferase responsible for H3K36 trimethylation (Goldstein *et al*., 2020). Similar single-edge connections linked GRHL1, KLF5, and MYT1L to the metabolite regulators 2-HG and L-methionine, illustrating how integration of multimodal data uncovers compact gene–metabolite regulatory relationships. Metabolic pathways involving both 2-HG and L-methionine were also highly interconnected (Fig. 5C). Overall, these results demonstrate how Lemonite combines data-driven module detection and regulator assignment with knowledge-graph contextualization to reveal biologically meaningful metabolite–gene associations and generate testable hypotheses on the interplay between gene regulation and metabolism.

### Lemonite population-level gene modules facilitate cell-type specific exploration of metabolite-gene interactions in GBM

Although Lemonite primarily facilitates integration of bulk transcriptomics and metabolomics data, also other bulk omics data types (proteomics, phosphoproteomics, copy number variation, microbiomics, epigenomics…) and even single cell transcriptomics can be integrated. To obtain a cell-type specific view on the metabolite-TF module network, we mapped the gene modules onto single cell transcriptomics data. We therefore preprocessed snRNA-seq data from 6 matched samples (two from each subtype excluding IDH-mutant, Methods). This resulted in a dataset containing 45 144 cells, which were clustered and annotated using GBmap as a reference atlas (Ruiz-Moreno *et al*., 2025) (Methods). Projecting the population-level gene modules identified in bulk transcriptomics data onto single cell transcriptomics data uncovered cell-type specific module expression patterns (Fig. 4C). Modules 5, 13, 23, 36 and 50 were mainly expressed in TAMs and monocytes, coinciding with these modules being upregulated in mesenchymal-like samples, and having immune system functional enrichment results (GSEA) (Fig. 4B-C, Supplementary Fig. 4,12-15, Supplementary Data 1). Modules 26, 9, 30, 2, 12, 62, 17, and 53 were highly expressed across OPCs as well as NPC-like and OPC-like cells (Supplementary Fig. 16-22). In contrast, module 4, 6 and 8 expression was largely restricted to OPCs. This cell-type specificity was consistent with the bulk-level expression patterns which showed high expression in pro-neural like samples (Fig. 4B–C, Supplementary Text). Notably, module 6 was upregulated in nmf1 samples and was predicted to be regulated by MYT1L and EMX1, two transcription factors involved in neuronal development (Kobeissy *et al*., 2016; Mall *et al*., 2017). Metabolite regulators included heptadecanoic acid, 2-HG and creatinine, the latter a breakdown product of creatine which can be produced by oligodendrocytes to support neighboring neural cells (Flögel *et al*., 2025). Module 10 was highly expressed in mesenchymal-like samples at bulk level, specifically expressed in CD4/CD8 T cells at single cell level, and enriched for ‘T cell activation’ (p-adj < 0.05, GSEA, Supplementary Data 1). Thus, even in the absence of coupled single-cell transcriptomics- metabolomics data, Lemonite results obtained using bulk datasets can be integrated with single- cell expression data to investigate cell-type specific metabolite-TF-module interactions. All aforementioned modules have been described in more detail in Supplementary text.

### Lemonite identifies metabolite-gene regulatory interactions in an IBD multi-omics cohort

We next applied Lemonite to a coupled dataset of 75 UC and control patient samples to integrate stool metabolomics with colon tissue transcriptomics data. Lemonite identified 97 coexpression modules, and 86 were retained after filtering out modules with low functional coherence, after which regulators were selected as described in the Methods section (Supplementary Fig. 2B). Together, the integrated metabolite-TF-module network comprised 3 543 genes, 237 TF regulators and 199 metabolite regulators (Supplementary Data 2). Highly connected TF regulators included SP140, RFX6, SPI1, STAT3 and JPH2. SP140 is a macrophage-intrinsic epigenetic reader that protects against IBD by enabling antibacterial defense programs (Fraschilla *et al*., 2022). STAT3 is a major immune signaling hub that drives mucosal inflammation and promotes intestinal barrier integrity (Pang *et al*., 2021). *SPI1* encodes the PU.1 protein, which is involved in macrophage function (Y. Yu *et al*., 2025), while RFX6 is essential for the development and maintenance of enteroendocrine cells in the gut (Sanchez *et al*., 2024). JPH2 is primarily known as a member of the junctional membrane complex in heart muscle cells, and has not been implicated in IBD (Garbino and Wehrens, 2010). Highly connected metabolite regulators comprised several plasmalogens, carnitine species, adrenate, gabapentin and putrescine (Fig. 6, Table 3).

**Figure 6.**
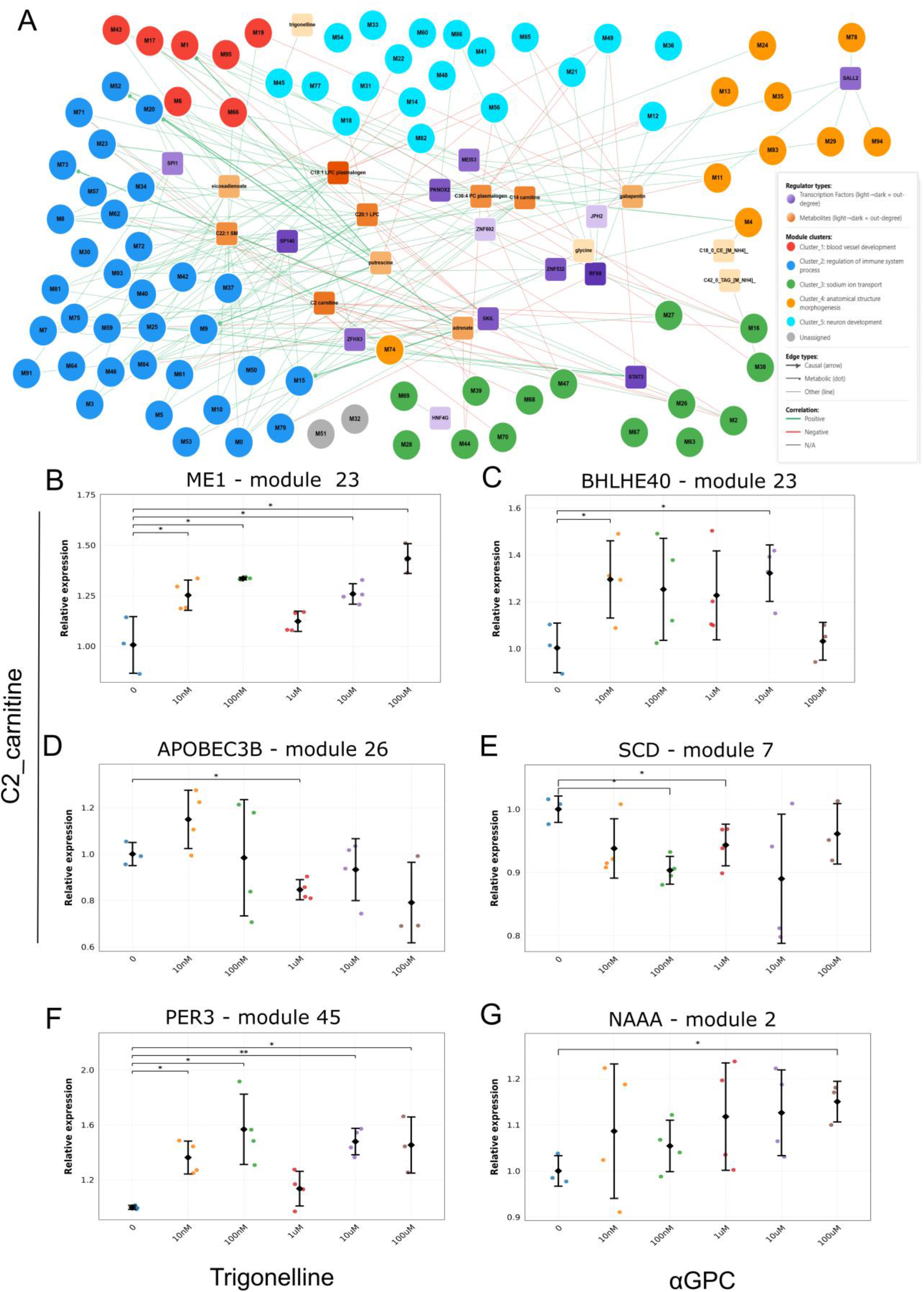
Lemonite infers regulatory metabolite-gene module interactions in an IBD cohort. **A.** Lemonite IBD network containing 46 gene modules and most important metabolite, lipid and TF regulators (Methods). Gene modules are colored by and labeled GO enrichment group (Methods). Edges end in a dot or arrow if the regulator-module connection captures known metabolic pathway or causal interactions resp. Edge color represents the sign of correlation between regulator expression and module expression. The complete network can be explored and searched interactively at www.lemonite.ugent.be.. **B-G.** Experimental validation of a prioritized set of predicted metabolite-gene regulatory interactions. **B.** Relative gene expression of ME1 24 h after treatment with C2_carnitine. **C.** Relative gene expression of BHLHE40 24 hours after treatment with C2_carnitine. **D.** Relative gene expression of APOBEC3B 24 hours after stimulation with C2_carnitine. **E.** Relative gene expression of SCD 24h after treatment with C2 carnitine. **F.** Relative gene expression of PER3 24 hours after stimulation with Trigonelline. **G.** Relative gene expression of NAAA 24h after treatment with α-glycerophosphocholine. Statistical significance was assessed using t-test compared to baseline (concentration 0) and p-values corrected using Benjamini & Hochberg procedure (p-adj < 0.1).

**Table 3.** Hub regulators in the integrated metabolite-TF-module IBD network. Top 5 regulators for TFs, metabolites, ranked by ‘Combined score’. Combined score represents the sum of all regulator-module scores for each regulator, ‘# Target modules’ the number of modules each regulators targets, and ‘# Target genes’ the total combined number of genes across all modules targeted by a given regulator.

| **Regulator** | **Combined score** | **# Target modules** | **# Target genes** |
|---|---|---|---|
| **C18:1 LPC Plasmalogen** | 47.14 | 20 | 1204 |
| **C2 carnitine** | 40.29 | 16 | 766 |
| **C22:1 SM** | 39.10 | 13 | 911 |
| **C20:1 LPC** | 35.77 | 15 | 860 |
| **C14 carnitine** | 35.74 | 14 | 844 |
| **Adrenate** | 20.90 | 9 | 511 |
| **Gabapentin** | 18.85 | 9 | 343 |
| **Putrescine** | 19.94 | 8 | 344 |
| **5-aminovulenic acid** | 16.38 | 8 | 416 |
| **Quinine** | 13.08 | 4 | 89 |
| **SP140** | 42.82 | 6 | 609 |
| **RFX6** | 28.63 | 7 | 286 |
| **SPI1** | 20.81 | 4 | 352 |
| **STAT3** | 20.26 | 7 | 470 |
| **JPH2** | 19.70 | 2 | 76 |

Plasmalogens are abundant in intestinal epithelium, protect against oxidative stress, and are decreased in IBD although their exact role remains to be elucidated (Longo *et al*., 2020). Carnitine metabolism is altered in IBD, with changes reported across multiple acylcarnitine species, including short-chain species such as acetylcarnitine (C2 carnitine) and longer-chain acylcarnitines (C14 carnitine) reflecting disrupted mitochondrial fatty acid oxidation (Lloyd-Price *et al*., 2019; Smith *et al*., 2021). Adrenate (22:4 n-6), a long-chain omega-6 polyunsaturated fatty acid derived from arachidonic acid, is incorporated into membrane phospholipids and contributes to the pool of substrates available for bioactive lipid mediator synthesis. Altered PUFA metabolism in IBD may therefore reflect disrupted lipid homeostasis and inflammatory signaling (Di’Narzo *et al*., 2022). Altered adrenate metabolism in IBD may reflect disrupted PUFA homeostasis and inflammatory lipid signaling. Gabapentin has been reported to exert anti-inflammatory effects in experimental colitis models through modulation of PPARy/Nf-kB signaling and reducing inflammatory cytokine production (de Brito *et al*., 2020). Putrescine is produced by both the host and gut microbiota and may have a dual role in IBD, promoting epithelial repair and barrier integrity under homeostatic conditions while exacerbating inflammation under certain pathological conditions (Grosheva *et al*., 2020; Nakamura *et al*., 2021). Modules were further prioritized based on differential expression (63 out of 86 with p-adj < 0.05, Mann-Whitney U test) or PPI enrichment among module genes (33 out of 86 with p-adj < 0.05, hypergeometrical test).

Module 49 was the top DE module (p-adj = 5.96e-10, Mann-Whitney U test, Supplementary Data 2) and was downregulated in UC patients compared to controls (Supplementary Fig 5A, Supplementary Data 2). Genes in this module were enriched for GO terms related to ion transport (Supplementary Data 2), which is well known to be deregulated in IBD patients (Magalhães *et al*., 2016; Prasad and Visweswariah, 2021). Interestingly, the top 2 metabolite regulators for module 49 were both plasmalogens, which are integral components of cellular membranes and thus intrinsically associated to ion transport across membranes (Messias *et al*., 2018). TF regulators included PURA, RFX6, TEAD3, ZNF443 and STAT3, several of which have previously described roles in IBD. Among these, RFX6 is a key regulator of intestinal enteroendocrine cell differentiation and gut hormone secretion (Piccand *et al*., 2019), TEAD3 belongs to the TEAD family that mediates Hippo-YAP signaling, a pathway increasingly implicated in intestinal epithelial regeneration and IBD pathogenesis (Xie *et al*., 2021), and STAT3 has a well-established, if context-dependent, role in both the innate and adaptive immune dysregulation underlying IBD (see above).

Lemonite is a data-driven data-integration method, which prioritizes metabolite-gene predictions, and can be experimentally validated both *in silico* (using the Lemonite KG e.g.) and experimentally. Several examples of how predicted interactions between metabolite regulators and module genes can be validated *in silico* were discussed above, here we explored prioritization of predictions for experimental validation *in vitro*. We selected three metabolites, on the one hand based on their out-degree in the metabolite-TF-module network, with C2 carnitine and α-glycerophospholine (α- GPC) both being highly connected (Fig. 6A), and/or on the other hand based on a high regulator- module score (Fig. 6A, trigonelline-module 45 was the second highest scoring interaction). We next selected target modules for these regulators through manual prioritization; module 23 was the 3d ranking DE module among target modules for these regulators (p-adj = 4.42e-7, Mann- Whitney U test, 5^th^ DE rank overall, Supplementary Fig 6), module 7 was significantly enriched for positive regulation of cytokine production (p-adj < 0.05, GSEA, Supplementary Fig. 7A-B), module 2 was significantly enriched for known metabolite-gene interactions (p-adj=2.68e-14, hypergeometric test, Supplementary Fig. 8A-C), and module 45 – trigonelline was the 2^nd^ highest scoring regulator-module pair in the network (Supplementary Fig. 9). Next, we identified a number of genes in these modules that were adequately expressed in the colonic epithelial cell line HT29, according to the DepMap portal (Arafeh *et al*., 2025), and that had a biological relation to IBD, inflammation or the respective metabolite regulator (Supplementary Table 3).

HT29 cells were stimulated with a dose range of respective metabolites, and gene expression was evaluated 24 hours after treatment through quantitative PCR (qPCR). Treatment of HT29 cells with C2 carnitine, trigonelline or α-GPC did not induce toxicity at concentrations ranging between 10nM and 100µM (Supplementary Fig. 10). C2 carnitine perturbation caused a significant upregulation of *ME1* (module 23) gene expression after treatment in all doses tested, without dose–response relationship (Fig. 6B). *BHLHE40*, which was also assigned to Lemonite IBD module 23, was upregulated after treatment with carnitine. Because of the large variation, only 10nM and 10µM conditions reached significance (Fig 6C). For module 26, *APOBEC3B* was downregulated after treatment with 1µM C2 carnitine (Fig 6D). C2 carnitine also led to a downregulation of *SCD* (module 7) expression at concentrations of 10nM and 1µM (Fig. 6E). Trigonelline was only assigned to a single module as a regulator, module 45, and had the second highest regulator-module score overall. Treatment of HT29 cells with trigonelline caused an upregulation of *PER3* expression at 10nM, 100nM, 10 µM and 100µM (Fig. 6F). Finally, α-GPC caused an upregulation of *NAAA* (module 2) 24h after treatment with the highest dose (Fig. 6G). For 21 out of 27 selected metabolite-gene pairs, we could not observe a significant regulatory effect (Supplementary Table 4). It must be noted that none of these metabolite-gene pairs were associated with known interactions in the Lemonite KG and were detected in a purely data-driven manner by Lemonite, thus showing that Lemonite identified novel metabolite-gene interactions that are not known a priori.

## Discussion

Metabolites (including lipids), along with nucleic acids and proteins, play pivotal roles within complex, interdependent molecular and cellular networks. As such, phenotypical variation in complex living systems does not arise from a single omics layer acting in isolation but from the dynamic interplay among these interconnected components (Klein et al., 2021; Ball, 2025). Metabolic reprogramming in cancer, for instance, is not limited to changes in metabolite levels: these in turn actively affect chromatin accessibility and gene expression by providing substrates and cofactors for chromatin-modifying enzymes, such that metabolites act as true molecular ’regulators’ (Sdelci *et al*., 2019). Yet current multi-omics integration methods largely fail to capture this regulatory role of metabolites directly. Lemonite addresses this gap by 1) building an extensive protein/metabolite-gene knowledgebase, the Lemonite KG, which unifies information across databases under a single unified metabolite identifier (HMDB ID), 2) including all metabolites — also un- or ambiguously annotated ones — in the data integration process, 3) requiring no prior differential analysis or feature selection before integration, 4) performing integration in a fully data-driven manner, unconstrained by known metabolite-gene interactions or pathways, and 5) producing an interpretable metabolite-TF-module network that is further contextualized by the Lemonite KG. Together, these design choices distinguish Lemonite from existing approaches, which typically require choosing between unbiased, data-driven discovery and mechanistic interpretability (see below).

Multi-omics data integration methods traditionally fall into one of two categories. Data-driven methods such as MixOmics (Rohart *et al*., 2017), MOFA+ (Argelaguet *et al*., 2020), NMF (Chalise and Fridley, 2017; L.-B. Wang *et al*., 2021) or MOMA (Moon and Lee, 2022) identify hidden underlying patterns in the data and allow unbiased exploration across a broad range of omics, but suffer from low interpretability and are sensitive to noise (Cavill *et al*., 2016; Vandemoortele and Vermeirssen, 2023). Prior-knowledge-based methods such as COSMOS(+) or joint pathway analysis are more easily interpretable, but restricted to known pathways and interactions (Dugourd, 2021; Pang *et al*., 2024), typically require a pre-specified set of ’relevant features’ (e.g. from differential abundance analysis), and rely on databases that use inconsistent metabolite identifiers, complicating cross-database integration (Kanehisa and Goto, 2000; Orchard *et al*., 2014; Szklarczyk *et al*., 2016; Gillespie *et al*., 2022; Berg *et al*., 2023; Braisted *et al*., 2023; The UniProt Consortium, 2023). This trade-off is inherent to metabolomics itself: unlike sequencing- based omics, metabolomics relies on chromatography-mass spectrometry outputs indirectly matched to incomplete reference libraries, leaving a substantial fraction of features only partially identified (Gauglitz *et al*., 2022) including in our data, C18:1 LPC plasmalogen, a top regulator in IBD module 49 whose exact structure remains unknown. Excluding such ambiguously annotated features, as prior-knowledge methods require, therefore risks discarding genuine biological signal.

Applied to our GBM and IBD cohorts, DIABLO and MOFA+ recovered biologically relevant metabolites and genes, including 2-HG in GBM and C18:1 LPC plasmalogen in IBD, alongside consistent transcriptomic features such as *FOLR2* and *FBXO17* (see above). However, neither method could resolve how these molecules relate mechanistically to one another or to disease- relevant gene programs. COSMOS+, in turn, was limited by the number of metabolites that could be mapped onto its meta-network after MOFA+ weight-based filtering (9/93 in factor 1, 1/43 in factor 2, 2/6 in factor 3, 0/2 in factor 4 and 3/40 in factor 6 for GBM), and the resulting networks contained no metabolites at all. Lemonite, by contrast, directly linked metabolite and TF regulators to gene modules: it implicated myo-inositol, phosphatidylcholines and IRF6 in EMT and mesenchymal-like immune programs in GBM (Brosch *et al*., 2022; Kim *et al*., 2024). This extends the original characterization of the GBM mesenchymal subtype, which reported depleted phosphatidylcholines as a subtype-associated lipid signature but did not link this lipid class to a transcriptional driver of the EMT program (L.-B. Wang *et al*., 2021). Similarly in IBD, Lemonite connected plasmalogens to the expression of an epithelial ion-transport gene module via TF regulators including STAT3 and RFX6. The original study identified only three individual host genes (ileal *GIP*, *NXPE4*, and *ANXA10*) as nodes in its combined multi-omic network, with no TF regulators or mechanistic model connecting them to microbial or metabolite features (Lloyd-Price *et al*., 2019). Lemonite’s TF-resolved, module-level metabolite-gene assignment therefore adds a layer of host-side mechanistic resolution absent from the original study. For metabolite-gene pairs consistently prioritized across methods, Lemonite thus contributes mechanistic hypothesis- generation that purely data-driven or overly filtered prior-knowledge approaches, and the original disease-specific multi-omics studies themselves, did not provide.

Several limitations warrant discussion. First, the high degree of incompleteness and complementarity across metabolite-gene databases, likely in part driven by the understudied nature of these interactions, argues against restricting integration to prior knowledge from the outset. None of the metabolite-gene pairs selected for *in vitro* validation, for instance, were annotated as ’interacting’ in the Lemonite KG itself, which is why we did not use the KG as ground truth for precision/recall benchmarking as is traditional for GRN inference methods (Pratapa et al., 2020; Badia-i-Mompel, 2024; Nguyen et al., 2024). Second, experimental validation was performed in a single cell line (HT29), and given that Lemonite modules display cell-type-specific expression patterns (Fig. 4C), this design likely underestimates true positives as well as producing false positives, and results should be interpreted with this caveat in mind rather than as a definitive positive/negative rate. Third, Lemonite’s current regulator assignment integrates omics pairwise, i.e. TFs and metabolites independently from each other, which may miss higher-order dependencies detectable only through cooperative modeling. Finally, like all data-driven approaches, Lemonite’s statistical power depends on cohort size; the moderate size of the GBM and IBD cohorts analyzed here constrains the robustness of module-level and regulator-level estimates, and larger cohorts will be needed to validate and extend these findings.

Technological advancements have recently enabled mass-spectrometry-based single-cell and spatial metabolomics and proteomics (Alexandrov, Saez-Rodriguez and Saka, 2023), which, combined with single-cell transcriptomics and chromatin accessibility data, promise unprecedented resolution for studying cellular development and behavior. Integration methods tailored to these single-cell and spatial multi-omics modalities remain largely limited to dimensionality reduction- or correlation-based approaches, in contrast to single-cell transcriptomics/chromatin accessibility integration, which already leverages TF binding motif information in open chromatin regions (Badia-i-Mompel *et al*., 2023; Loers and Vermeirssen, 2024). Growing data availability also opens the door to regularized machine and deep learning models that incorporate, without being restricted to, prior knowledge such as metabolic pathway information or known allosteric interactions (Doncevic and Herrmann, 2023; Lobentanzer *et al*., 2024; Tejada-Lapuerta *et al*., 2025). Metabolic modeling based on single-cell transcriptomics data offers an exciting complementary avenue, but current approaches do not directly model metabolomics data nor identify regulatory metabolites and their target genes (Alghamdi *et al*., 2021; Hrovatin, Fischer and Theis, 2021; Wagner *et al*., 2021; Zheng, 2022; B. Wang *et al*., 2025). We therefore foresee substantial potential for extending Lemonite towards single-cell and/or spatial metabolomics integration with other data modalities.

## Materials & methods

### Construction of the Lemonite knowledge graph

The knowledge graph was constructed using the Python (3.11.11) programming language, and scripts are available at www.github.com/CBIGR/Lemonite/build_PKN. A dedicated pipeline has been built that allows to regularly update the KG. We downloaded the human metabolome database (HMDB, https://www.hmdb.ca/) (Wishart *et al*., 2022) on October 7^th^ 2024 and parsed the xml file to a dataframe containing all metabolites in the HMDB with their respective HMDB ID ChEBI ID, KEGG ID, PubChem ID, IUPAC name, SMILES, InChIKey and PDB ID, next to ‘subclass’, ‘superclass’ and ‘kingdom’ annotations. Using these identifiers, we queried IntAct, UniProt, chEMBL, LINCS, BIOGRID and STITCHdb, Human1GEM and MetalinksDB for interacting genes/proteins (see below for details per database).

**STITCHdb** (Szklarczyk *et al*., 2016). We downloaded all human chemical-protein interactions from the STITCH database on October 9^th^ 2024 from the web portal (http://stitch.embl.de/cgi/download.pl?UserId=hQ516L8aOiqg&sessionId=gW2dEJfn9Ggp&species_text=Homo+sapiens). Ensemble protein IDs were mapped to gene symbols using the mygene Python package and interactions involving features with an invalid gene symbol were removed. Next, we retrieved all interactions involving HMDB metabolites through their PubChem IDs.

**IntAct** (Orchard *et al*., 2014). The IntAct database was queried for chemical-protein interactions using ChEBI IDs on 13/10/2025 through its Python API without specifying a minimum score. Only interactions containing ‘9606’ (human taxonomic ID) in columns ‘taxIdAStyled’ or ‘taxIdBStyled’ were retained.

**ChEMBL** (Zdrazil *et al*., 2024). SMILES codes for all HMDB metabolites with a valid SMILES in the database were first converted to canonical SMILES using the RDKit Python module. The chEMBL activity database was next queried using canonical SMILES IDs through its Python package ‘chembl_webresource_client’. The chEMBL activity database was first filtered for interactions involving human proteins by dropping all interactions not annotated with ‘target_tax_id’ = 9606 and all interactions in which ‘activity_comment’ was not either ‘active’ or ‘substrate’. chEMBL interactions were therefore considered to be ‘causal’ interactions in the Lemonite knowledge graph.

**UniProt** (The UniProt Consortium, 2023). The UniProt database was queried using InchiKeys for chemical-protein interactions involving human proteins through its Python REST API. The ’reviewed’ parameter was set to False.

**LINCS** (Keenan *et al*., 2018). The LINCS small molecule suite (Moret *et al*., 2019) was downloaded from the web portal (<u>Small Molecule Suite</u>) on 15/10/2025. This included multiple files: lsp_compound_dictionary.csv (ID mapping for chemical compounds), lsp_target_dictionary.csv (ID mapping for targets) and lsp_biochem_agg.csv. Interactions between small molecules and gene were then retrieved from lsp_biochem_agg.csv using metabolite chEMBL IDs, an IC50 threshold of 10 µM and after filtering out all non-human targets.

**BIOGRID** (Oughtred *et al*., 2021). The BioGRID chemicals database was downloaded on 15/10/2024 from the web portal (https://downloads.thebiogrid.org/File/BioGRID/Release-Archive/BIOGRID-4.4.238/BIOGRID-CHEMICALS-4.4.238.chemtab.zip ). Interactions were parsed by querying InChiKeys.

**Human1-GEM** (Robinson *et al*., 2020, p. 1). The Human1-GEM is an extensively curated atlas of human metabolic pathways comprising both the Recon and HMR metabolic reconstruction lineages. The human1 metabolic model was downloaded on 06/06/23 from its Zenodo archive (https://zenodo.org/records/7912808). The model was first parsed to a Python networkx graph by adding all unique metabolites in the model as node to an empty graph, after which edges were drawn between nodes representing metabolites that are annotated as ‘products’ or ‘reactants’ within a single metabolic reaction. Nodes with too many connections were removed from the model in order to avoid the emergence of non-meaningful interactions (Supplementary Table 5), resulting in L-carnitine being the most highly connected node with 254 remaining interactions. Each edge was assigned the reactions name as attribute. Next, all reactions without a valid ‘Gene- reaction association’ were dropped from the model and a dictionary was created to map each individual metabolic reaction to a set of genes facilitating the metabolic reaction. The resulting metabolic network with metabolites as nodes and metabolic reactions as edges, with each reaction/edge coupled to a set of genes using the model’s gene protein reactions rules, was then used to identify metabolite-gene interactions. Specifically, a gene was considered to interact with a given metabolite if it encoded an enzyme participating in a reaction that was reachable from that metabolite within a maximum path length of two reaction steps in the metabolic network. Subcellular localization and compartmentalization of metabolic reactions were not taken into account. Human1 metabolite IDs were then mapped onto ChEBI IDs using the ‘metabolites.txt’ file provided with Human1-GEM.

**MetalinksDB** (Farr *et al*., 2024). MetalinksDB was accessed on 16/10/24 through the ‘resource.get_metalinks (source = [’CellPhoneDB’, ’Cellinker’, ’scConnect’, ’recon’, ’hmr’, ’rhea’, ’hmdb’]) function from the LIANA+ package implemented in Python (Dimitrov *et al*., 2024). Metabolite HMDB IDs were coupled to interacting genes through the ‘hmdb’ and ‘gene_symbol’ columns present in the database.

**Protein-protein interactions.** We first compiled a list of all genes involved in metabolite-gene interactions. Next, STRING, BIOGRID and HuRI (Luck *et al*., 2020) were queried for protein-protein interactions involving at least one protein of which the corresponding gene was present in our list of genes involved in metabolite-gene interactions. The STRING database (version 11.5) was first queried through its Python API with method ‘get_string_ids’ on 17/10/2025 in order to map gene symbols to string IDs. Next, the ‘interaction_partners’ method was used to return all interacting genes for each string ID in the dataset. Second, the full BIOGRID database was downloaded from the web portal on 02/10/2024 and all interactions involving at least one non-human protein were dropped. Genes involved in PPIs were then extracted using the ‘Official Symbol Interactor A’ and ‘Official Symbol Interactor B’ columns, and only interactions involving at least one gene of the metabolite-gene list were retained. Finally, the human reference interactome (HuRI) was downloaded on 03/10/2024 from the web portal (http://www.interactome-atlas.org/data/HuRI.tsv).

Interactions not involving at least one gene involved in metabolite-gene interactions were dropped from the database. Protein IDs and Ensemble gene IDs were mapped to gene symbols using BiomaRt (version January 2024).

**HumanNet** (Kim *et al*., 2022). The HumanNet-FN network (without co-citation interactions) was downloaded from the HumanNet web portal on 28/04/26 (https://www.inetbio.org/humannetv3/download.php). This network contains both physical PPIs and functional interactions, and were annotated as such before merging them with the rest of the PPIs.

**The Lemonite KG.** Finally, the metabolite-gene network and protein-protein interaction networks were combined into a single integrated metabolite-gene PKN network in simple interaction file (SIF) format. This file format contains one interaction per line, with HMDB IDs being used as a uniform metabolite identifier and gene represented by gene symbols, accompanied by the source from which the interaction was retrieved. This allows to keep track of the source of each individual metabolite-gene or protein-protein interaction in the integrated PKN, and also provide a hyperlink to the original database for each interaction in the metabolite-gene interaction in the Lemonite knowledge graph. This resource can be accessed and searched online: www.lemonite.ugent.be.

## Data preprocessing

### GBM dataset

Bulk total RNA sequencing, untargeted metabolomics (GC-MS) and lipidomics data (LC-MS/MS) were downloaded as .xlsx files from the Supplementary information in the original publication , gene symbols’ names were manually checked and files were saved as .txt for further use. Control GTEx samples were removed and only samples for which RNA sequencing, metabolomics and lipidomics data were available were kept for further analysis. Also two matching single-nuclei RNAseq samples were downloaded (https://portal.gdc.cancer.gov/projects/CPTAC-3) for the following multi-omics subtypes: proneural-like (nmf1): C3N-03188 & C3N-01334; mesenchymal-like (nmf2): C3N-02190 & C3N- 02784; classical-like (nmf3): C3L-03986 & C3L-03405 .

### IBD dataset

Publicly available bulk total RNA sequencing and untargeted metabolomics data (Lloyd-Price *et al*., 2019) were respectively downloaded from NCBI’s Gene Expression Omnibus (GSE111889) and the Metabolomics Workbench (project ID PR000639). Sample IDs from the RNAseq dataset were mapped to patient IDs, and metabolomics samples not corresponding to these patients were removed. Furthermore, since metabolomics data were of longitudinal nature, only the sample taken at the earliest point in time for each patient was retained to match as close as possible the RNAseq sampling time. Metabolomics data from the different platforms (IBD: C18 pos, C18 neg, HILIC pos & HILIC neg; GB: lipidome pos & lipidome neg) was merged into a single dataframe.

### Metabolite annotation

Both datasets provided metabolomics data in which metabolites were identified by their common name. These were mapped to HMDB IDs using the Compound ID Conversion tool from MetaboAnalyst (Pang *et al*., 2024). Entries for which no exact match was found were manually inspected (using the pop-up that MetaboAnalyst provides) in order to maximize the number of metabolites with a valid HMDB ID in the dataset.

### Preprocessing bulk data

Data preprocessing was performed in the R (4.4) programming language. Genes not corresponding to protein coding genes or duplicated gene names were removed from the RNA sequencing datasets, as well as genes with count <= 10 in at least 3 samples (ensemble database January 2024). Highly variable genes (HVGs) were selected based on a variability cutoff (> 0.7 in GBM and > 0.35 in IBD) after visual identification of the knee point in variability distribution, resulting in 5817 retained genes for GBM and 4429 for IBD, including 781 and 695 TFs respectively from the Lovering TF list (Lovering *et al*., 2021). Count data from the IBD dataset were normalized and log transformed using default DESeq2 (1.46.0) (Love, Huber and Anders, 2014). For metabolomics and lipidomics abundancy data, NA entries were replaced by zero, after which abundances were log transformed (after adding a pseudo-abundance of 1) and scaled by the *pareto_scale()* function in the IMIFA R library (2.2.0). Transcriptomics and metabolomics data were then vertically concatenated to create a single multi-omics dataframe.

### TFA inference

TFAs were inferred from log-normalized transcriptomics data using DecouplR’s (2.12.0) consensus function with default arguments and the CollecTri TF-target gene network (Badia-i-Mompel *et al*., 2022; Müller-Dott *et al*., 2023). TFs that were present in the CollecTri network but previously removed from the dataset due to HVG selection were re-introduced into the preprocessed gene expression dataframe with their original log-normalized, scaled gene expression values. Next, for all TFs for which TFA was calculated, gene expression values were replaced by TFA scores.

### MixOmics

(6.30.0) (Rohart *et al*., 2017). Transcriptomics and metabolomics data (and lipidomics data in case of GBM) were preprocessed as described above. A DIABLO (Data Integration Analysis for Biomarker Discovery using Latent Variable Approaches for Omics Studies) (Singh *et al*., 2019) model was then constructed using MixOmics *block.splsda* function (ncomp = 5, default design with value 0.1), after which the ideal number of components was determined using a weighted vote that considers *‘Overall.BER’* and *‘centroid.dist’* after running the *perf* function with 10 fold cross-validation and 10 repeats. The optimal number of features in each omics was selected using the *tune.block.splsda* and these were used to construct a final DIABLO model. Figures were created with the *plotDIABLO, plotIndiv* and *plotLoadings* functions. Gene set enrichment analyses were performed by first ranking all features per component by their feature weights and then applying *ClusterProfiler* (nPermSimple = 10000, minGSSize = 3, maxGSSize = 800, pvaueCutoff = 0.05 & pAdjustMethod = ‘BH’).

#### MOFA

(1.16.0) (Argelaguet *et al*., 2018). Transcriptomics and metabolomics data were preprocessed as described above. Datasets were summarized as *MultiAssayExperiment (1.42.0)* and converted to a MOFA object. A MOFA model was then trained using default options (convergence_mode = slow) after which the variance explained by each factor was visualized using *plot_variance_explained().* Sample loadings on each factor were investigated using the *plot_factor* in combination with a Kruskal-Wallis test to identify factors in which sample groups displayed significantly different sample loadings (p-adj < 0.05). Feature weights of factors were extracted using the *get_geneweights()* from the *MOFAcellulaR* R package, after which the top 10000 genes with the highest absolute weights for each factor were selected, ordered, and used for gene set enrichment analysis. *Gene set enrichment analyses were performed by first ranking all features per component by their feature weights and then applying ClusterProfiler* (nPermSimple = 10000, minGSSize = 3, maxGSSize = 800, pvaueCutoff = 0.05 & pAdjustMethod = ‘BH’).

#### COSMOS+

(CosmosR 1.14.0) (Dugourd *et al*., 2024). The tutorial provided by the authors of the MOFA2COSMOS method was followed (<u>MOFA to COSMOS tutorial • cosmosR</u>). Feature weights were extracted from the MOFA model using the *get_weights()* function. The distribution of weights for metabolite features was visualized and a cutoff was set to abs(weight) > 0.2 (GB). Metabolite names were mapped to their corresponding HMDB ID, after which they were prepared for further COSMOS analysis using the *prepare_metab_inputs()* function from the COSMOS package (Dugourd, 2021). The overlap between remaining metabolites and metabolites in the COSMOS meta-network was then evaluated.

## Lemonite pipeline

### Clustering

Genes were clustered into coexpression modules using LemonTree’s Ganesh function (Bonnet, Calzone and Michoel, 2015). Briefly, 100 independent G model-based Gibbs samplers were applied to simultaneously infer coexpression modules and condition clusters within each module using the preprocessed transcriptomics data. Next, genes that systematically clustered together across multiple Ganesh runs were grouped in consensus clusters through spectral edge clustering, which identifies densely connected sets of nodes in a weighted graph, in which weights represent the number of times individual nodes were assigned to the same cluster across multiple Ganesh runs.

### Regulator assignment

Regulators were assigned to modules in separate runs for TFs, lipids and metabolites. A list of putative TF, lipid and metabolite regulators were provided to the Lemonite pipeline These lists encompassed all TFs from the Lovering TF list (Lovering *et al*., 2021) that were expressed in the dataset, all lipids, and all metabolites expressed in the dataset. Lemonite infers an ensemble of regulatory programs for a set of coexpression modules and calculates a consensus regulator-to-module score. Each regulatory program is represented as a decision tree in which the expression levels of the regulators represent the internal nodes, and the regulator score considers the number of trees a given regulator is assigned to, with what score, and at which level of the tree. Regulators are also randomly assigned to modules and their scores are computed, which are used to assess significance of the predicted regulators for each module.

### Construction and visualization of an integrated metabolite-TF module GRN and downstream analyses

All downstream analyses were performed in Python. Module coherences were calculated by calculating the average correlation between module genes and the module eigengene, and modules with coherence < 0.5 were removed as these represent low-quality coexpression modules. Next, only the top 2% regulators for which the regulator-to-module score was higher than the maximum random score, were retained. An additional filtering step was applied in which all regulators with a score 10 times smaller than the highest scoring regulator for that module were removed. Regulator scores were than normalized by dividing their score by the total sum of scores of all selected regulators across all modules (per regulator type), and then multiplying the resulting scores by 1000 and rounding them to the nearest integer. Hub regulators were identified by counting the number of modules each regulator was assigned to and calculating the summed score of each regulator across all modules. Differentially expressed modules were identified using a non-parametric Mann-Whitney U test in case of 2 sample groups (IBD), or a Kruskall-Wallis test in case of > 2 sample groups (GB), both with Benjamini & Hochberg correction for multiple testing. Enrichment analysis (overrepresentation analysis) on module genes and combined sets of target genes for each regulator (some regulators regulate > 1 module) was performed using EnrichR against GO Molecular function 2025, GO biological process 2025 and GO Cellular Component 2025 databases. Also, for each module, all genes in the Lemonite network were ranked based on correlation (PCC) between the module eigengene and gene expression across samples, after which this ranked list was used as input for gene set enrichment analysis using *ClusterProfiler* (nPermSimple = 10000, minGSSize = 3, maxGSSize = 800, pvaueCutoff = 0.05 & pAdjustMethod = ‘BH’). For module heatmap visualizations, we adapted, extended and translated (from Java to Python) the existing ModuleViewer software (<u>GitHub -</u> <u>thpar/ModuleViewer: Interactive figure generator for clustered expression data · GitHub</u>), and implemented this within the Lemonite framework Python scripts using matplotlib and seaborn packages. The presence of ‘known interactions’ between metabolite regulators and module genes was assessed by comparing the Lemonite data-driven network to the Lemonite KG. (Kim *et al*., 2022). Module subnetworks were created per module and show TF and metabolite regulators, module genes which have ‘known interactions’ with metabolite regulators in the Lemonite KG, and genes that allow to connect metabolite and TF regulators through a single PPI in the Lemonite KG. On these subnetworks, metabolite-gene interactions were annotated as ‘causal’ (LINCS, chEMBL), ‘other’ (UniProt, BioGRID, STITCHdb, IntAct, MetalinksDB) or ‘Metabolic pathway’ (Human1-GEM). Subnetworks were created using the networkx python package.

### PPI enrichment

PPI data were obtained from the Lemonite prior knowledge network and restricted to interactions for which both genes were present in at least one module. Enrichment was assessed using the hypergeometric distribution in which the background population was defined as all possible gene–gene pairs among genes included in the analysis, and success states corresponding to gene pairs annotated as PPIs in the LemonIte knowledge graph. For each module, the sample consisted of all possible gene–gene pairs within that module, and the observed number of successes corresponded to the number of PPIs detected among its genes. Upper-tail p-values were calculated to test whether modules contained more PPIs than expected by chance given the global PPI density. P-values were corrected using the B&H procedure and modules with p-adj < 0.05 were considered significantly enriched for PPIs. PPI data were plotted next to module figures through lines that connect corresponding gene-gene pairs.

### Module overview

A large module overview file was created that summarizes for each module the set of TF regulators, lipid regulators, metabolite regulators, module genes, DE analysis results, PPI enrichment results and the top 3 enriched pathways per database. Only modules that passed coherence filtering were included, and modules were ordered based on differential expression results (lowest to highest p-value).

### Network visualization

For network visualizations, we kept all module nodes together with regulator nodes that were 1) among the top 10 highly connected regulator nodes per regulator type or 2) among the top 5 regulator-module scores per regulator type. Modules were colored based on megaGO groupings. Briefly, for each module, the top 30 significantly enriched GO biological process pathways were listed and megaGO (Verschaffelt *et al*., 2021) was used to calculate a pairwise module-to-module similarity score. The resulting scores were used to construct a similarity matrix to which hierarchical clustering was applied to group modules into 5 groups. Next, for each cluster of modules, rrgvo was used to reduce the GO categories for these modules into a single class/term, which was then assigned as cluster label to the respective module clusters. Regulator nodes were colored based on the number of modules a given regulator is connected. Interactive network visualizations were created and visualized using pygraphviz.

### Module expression at single-cell level

6 snRNA-seq samples (C3L-03405, C3L-03968, C3N- 01334, C3N-02190, C3N-02784, and C3N-03188) provided by (L.-B. Wang *et al*., 2021) were downloaded from https://portal.gdc.cancer.gov/projects/CPTAC-3 as raw_feature_bc_matrix.

Raw count matrices in mtx format were imported using scanpy and the following filtering measured were applied: minimum 200 and maximum 10 000 expressed genes per cell, minimum 1000 and maximum 10 000 UMI counts per cell, and maximum 10% mitochondrial gene count per cell. Genes expressed in fewer than 3 cells were also removed from the analysis. Samples were then concatenated into a single object, which underwent an additional round of quality control with the same parameters. Cells were annotated using scANVI within scvi-tools and the GBmap_core reference atlas (Gayoso *et al*., 2022; Ruiz-Moreno *et al*., 2025). We first trained a single- cell variational inference (scVI) model on the reference atlas using the following hyperparameters: the batch key specified ’author’ to correct for study-specific batch effects, layer normalization was ’both’, batch normalization was ’disabled’, covariate encoding was ‘enabled’, the dropout rate was set to 0.2, and two hidden layers were used in the network architecture. The trained scVI model was then used to initialize a scANVI model via scvi.model.SCANVI.from_scvi_model(), using the ‘annotation_level_3’ labels as the reference cell types. The scANVI model was trained for 200 epochs with early stopping, with validation metrics evaluated every 25 epochs. The preprocessed query dataset (combined across samples) was prepared for mapping to the reference using scvi.model.SCANVI.prepare_query_anndata(), which harmonized gene sets between the reference and query and retained only the shared genes. A query-specific scANVI model was then initialized and trained for 150 epochs with a learning rate of 1×10⁻⁴ and no weight decay. Cell type predictions were generated using scanvi_query.predict(), and latent embeddings were obtained with scanvi_query.get_latent_representation(). The resulting scANVI-derived predictions were stored in the AnnData object under .obs["predictions_scanvi"].

The merged and annotated data object was further processed by normalizing to 10 000 counts per cell, followed by log-transformation using scanpy’s *normalize_total()* and *log1p()* functions. The top 3000 most highly variable genes were selected in a batch-aware manner using according to the Seurat flavor method implemented in scanpy. The effects of total counts per cell were regressed out using scanpy’s regress_out() function. The expression of bulk Lemonite modules was then projected onto single cells using scanpy’s *score_genes()* function and visualized using scanpy’s *dotplot()* function.

### NextFlow pipeline

Data preprocessing, clustering and regulator assignment using LemonTree software, downstream processing to an integrated metabolite-TF-module network, enrichment analyses and visualizations have been implemented in a NextFlow pipeline. This pipeline was designed for integration of transcriptomics and metabolomics data specifically, but allows integrating other omics data as well. The pipeline is available at www.github.ugent.be/CBIGR/Lemonite/nextflow. GitHub CoPilot was used as a coding assistant during the creation and testing of this pipeline.

### Perturbation experiments

#### Cell culture and perturbation

HT29 cells were thawed, propagated and seeded in 24 well plates at 500 000 cells/well (McCoy mdium). Cells were treated with C2_carnitine (Larodan, 17-0200-7), trigonelline (Sigma-Aldrich, 1686411) or α-glycerophosphocholine (Santa Cruz, sc-301813) at concentrations ranging between 100µM and 10 nM (100 µM, 10 µM, 1 µM, 100 nM, 10 nM and 0). After 24 hours, the perturbed medium was removed and cell were washed with sterile PBS, after which they were lysed with RTL + β-mercapto buffer. Toxicity of the treatments was assessed using an LDH assay in which 1% Triton-X 100 was used as a positive control (Supplementary Fig 9).

#### RNA extraction and qPCR

Total RNA was extracted using BioRad Aurum kits according to the manufacturer’s instruction and cDNA was synthesized using Bioline SensiFAST kit following the manufacturer’s instructions. All primers used for qPCR are listed in Supplementary Table 6. Samples were loaded in duplicate and two housekeeping genes were selected for each metabolite using RefFinder (HPRT and HMBS for alpha-glycerophosphocholine, GAPDH and HMBS for C2_carnitine and trigonelline). Data analysis was done in R. Each treatment was compared to baseline (untreated) using a Student’s T test, after which p-values were adjusted using Benjamini & Hochberg’s procedure. FDR cutoff for significance was set at p-adj < 0.1.

#### qPCR data analysis

qPCR data analysis was performed in the R programming language. For each sample, the geometric mean of the two reference genes was calculated (HPRT and HMBS for aGPC-treated samples, and GAPDH and HMBS for Carbachol- or Trigonelline-treated samples). Technical replicates were averaged pairwise to obtain a single Ct value per biological replicate. Δ Ct values were computed as the absolute difference between the target gene Ct and the geometric mean of the reference Ct values. For each gene, ΔΔCt values were calculated relative to the mean Δ Ct of the untreated control group (dose = 0). Relative gene expression (fold change) was expressed as 2^-ΔΔCt. Normality of the data was assessed using a Shapiro-Wilk test. To test for differential expression, each treatment group was compared with the corresponding untreated baseline (0 concentration) using two-sample, two-sided t-tests. P-values were corrected for multiple testing using the Benjamini–Hochberg procedure. Adjusted p-values were categorized as significant at thresholds of <0.1 (*), <0.01 (), and <0.001 (*). Data are plotted as relative gene expression compared to baseline (untreated).

## Data and code availability

Bulk transcriptomics and metabolomics data analyzed in this study are publicly available from the original publications (Lloyd-Price *et al*., 2019; L.-B. Wang *et al*., 2021). All scripts used for data analysis in this study are available at https://github.com/CBIGR/Lemonite. A website has been built at https://www.lemonite.ugent.be/ on which the Lemonite knowledge graph and GBM and IBD Lemonite networks can be queried.

## Author contributions

B.V. developed Lemonite software, documentation and the interactive website. B.V. build and curated metabolite-gene interaction data for the prior knowledge graph. B.V. conducted data analysis and visualization. B.V., V.V conceived the computational framework and interpreted data analysis. B.V., D.L., R.V., V.V. conceived the in vitro experimental validation. D.L. provided the experimental resources and supervised the in vitro experiments. H.D. guided and together with B.V. performed the in vitro experiments. T.M. offered advice on the computational methodology.

L.V. recommended metabolomics data preprocessing and metabolite naming. B.V., V.V. wrote the manuscript. T.M., L.V. and D.L. reviewed and edited the manuscript. V.V. acquired funding.

V.V. conceived and supervised the study overall.

## Acknowledgements

The authors thank Matthijs Vynck for reviewing the manuscript and associated code. The authors also thank Mirte de Temmerman for preprocessing the single nuclei RNAseq data, and providing tips and feedback during the study. Finally, we thank Ward Valcke for helping build the interactive website.

## Declaration of generative AI and AI-assisted technologies in the manuscript preparation process

During the preparation of this work, the authors used Claude Code for the development of the nextflow pipeline and as a general coding assistant. The author(s) reviewed and edited the output as needed and take full responsibility for the content of the published article.

