## Supplemental text for "Lemonite: identification of regulatory metabolites through data-driven, interpretable integration of transcriptomics and metabolomics data"

**Supplementary file Lemonite manuscript**

### Supplementary tables

**Supplementary Table 1.** List of abbreviated metabolite and lipid names.

| Abbreviated metabolite name | Full metabolite name |
|---|---|
| N-acetyl-L-aspartic… | N-acetyl-L-aspartic acid |
| 2-hydroxyglutaric ac… | 2-hydroxyglutaric acid |
| 2-hydroxybutyric aci… | 2-hydroxybutyric acid |
| DL-3-aminoisobutyric… | DL-3-aminoisobutyric acid |
| PE(P-18:0/20:5);PE(P… | PE(P-18:0/20:5);PE(P-16:0/22:5) |
| D-glucose-6-phosphat… | D-glucose-6-phosphate |
| DL-2-Aminoadipic aci… | DL-2-Aminoadipic acid |
| C18:1 LPC plasmaloge… | C18:1 LPC plasmalogen |
| N2_N2-dimethylguanos… | N2_N2-dimethylguanosine |
| 4-hydroxy-3-methylac… | 4-hydroxy-3- methylacetophenone |

**Supplementary Table 2.** MOFA+ analysis on GBM dataset. Columns 1-3 contain the percentage of total variance explained by the respective MOFA+ factor in transcriptomics, metabolomics and lipidomics data respectively. 'Functional analysis' contains the top 3 GO Biological Process enriched pathways (gene set enrichment analysis, ranked by p-adj). Only factors discussed in the manuscript are shown here, full data can be investigated in Supplementary Data 1.

| Factor | Transcriptomics | Metabolomics | Lipidomics | Functional analysis (GO Biological Process, top 3) |
|---|---|---|---|---|
| Factor1 | 0.006 | 1.848 | 19.748 | Cellular response to copper ion (NES -3.074, p-adj 1.48e-6); Estrogen metabolic process (NES -3.031, p-adj 2.33e-6); Stress response to copper ion (NES -2.933, p-adj 1.64e-5) |
| Factor2 | 2.065 | 1.262 | 16.365 | Neutrophil chemotaxis (NES -2.134, p-adj 4.67e-3); Cellular response to copper ion (NES 2.099, p-adj 1.72e-2); Regulation of cell division (NES -2.072, p-adj 9.76e-3) |
| Factor3 | 11.099 | 0.826 | 4.931 | Defense response to virus (NES 3.116, p-adj 1.72e-8); Cellular response to lipopolysaccharide (NES 2.975, p-adj 1.72e-8); Cellular response to biotic stimulus (NES 2.882, p-adj 1.72e-8) |
| Factor4 | 6.852 | 0.592 | 2.769 | Neurotransmitter receptor transport to plasma membrane (NES -1.969, p-adj 6.56e-2); Neurotransmitter receptor transport to postsynaptic membrane (NES -1.910, p-adj 6.56e-2); Regulation of phospholipid transport (NES 1.865, p-adj 6.70e-2) |
| Factor6 | 3.689 | 1.640 | 2.130 | Estrogen metabolic process (NES 2.377, p-adj 5.81e-4); Epoxygenase P450 pathway (NES 2.144, p-adj 4.59e-3); Intestinal lipid absorption (NES 2.058, p-adj 2.73e-2) |

**Supplementary Table 3.** MOFA+ analysis on IBD dataset. Columns 1-3 contain the percentage of total variance explained by the respective MOFA+ factor in transcriptomics, metabolomics and lipidomics data respectively. 'Functional analysis' contains the top 3 GO Biological Process enriched pathways (gene set enrichment analysis, ranked by p-adj). Only factors discussed in the manuscript are shown here, full data can be investigated in Supplementary Data 1.

| Factor | Transcriptomics | Metabolomics | Functional analysis (GO Biological Process, top 3) |
|---|---|---|---|
| Factor1 | 27.888 | 0.219 | Cellular response to copper ion (NES -3.074, p-adj 1.48e-6); Estrogen metabolic process (NES -3.031, p-adj 2.33e-6); Stress response to copper ion (NES -2.933, p-adj 1.64e-5) |
| Factor2 | 0.0985 | 19.701 | Neutrophil chemotaxis (NES -2.134, p-adj 4.67e-3); Cellular response to copper ion (NES 2.099, p-adj 1.72e-2); Regulation of cell division (NES -2.072, p-adj 9.76e-3) |
| Factor3 | 13.330 | 0.944 | Defense response to virus (NES 3.116, p-adj 1.72e-8); Cellular response to lipopolysaccharide (NES 2.975, p-adj 1.72e-8); Cellular response to biotic stimulus (NES 2.882, p-adj 1.72e-8) |

**Supplementary Table 4**. Overview of selection strategy to prioritize Lemonite predictions for experimental validation. Three metabolites with high strong regulator capacity (high outdegree or high regulator score in the Lemonite network) were selected as described in the main text**.** Information between () represents the expression level of the respective gene in HT29 cells according to DepMap (log(transcripts per million+1)), module number that contains the respective gene and overall rank of the respective metabolite-module score in the Lemonite analysis.

| **C2 carnitine** | **Trigonelline** | **Alpha-glycerophosphocholine** |
|---|---|---|
| HK2 (4.0, 7, 37) | UGT1A7 (0.8, 45, 2) | SLC16A1 (5.2, 2, 182) |
| SCD (7.49, 7, 37) | NETO2 (2.7, 45, 2) | CLCN2 (4.6, 2, 182) |
| CXCL1 (1.8, 7, 37) | C2orf15(3.2, 45, 2) | SLC36A1 (1.75, 2, 182) |
| ETV4 (5.8, 23, 360) | PDK4 (1.7, 45, 2) | NAAA (3.25, 2, 182) |
| ME1 (4.15, 23, 360) | PER3 (1.95, 45, 2) | GUCA2A (2.16, 2, 182) |
| BHLHE40 (2.6, 23, 360) | PDXP (2.9, 45, 2) | GUCA2B (1.4, 2, 182) |
| SLC16A1 (5.2, 2, 121) | | FREM2 (2.3, 26, 101) |
| CDKN2B (2.25, 2, 121) | | NRG4 (0.95, 26, 101) |
| ASS1 (4.0, 7, 37) | | DZIP1L (0.16, 93, 80) |
| APOBEC3B (2.84, 26, 11) | | |
| FREM2 (2.3, 26, 11) | | |
| NRG4 (0.95, 26, 11) | | |
| APOH (4.58, 16, 59) | | |

**Supplementary Table 5**. List of highly-connected metabolites that were excluded from Human1-GEM during the construction of the Lemonite knowledge graph.

| Metabolite name | Metabolite ID in Human1-GEM |
|---|---|
| H+ | MAM02039 |
| H2O | MAM02040 |
| ATP | MAM01371 |
| Pi | MAM02751 |
| ADP | MAM01285 |
| NADPH | MAM02555 |
| NADP+ | MAM0254 |
| O2 | MAM02630 |
| CoA | MAM01597 |
| NAD+ | MAM02552 |
| NADH | MAM02553 |
| PPi | MAM02759 |
| HCO3 | MAM02046 |
| AMP | MAM01334 |

**Supplementary Table 6.** Primers used for qPCR in perturbation experiment.

| Gene | Forward | Reverse |
|---|---|---|
| HK2 | CCC TGC CAC CAG ACT AA | GGA TCA GAG CCA CAA CG |
| SCD | TCT AGC TCC TAT ACC ACC ACC A | TCG TCT CCA ACT TAT CTC CTC C |
| ETV4 | GAT GAA AGC CGG ATA CTT GGA C | TTC GCG CAA GCT CCC ATT T |
| ME1 | CTG CTG ACA CGG AAC CCT C | GAT CTC CTG ACT GTT GAA GGA AG |
| BHLHE40 | GAC GGG GAA TAA AGC GGA GC | CCG GTC ACG TCT CTT TTT CTC |
| SLC16A1 | | |
| CDKN2B | GGG ACT AGT GGA GAA GGT GC | CAT CAT CAT GAC CTG GAT CGC |
| ASS1 | TCC GTG GTT CTG GCC TAC A | GGC TTC CTC GAA GTC TTC CTT |
| APOBEC3B | CGC CAG ACC TAC TTG TGC TAT | CAT TTG CAG CGC CTC CTT AT |
| FREM2 | CCT GCA TGA CCT GGT GTT G | GCC AGT GCG TCG TTG TCT A |
| NRG4 | ATG CCA ACA GAT CAC GAA GAG | AAT GGG CTG GGA ATA GTA GGT |
| APOH | CCC AAG CCA GAT GAT TTA CCA T | ACA GTC CTG TGA GAG GGC A |
| CXCL1 | GCT TGC CTC AAT CCT GCA TC | AGT TGG ATT TGT CAC TGT TCA GC |
| UGT1A7 | CCT CCT TCC CCT ATA TGT GTG T | GCA TCG GCA AAA ACC ATG AAC |
| NETO2 | AGA TGG GCC ATT TGG TTT CTC | TGC TCG AAA TCC CAG TCC TTC |
| C2orf15 | GTT GGA ACC AGC GAC TCA GTT | GCC AGT CCC TTC AAT CCT TGT A |
| PDK4 | GGA GCA TTT CTC GCG CTA CA | ACA GGC AAT TCT TGT CGC AAA |
| PER3 | GCA GAG GAA ATT GGC GGA CA | GGT TTA TTG CGT CTC TCC GAG |
| PDXP | CTG GAG ACC GAC ATC CTC TTT | TTC TAG GCG GGA GAC TCC TG |
| SLC16A1 | ACG CAG AGA CTT CGG AAT GAA | CAT CAC GAT GCC TGC ATT TTT C |

| | | |
|---|---|---|
| CLCN2 | CTG GGT CAC CTA CCC TGT TG | GAG TGT GAG GTA TTC TTT CAG CA |
| SLC36A1 | ACG CAG AGA CTT CGG AAT GAA | CAT CAC GAT GCC TGC ATT TTT C |
| NAAA | CAA CCT GGC CTA CGA GTC C | GCT TGC GTA AGA CAT TCC CAA AA |
| GUCA2A | GTA GCA ACC CGA ACT TTC CAG | GGC AGC GTA GGC ACA GAT TT |
| GUCA2B | CAG AGC ACA CAG TCA GTC TAC A | TCG TCG TTA GCG ATG GTC CT |
| DZIP1L | AGA GCT ACG GGC CAA GCT AA | TTC CCC ATA AAG TTT GGT CCA C |

**Supplementary Table 7.** Reagents and Resources table.

| Reagent/Resource | Version/Catalog number |
|---|---|
| qPCR primers | Supplementary table 6 |
| Alpha-glycerophosphocholine | sc-301813 (Santa Cruz) |
| C2 carnitine | 17-0200-7 (Larodan) |
| Trigonelline | 1686411 (Sigma Aldrich) |
| Java | |
| Nextflow | 25.04.4 |
| R | 4.4 |
| DESeq2 | 1.46.0 |
| Biomart | 2.26.1 |
| data.table | 1.17.8 |
| Tidyverse | 2.0.0 |
| Dplyr | 1.1.4 |
| Ggplot2 | 4.0.0 |
| Caret | 7.0-1 |
| IMIFA | 2.2.0 |
| MixOmics | 6.30.0 |
| Clusterprofiler | 4.14.6 |
| Org.Hs.eg.db | 3.20.0 |
| ReactomePA | 1.50.0 |
| Reactome.db | 1.89.0 |
| MOFA2 | 1.16.0 |
| MultiAssayExperiment | 1.42.0 |
| reticulate | 1.43.0 |
| MOFAdata | 1.22.0 |
| MOFAcellulaR | 0.0.0.9000 |
| cosmosR | 1.14.0 |
| Liana | 0.1.14 |
| decoupleR | 2.12.0 |
| Reshape2 | 1.4.4 |
| Ggfortify | 0.4.19 |
| Pheatmap | 1.0.13 |
| gridExtra | 2.3 |
| RcolorBrewer | 1.1-3 |
| ggvenn | 0.1.19 |
| AnnotationDbi | 1.68.0 |
| Readxl | 1.4.5 |
| FSA | 0.10.0 |
| Rstatix | 0.7.2 |
| drc | 3.0-1 |
| stringr | 1.5.2 |
| Fgsea | 1.32.4 |
| Enrichplot | 1.26.6 |
| WGCNA | 1.73 |
| enrichR | 3.4 |
| BiocParallel | 1.40.2 |
| BiocManager | 1.30.26 |
| Future | 1.67.0 |
| Future.apply | 1.20.0 |

| | |
|---|---|
| OmnipathR | 3.17.4 |
| Python | 3.11.11 |
| Pandas | 2.2.3 |
| Numpy | 2.2.6 |
| Scipy | 1.15.3 |
| Statsmodels | 0.14.4 |
| Matplotlib | 3.10.3 |
| Seaborn | 0.13.2 |
| networkx | 3.4.2 |
| Scikit-learn | 1.6.1 |
| Plotly | 6.1.1 |
| mygene | 3.2.2 |
| Scipy | 1.15.3 |
| adjustText | 1.3.0 |
| megago | 1.0.0.2021.4 |
| Chembl_webresource_client | 0.10.9 |
| rdkit | 2025.3.2 |
| tqdm | 4.67.1 |
| requests | 2.32.3 |
| Upsetplot | 0.9.0 |
| Pybiomart | 0.2.0 |
| Streamlit | 1.45.1 |
| Typing_extensions | 4.13.2 |

### Supplementary figures

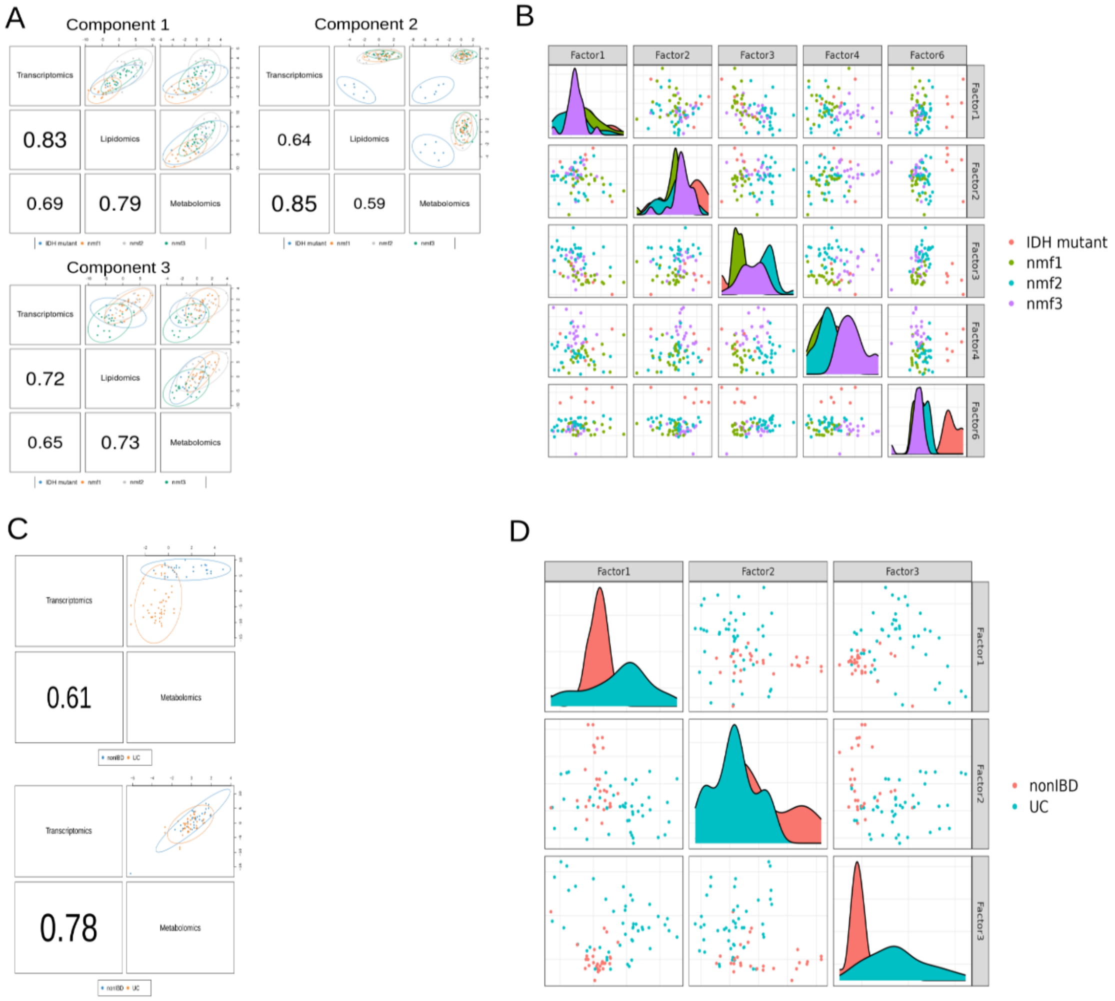

**Supplementary Fig. 1. Exploratory analysis of GBM (A,B) and IBD (C,D) cohorts through integration of metabolomics transcriptomics data. A.** Components 1-3 of the MixOmics DIABLO model inferred on transcriptomics, metabolomics and lipidomics data. Numbers represent absolute correlation values between omics in the respective component. Scatter plots show sample distributions in each component colored by multi-omics GBM subtype. **B.** Sample distribution in MOFA+ factors 1,2,3,4 and 6 in GBM. **C.** Components 1 and 5 of the MixOmics DIABLO model inferred on transcriptomics and metabolomics data. Numbers represent absolute correlation values between omics in the respective component. Scatter plots show sample distributions in each component colored by multi-omics GBM subtype. **D.** Sample distributions in MOFA+ latent factors 1,2 and 3.

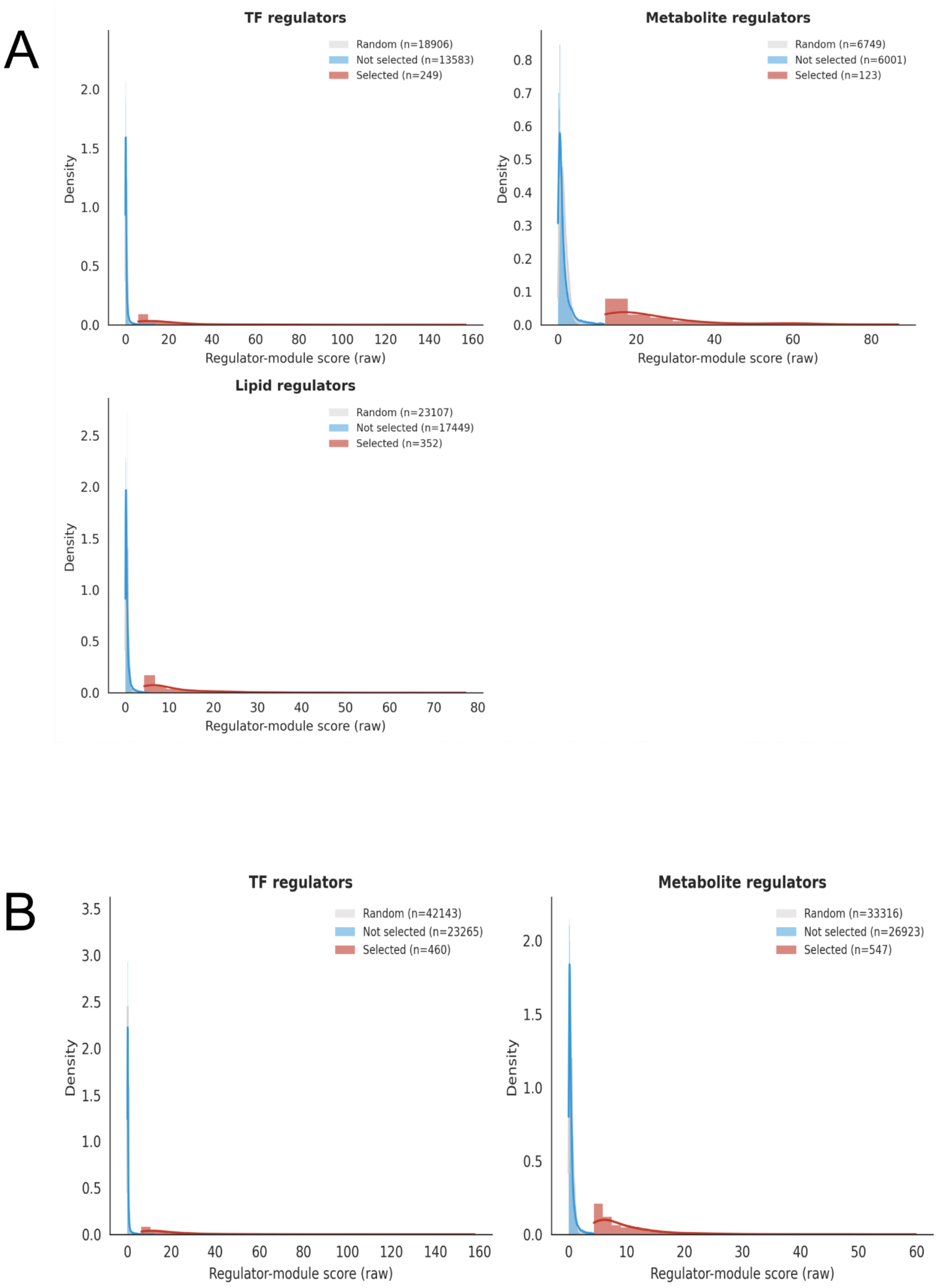

**Supplementary Fig. 2. Regulator score distributions (random, true and selected) in Lemonite networks.** Distribution of regulator scores (per type) in the GBM (A) and IBD (B) Lemonite networks. Shown are the scores for random regulators, all 'true' regulators outputted by Lemonite, and the regulators that are retained after selection as described in the Methods section.

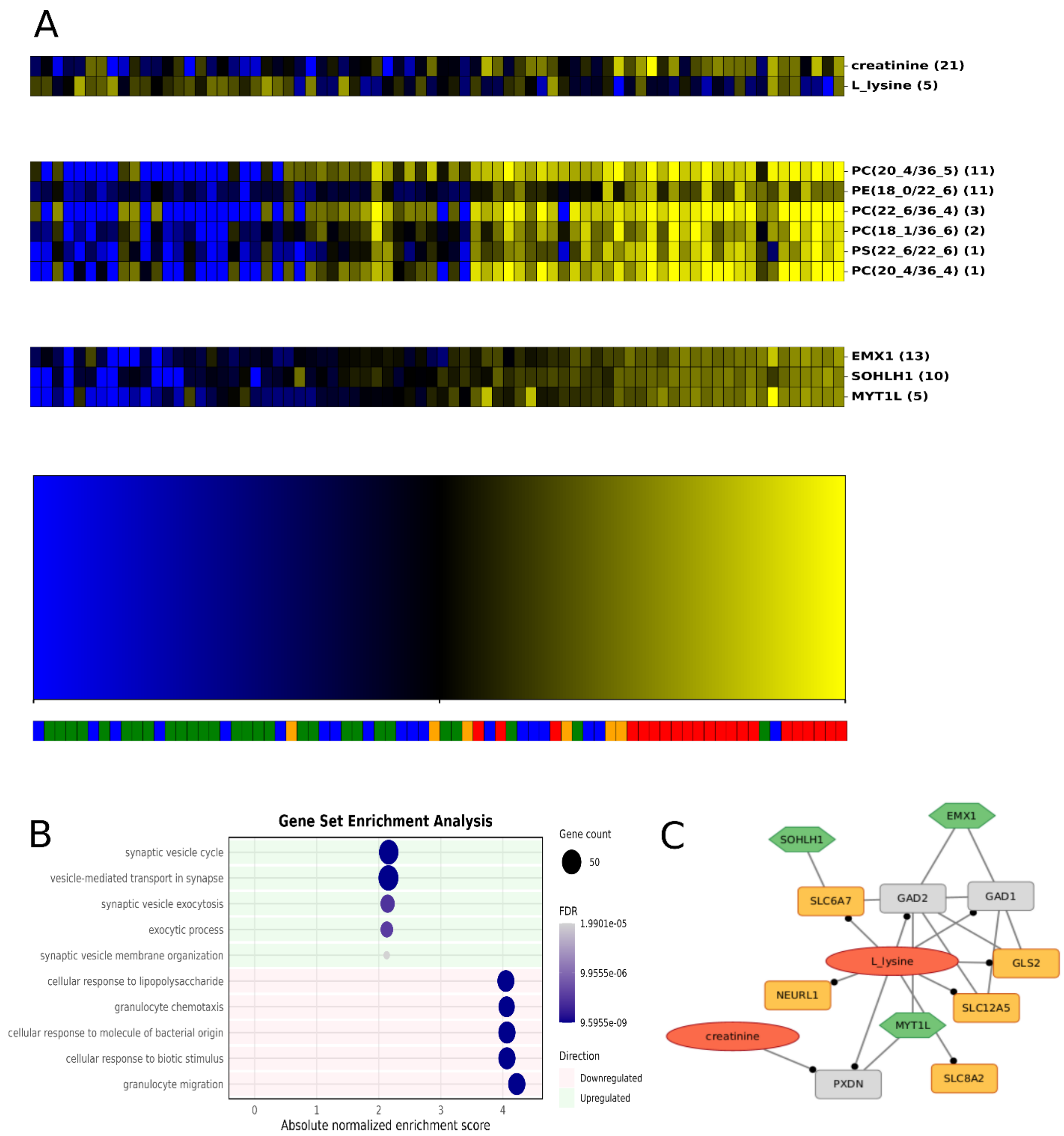

**Supplementary Fig. 3. GBM module 0. A.** GBM module 0, represented by a blue-to-yellow bar as there are too many genes to display, colored by Z-scored expression (blue: -2; yellow: +2). TF, lipid and metabolite regulators with respective regulator scores above the module. Annotations at the bottom indicate multi-omic subtypes (proneural/nmf1: red; mesenchymal/nmf2: green; classical/nmf3: blue; IDH mutant: orange). The complete module figure can be accessed at www.lemonite.ugent.be. **B.** Gene set enrichment analysis with GO biological process 2025 for module 0. **C.** Module 0 in the Lemonite KG. Metabolite regulators colored in red, TF regulators in green, module genes in orange. Nodes that connect metabolite regulators to TF regulators through a single PPI in the LemonIte KG colored in grey. Grey boxes represent genes involved in a protein-protein interaction that connects metabolite regulators to TF regulators in the Lemonite KG. Edges ending in a dot represent metabolic pathway interactions, normal lines represent PPIs and 'other' interactions.

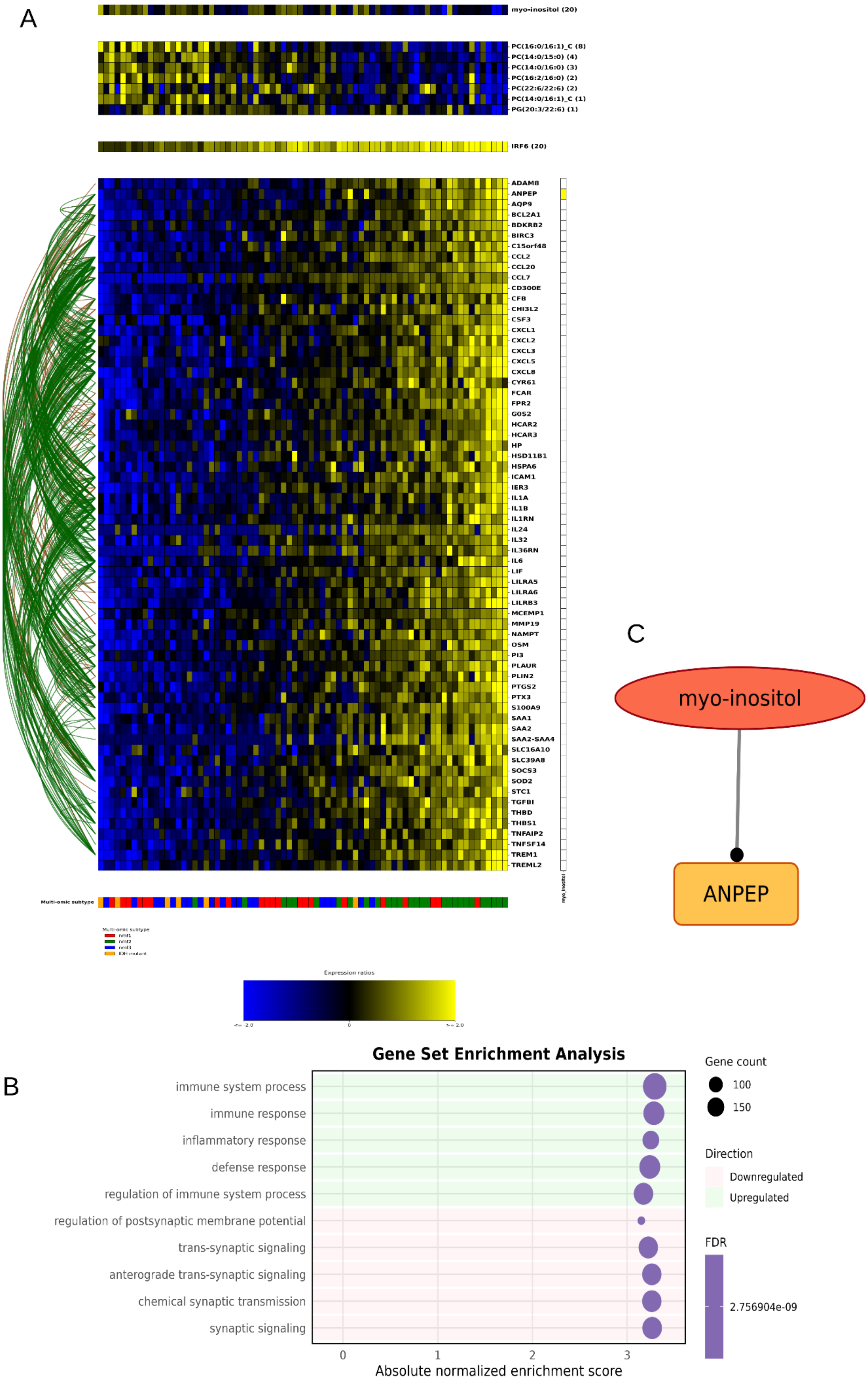
A
myo-inositol (20)
PC(16:0/16:1)_C (8)
PC(14:0/15:0) (4)
PC(14:0/16:0) (3)
PC(16:2/16:0) (2)
PC(22:6/22:6) (2)
PC(14:0/16:1)_C (1)
PG(20:3/22:6) (1)
IRF6 (20)
ADAM8
ANPEP
AQP9
BCL2A1
BDKRB2
BIRC3
C15orf48
CCL2
CCL20
CCL7
CD300E
CFB
CHI3L2
CSF3
CXCL1
CXCL2
CXCL3
CXCL5
CXCL8
CYR61
FCAR
FPR2
G0S2
HCAR2
HCAR3
HP
HSD11B1
HSPA6
ICAM1
IER3
IL1A
IL1B
IL1RN
IL24
IL32
IL36RN
IL6
LIF
LILRA5
LILRA6
LILRB3
MCEMP1
MMP19
NAMPT
OSM
PI3
PLAUR
PLIN2
PTGS2
PTX3
S100A9
SAA1
SAA2
SAA2-SAA4
SLC16A10
SLC39A8
SOCS3
SOD2
STC1
TGFBI
THBD
THBS1
TNFAIP2
TNFSF14
TREM1
TREML2
myo_inositol
Multi-omic subtype
Expression ratios
B
Gene Set Enrichment Analysis
immune system process
immune response
inflammatory response
defense response
regulation of immune system process
regulation of postsynaptic membrane potential
trans-synaptic signaling
anterograde trans-synaptic signaling
chemical synaptic transmission
synaptic signaling
0
1
2
3
Absolute normalized enrichment score
Gene count
100
150
Direction
Downregulated
Upregulated
FDR
2.756904e-09
C
myo-inositol
ANPEP

**Supplementary Fig. 4. GBM module 5. A.** GBM module 5 colored by Z-scored expression (blue: -2; yellow: +2). TF, lipid and metabolite regulators with respective regulator scores above the module. Annotations at the bottom indicate multi-omic subtypes (proneural/nmf1: red; mesenchymal/nmf2: green; classical/nmf3: blue; IDH mutant: orange). **B.** Gene set enrichment analysis with GO biological process 2025 for module 5. **C.** Module 5 in the Lemonite KG. Metabolite regulators colored in red, TF regulators in green, module genes in orange. Nodes that connect metabolite regulators to TF regulators through a single PPI in the LemonIte KG colored in grey. Grey boxes represent genes involved in a protein-protein interaction that connects metabolite regulators to TF regulators in the Lemonite KG. Edges ending in a dot represent metabolic pathway interactions, normal lines represent PPIs and ‘other’ interactions.

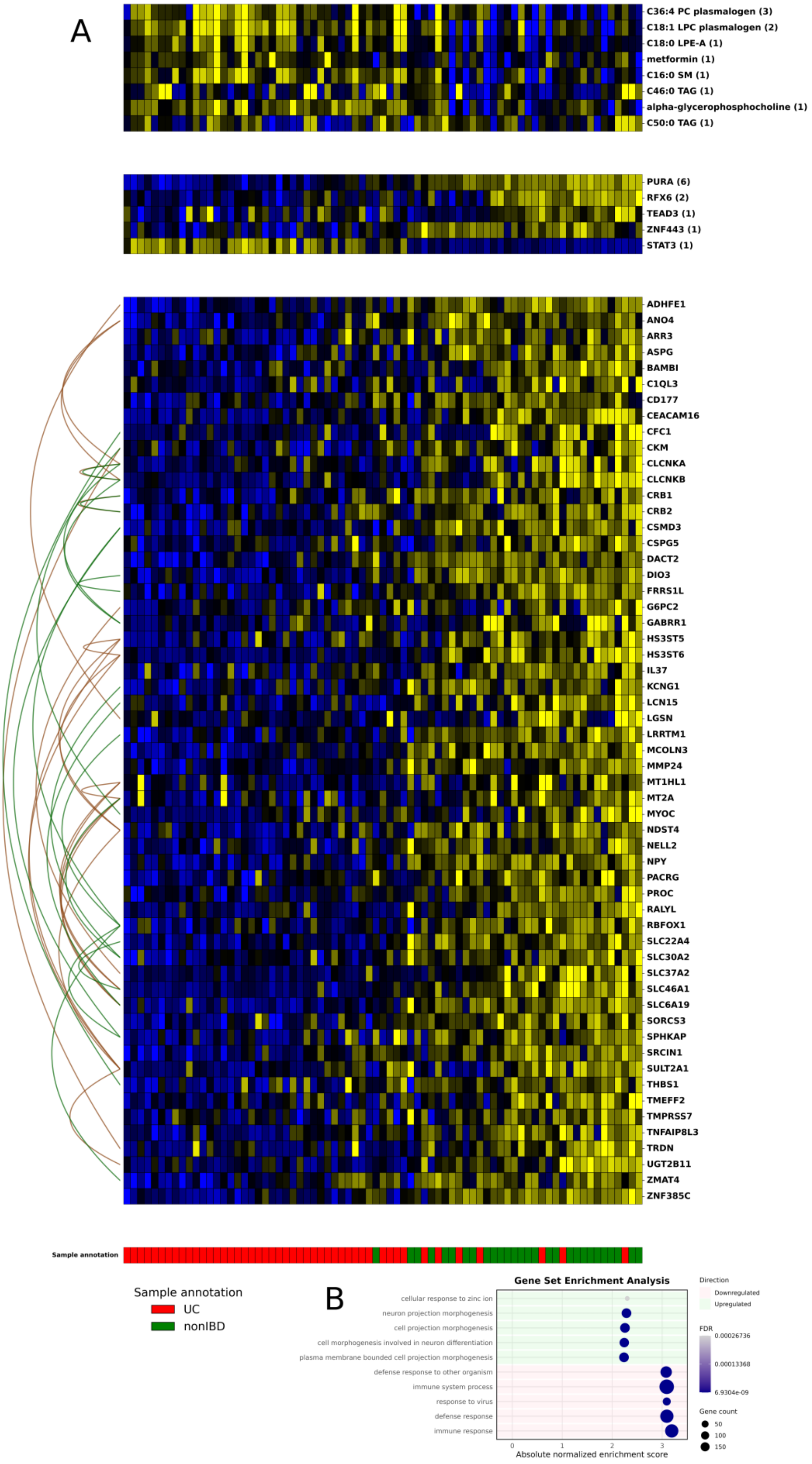

A
C36:4 PC plasmalogen (3)
C18:1 LPC plasmalogen (2)
C18:0 LPE-A (1)
metformin (1)
C16:0 SM (1)
C46:0 TAG (1)
alpha-glycerophosphocholine (1)
C50:0 TAG (1)
PURA (6)
RFX6 (2)
TEAD3 (1)
ZNF443 (1)
STAT3 (1)
ADHFE1
ANO4
ARR3
ASPG
BAMBI
C1QL3
CD177
CEACAM16
CFC1
CKM
CLCNKA
CLCNKB
CRB1
CRB2
CSMD3
CSPG5
DACT2
DIO3
FRRS1L
G6PC2
GABRR1
HS3ST5
HS3ST6
IL37
KCNG1
LCN15
LGSN
LRRTM1
MCOLN3
MMP24
MT1HL1
MT2A
MYOC
NDST4
NELL2
NPY
PACRG
PROC
RALYL
RBFOX1
SLC22A4
SLC30A2
SLC37A2
SLC46A1
SLC6A19
SORCS3
SPHKAP
SRCIN1
SULT2A1
THBS1
TMEFF2
TMPRSS7
TNFAIP8L3
TRDN
UGT2B11
ZMAT4
ZNF385C
Sample annotation
Sample annotation
UC
nonIBD
B
Gene Set Enrichment Analysis
cellular response to zinc ion
neuron projection morphogenesis
cell projection morphogenesis
cell morphogenesis involved in neuron differentiation
plasma membrane bounded cell projection morphogenesis
defense response to other organism
immune system process
response to virus
defense response
immune response
0
1
2
3
Absolute normalized enrichment score
Direction
Downregulated
Upregulated
FDR
0.00026736
0.00013368
6.9304e-09
Gene count
50
100
150

**Supplementary Fig. 5. IBD module 49. A.** IBD module 49, colored by Z-scored expression (blue: -2; yellow: +2). TF, lipid and metabolite regulators with respective regulator scores above the module. **B.** Gene set enrichment analysis with GO biological process 2025 for module 49. **C.** Module 49 in the Lemonite KG. Metabolite regulators colored in red, TF regulators in green, module genes in orange. Nodes that connect metabolite regulators to TF regulators through a single PPI in the LemonIte KG colored in grey. Grey boxes represent genes involved in a protein-protein interaction that connects metabolite regulators to TF regulators in the Lemonite KG. Edges ending in a dot represent metabolic pathway interactions, normal lines represent PPIs and ‘other’ interactions.

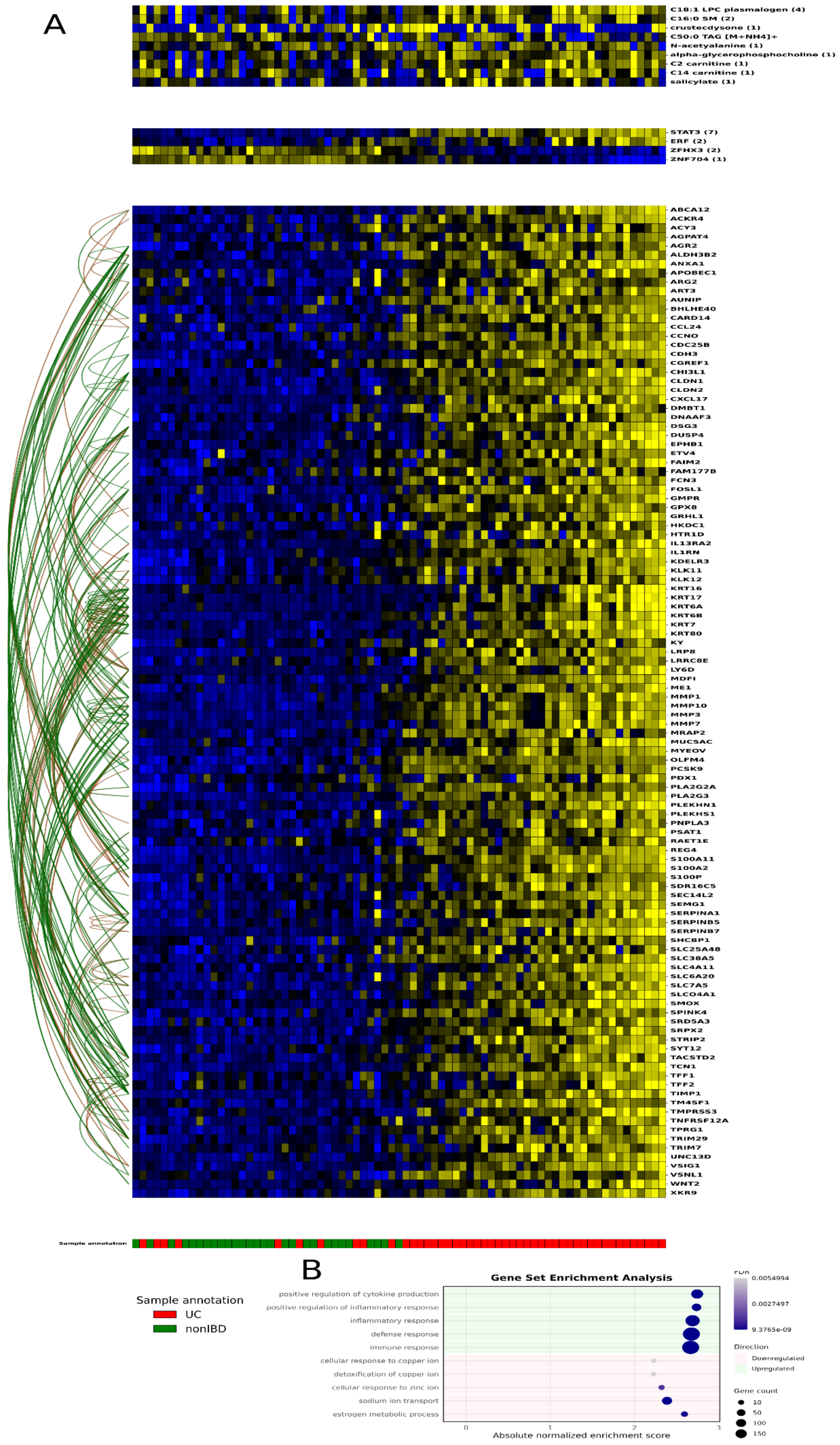
A
C18:1 LPC plasmalogen (4)
C16:0 SM (2)
crustecdysone (1)
C50:0 TAG [M+NH4]+
N-acetyalanine (1)
alpha-glycerophosphocholine (1)
C2 carnitine (1)
C14 carnitine (1)
salicylate (1)
STAT3 (7)
ERF (2)
ZFHX3 (2)
ZNF704 (1)
ABCA12
ACKR4
ACY3
AGPAT4
AGR2
ALDH3B2
ANXA1
APOBEC1
ARG2
ART3
AUNIP
BHLHE40
CARD14
CCL24
CCNO
CDC25B
CDH3
CGREF1
CHI3L1
CLDN1
CLDN2
CXCL17
DMBT1
DNAAF3
DSG3
DUSP4
EPHB1
ETV4
FAIM2
FAM177B
FCN3
FOSL1
GMPR
GPX8
GRHL1
HKDC1
HTR1D
IL13RA2
IL1RN
KDELR3
KLK11
KLK12
KRT16
KRT17
KRT6A
KRT6B
KRT7
KRT80
KY
LRP8
LRRC8E
LY6D
MDFI
ME1
MMP1
MMP10
MMP3
MMP7
MRAP2
MUC5AC
MYEOV
OLFM4
PCSK9
PDX1
PLA2G2A
PLA2G3
PLEKHN1
PLEKHS1
PNPLA3
PSAT1
RAET1E
REG4
S100A11
S100A2
S100P
SDR16C5
SEC14L2
SEMG1
SERPINA1
SERPINB5
SERPINB7
SHCBP1
SLC25A48
SLC38A5
SLC4A11
SLC6A20
SLC7A5
SLCO4A1
SMOX
SPINK4
SRD5A3
SRPX2
STRIP2
SYT12
TACSTD2
TCN1
TFF1
TFF2
TIMP1
TM4SF1
TMPRSS3
TNFRSF12A
TPRG1
TRIM29
TRIM7
UNC13D
VSIG1
VSNL1
WNT2
XKR9
Sample annotation
B
Gene Set Enrichment Analysis
positive regulation of cytokine production
positive regulation of inflammatory response
inflammatory response
defense response
immune response
cellular response to copper ion
detoxification of copper ion
cellular response to zinc ion
sodium ion transport
estrogen metabolic process
0
1
2
3
Absolute normalized enrichment score
FDR
0.0054994
0.0027497
9.3765e-09
Direction
Downregulated
Upregulated
Gene count
10
50
100
150
Sample annotation
UC
nonIBD

**Supplementary Fig. 6. IBD module 23. A.** IBD module 23, colored by Z-scored expression (blue: -2; yellow: +2). TF, lipid and metabolite regulators with respective regulator scores above the module. **B.** Gene set enrichment analysis with GO biological process 2025 for module 23.

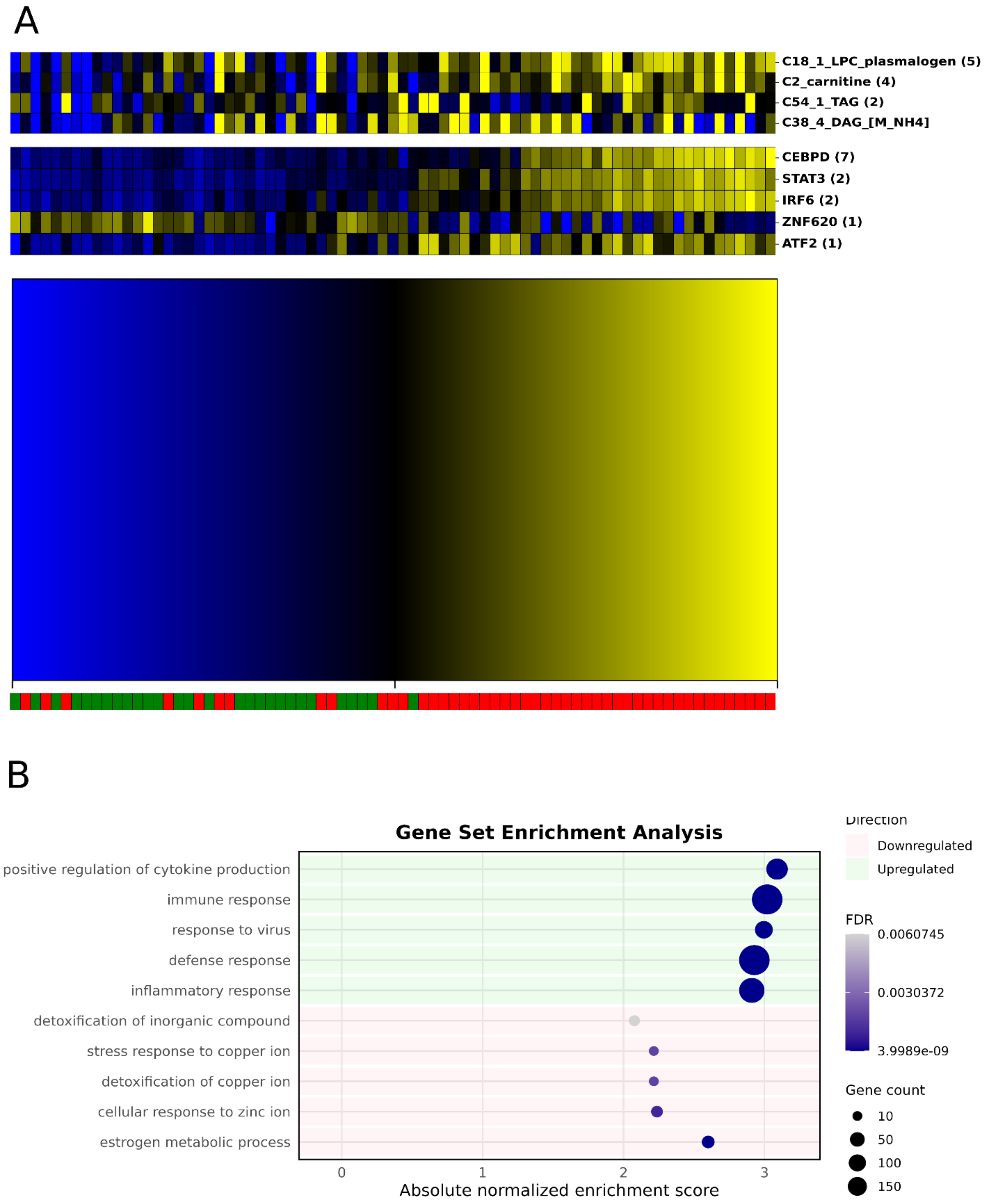

**Supplementary Fig. 7. IBD module 7.** **A.** IBD module 7, colored by Z-scored expression (blue: -2; yellow: +2). TF, lipid and metabolite regulators with respective regulator scores above the module. **B.** Gene set enrichment analysis with GO biological process 2025 for module 7.

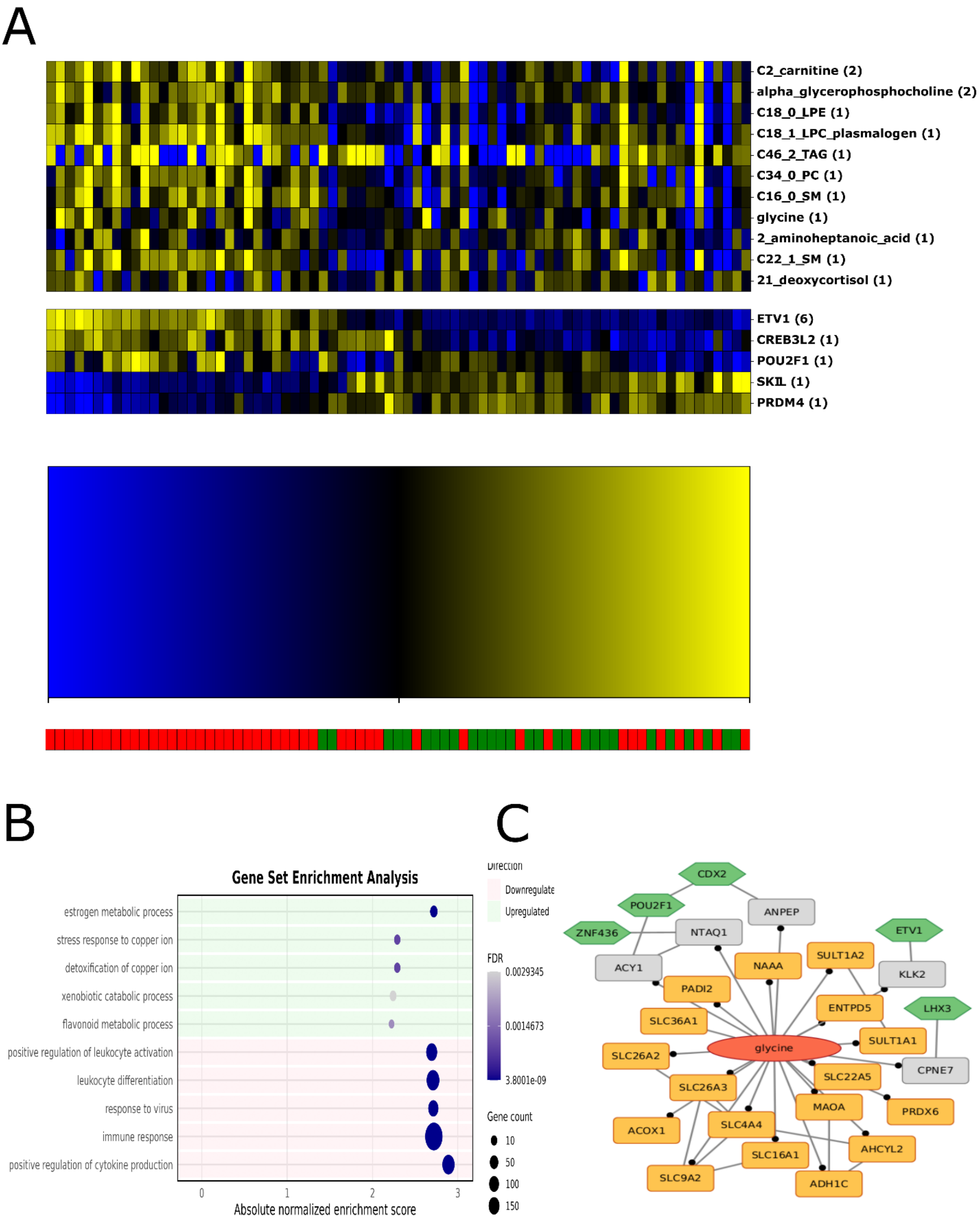

**Supplementary Fig. 8. IBD module 2. A.** IBD module 2, colored by Z-scored expression (blue: -2; yellow: +2). TF, lipid and metabolite regulators with respective regulator scores above the module. **B.** Gene set enrichment analysis with GO biological process 2025 for module 2. **C.** Module 2 in the Lemonite KG. Metabolite regulators colored in red, TF regulators in green, module genes in orange. Nodes that connect metabolite regulators to TF regulators through a single PPI in the LemonIte KG colored in grey. Grey boxes represent genes involved in a protein-protein interaction that connects metabolite regulators to TF regulators in the Lemonite KG. Edges ending in a dot represent metabolic pathway interactions, normal lines represent PPIs and ‘other’ interactions.

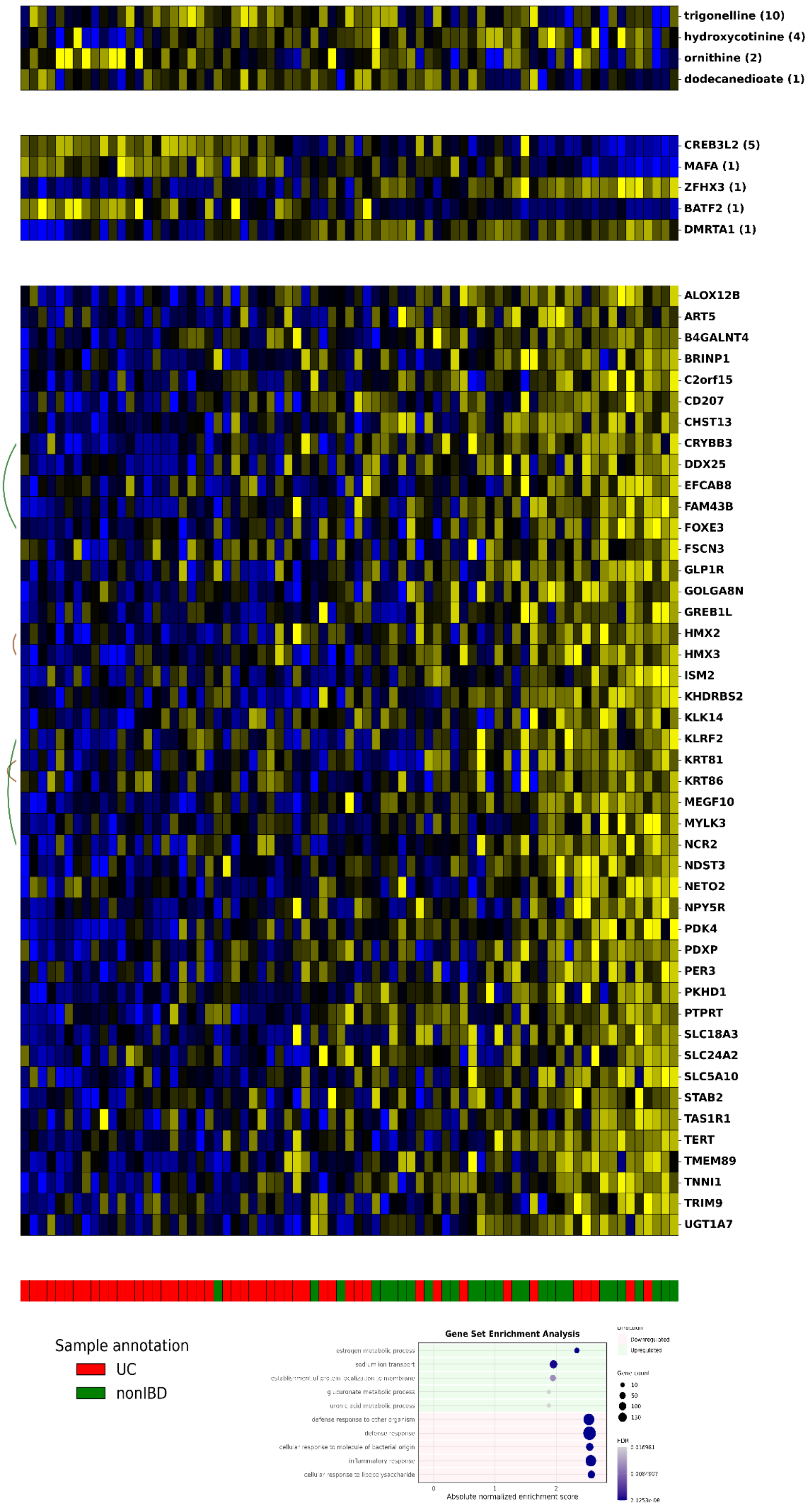
trigonelline (10)
hydroxycotinine (4)
ornithine (2)
dodecanedioate (1)
CREB3L2 (5)
MAFA (1)
ZFHX3 (1)
BATF2 (1)
DMRTA1 (1)
ALOX12B
ART5
B4GALNT4
BRINP1
C2orf15
CD207
CHST13
CRYBB3
DDX25
EFCAB8
FAM43B
FOXE3
FSCN3
GLP1R
GOLGA8N
GREB1L
HMX2
HMX3
ISM2
KHDRBS2
KLK14
KLRF2
KRT81
KRT86
MEGF10
MYLK3
NCR2
NDST3
NETO2
NPY5R
PDK4
PDXP
PER3
PKHD1
PTPRT
SLC18A3
SLC24A2
SLC5A10
STAB2
TAS1R1
TERT
TMEM89
TNNI1
TRIM9
UGT1A7
Sample annotation
UC
nonIBD
Gene Set Enrichment Analysis
estrogen metabolic process
sodium ion transport
establishment of protein localization to membrane
glucuronate metabolic process
uronic acid metabolic process
defense response to other organism
defense response
cellular response to molecule of bacterial origin
inflammatory response
cellular response to lipopolysaccharide
0
1
2
Absolute normalized enrichment score
Downregulated
Upregulated
Gene count
10
50
100
150
FDR

**Supplementary Fig. 9. IBD module 45. A.** IBD module 45, colored by Z-scored expression (blue: -2; yellow: +2). TF, lipid and metabolite regulators with respective regulator scores above the module. **B.** Gene set enrichment analysis with GO biological process 2025 for module 45. **C.** Module 45 in the Lemonite KG. Metabolite regulators colored in red, TF regulators in green, module genes in orange. Nodes that connect metabolite regulators to TF regulators through a single PPI in the LemonIte KG colored in grey. Grey boxes represent genes involved in a protein-protein interaction that connects metabolite regulators to TF regulators in the Lemonite KG. Edges ending in a dot represent metabolic pathway interactions, normal lines represent PPIs and ‘other’ interactions.

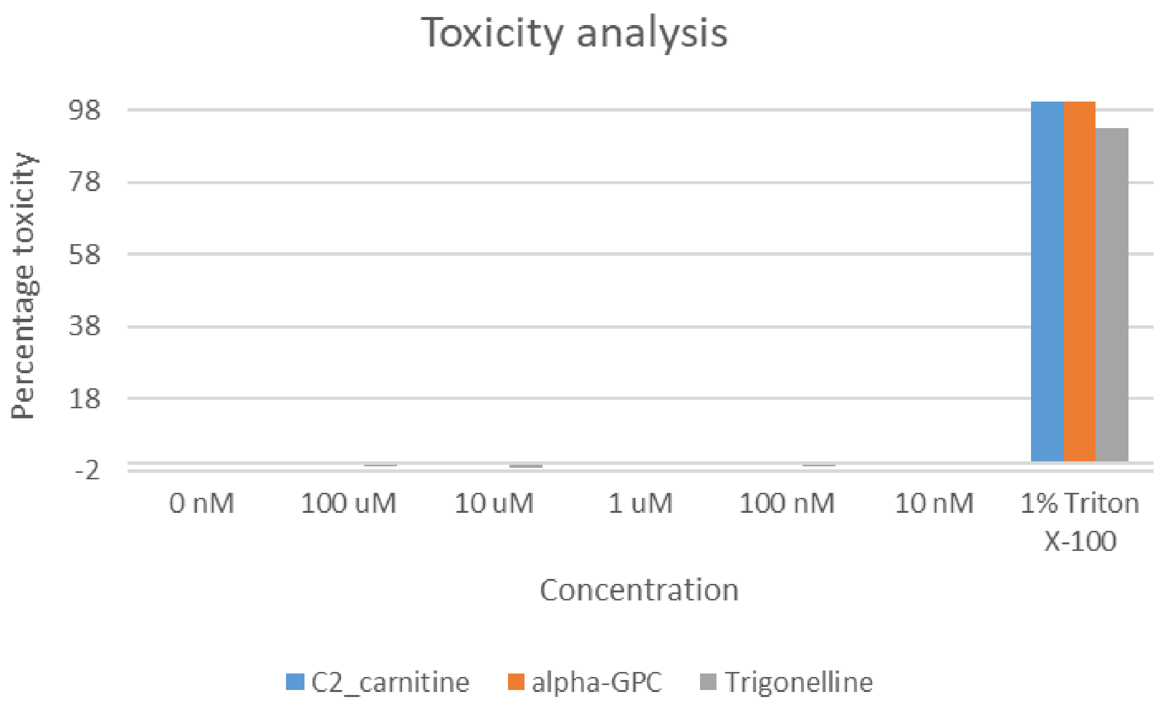

**Supplementary Fig. 10. Cytotoxicity analysis.** HT29 cells were perturbed with C2 carnitine, α-glycerophosphocholine (alpha-GPC) or trigonelline at concentrations between 10nM and 100 µM, and cell viability was measured 24h after treatment.

**Supplementary text**

POU2F3 and creatinine were identified as negative regulators of GBM module 43, next to several triglycerides as positive regulators. POU2F3 is a candidate tumor suppressor protein, while creatine, of which creatinine is a breakdown product, serves as an energy reserve product that promotes growth in GBM stem cells that are progenitors of GBM proneural- and mesenchymal-like cells [1]. In the hypoxic tumor niche, myeloid-to-tumor transfer of creatine fuels tumor growth [2]. Several triglycerides (TG) were assigned as regulator to module 43, while these molecules are known to act as energy reserves in lipid droplets within GBM cells [3].

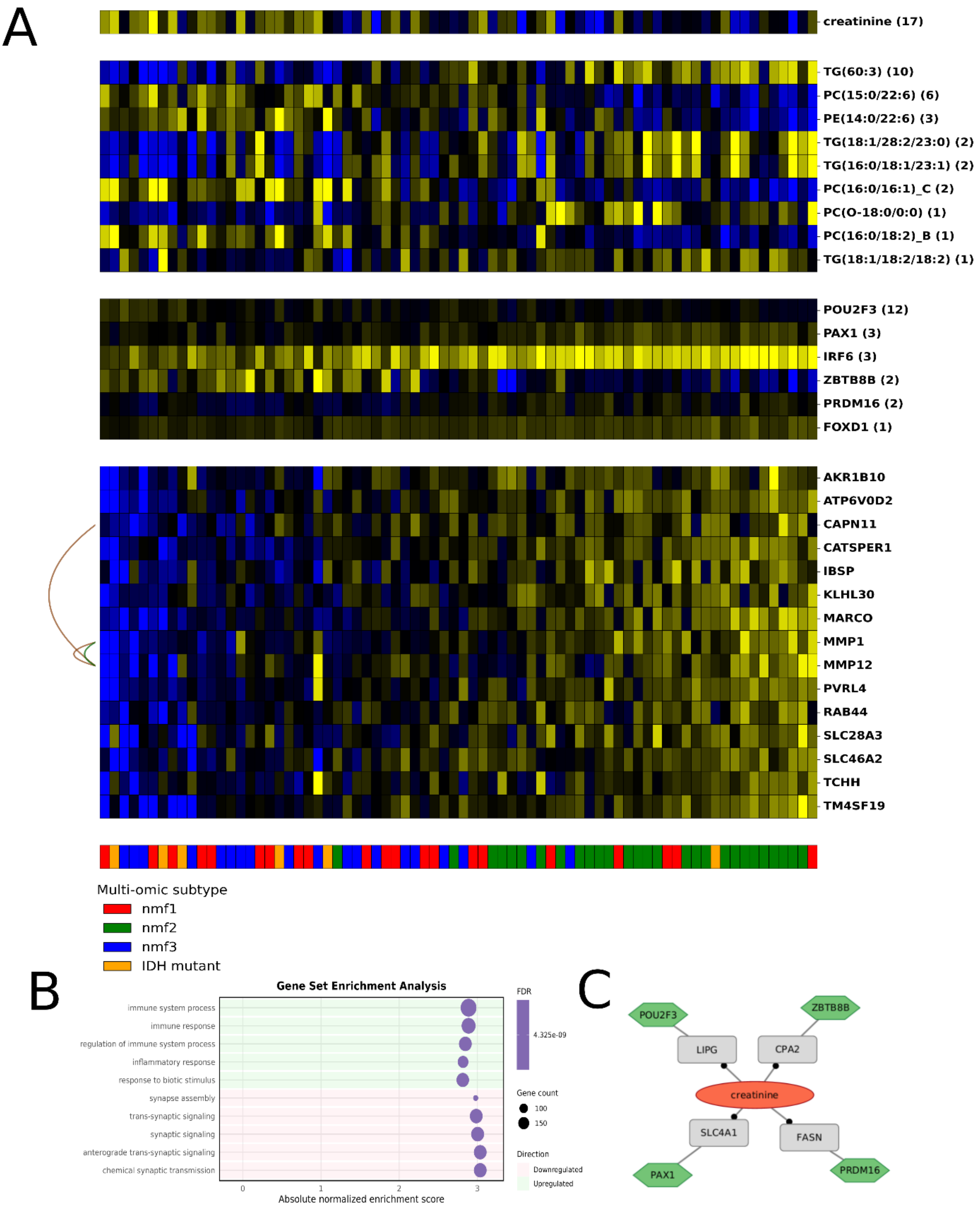

**Supplementary Fig. 11. GBM module 43. A.** GBM module 43 colored by Z-scored expression (blue: -2; yellow: +2). TF, lipid and metabolite regulators with respective regulator scores above the module. Annotations at the bottom indicate multi-omic subtypes (proneural/nmf1: red; mesenchymal/nmf2: green; classical/nmf3: blue; IDH mutant: orange). **B.** Gene set enrichment analysis with GO biological process 2025 for module 43. **C.** Module 43 in the Lemonite KG. Metabolite regulators colored in red, TF regulators in green, module genes in orange. Nodes that connect metabolite regulators to TF regulators through a single PPI in the LemonIte KG colored in grey. Grey boxes represent genes involved in a protein-protein interaction that connects metabolite regulators to TF regulators in the Lemonite KG. Edges ending in a dot represent metabolic pathway interactions, normal lines represent PPIs and ‘other’ interactions. GBM module 43 colored by Z-scored expression (blue: -2; yellow: +2). TF, lipid and metabolite regulators with respective regulator scores are shown above the module. Annotations on the bottom indicate multi-omics subtypes (proneural/nmf1: red; mesenchymal/nmf2: green; classical/nmf3: blue; IDH mutant: orange), yellow boxes on the right represent metabolite-gene interactions that are present in the Lemonite KG. Lines on the left represent physical PPIs (green) or HumanNet functional interactions (brown) between module genes.

GBM module 13 was significantly differentially upregulated in mesenchymal-like (nmf2) samples (p-adj 1.73e-7) and exhibited strong PPI enrichment (p-adj 8.31e-81). This module was enriched for immune response and regulation of immune system processes, including cytokine receptor activity and interleukin signalling (Supplementary Data 1). Phosphoric acid (spectral match), creatinine and N-formylglycine were identified as negative metabolic regulators, while IRF6, FOXD1 and FOXD2 acted as transcriptional regulators. Module 13 retrieved a known interaction between creatinine and the module gene ACP5.

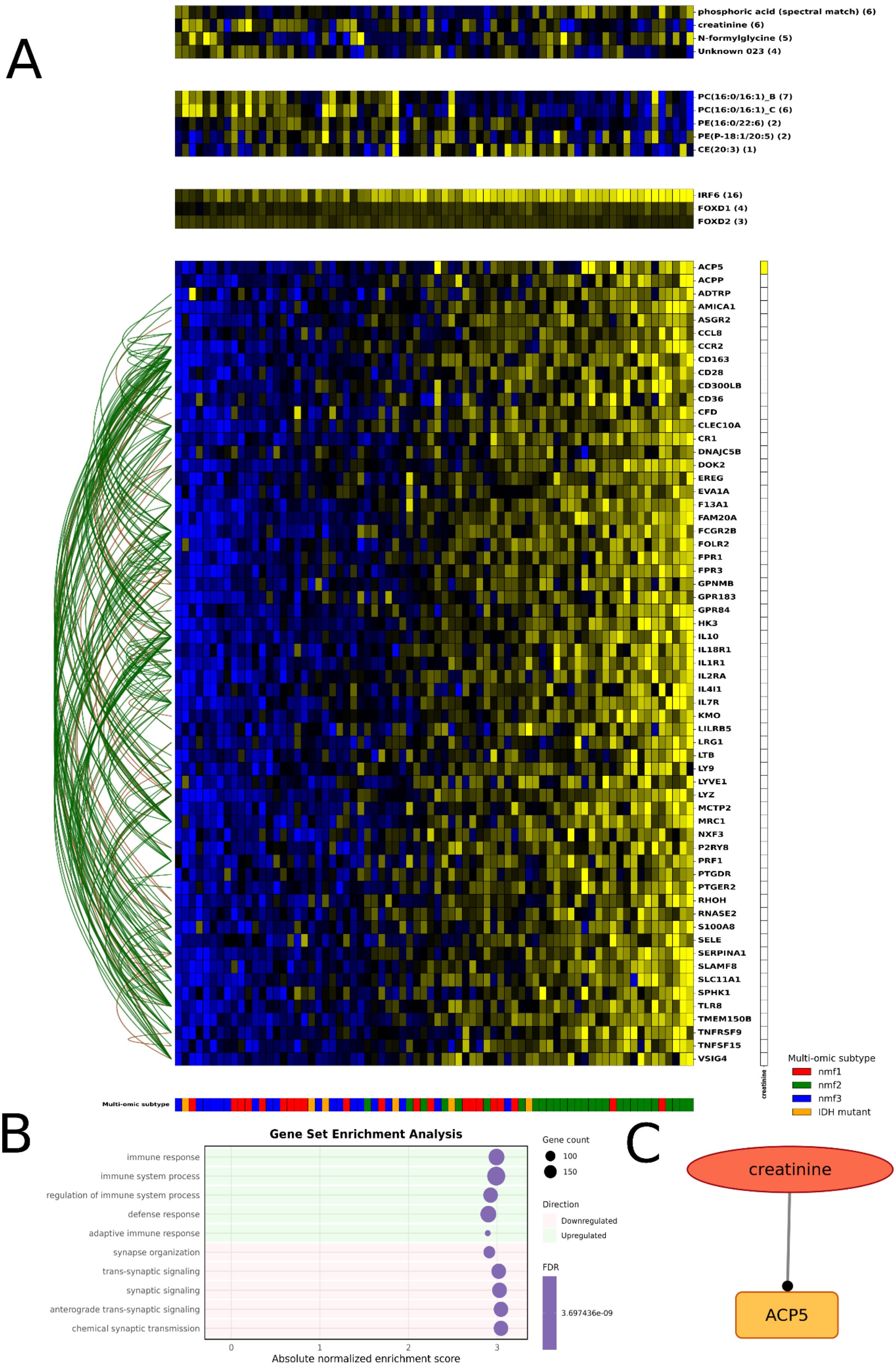
A
phosphoric acid (spectral match) (6)
creatinine (6)
N-formylglycine (5)
Unknown 023 (4)
PC(16:0/16:1)_B (7)
PC(16:0/16:1)_C (6)
PE(16:0/22:6) (2)
PE(P-18:1/20:5) (2)
CE(20:3) (1)
IRF6 (16)
FOXD1 (4)
FOXD2 (3)
ACP5
ACPP
ADTRP
AMICA1
ASGR2
CCL8
CCR2
CD163
CD28
CD300LB
CD36
CFD
CLEC10A
CR1
DNAJC5B
DOK2
EREG
EVA1A
F13A1
FAM20A
FCGR2B
FOLR2
FPR1
FPR3
GPNMB
GPR183
GPR84
HK3
IL10
IL18R1
IL1R1
IL2RA
IL4I1
IL7R
KMO
LILRB5
LRG1
LTB
LY9
LYVE1
LYZ
MCTP2
MRC1
NXF3
P2RY8
PRF1
PTGDR
PTGER2
RHOH
RNASE2
S100A8
SELE
SERPINA1
SLAMF8
SLC11A1
SPHK1
TLR8
TMEM150B
TNFRSF9
TNFSF15
VSIG4
creatinine
Multi-omic subtype
Multi-omic subtype
nmf1
nmf2
nmf3
IDH mutant
B
Gene Set Enrichment Analysis
immune response
immune system process
regulation of immune system process
defense response
adaptive immune response
synapse organization
trans-synaptic signaling
synaptic signaling
anterograde trans-synaptic signaling
chemical synaptic transmission
0
1
2
3
Absolute normalized enrichment score
Gene count
100
150
Direction
Downregulated
Upregulated
FDR
3.697436e-09
C
creatinine
ACP5

**Supplementary Fig. 121. GBM module 13. A.** GBM module 13 colored by Z-scored expression (blue: -2; yellow: +2). TF, lipid and metabolite regulators with respective regulator scores above the module. Annotations at the bottom indicate multi-omic subtypes (proneural/nmf1: red; mesenchymal/nmf2: green; classical/nmf3: blue; IDH mutant: orange). **B.** Gene set enrichment analysis with GO biological process 2025 for module 13. **C.** Module 13 in the Lemonite KG. Metabolite regulators colored in red, TF regulators in green, module genes in orange. Nodes that connect metabolite regulators to TF regulators through a single PPI in the LemonIte KG colored in grey. Grey boxes represent genes involved in a protein-protein interaction that connects metabolite regulators to TF regulators in the Lemonite KG. Edges ending in a dot represent metabolic pathway interactions, normal lines represent PPIs and 'other' interactions.

Module 23 was upregulated in mesenchymal-like (nmf2) samples compared to other subtypes (p-adj 1.03e-3) and significantly enriched for PPIs (p-adj 1.10e-4). Accordingly, the module was mostly expressed in monocytes and TAMs (main manuscript Figure 4C). Functionally, it was associated with adaptive immune response and response to bacteria (Supplementary data 1)**.** O-phosphocolamine and two unknown metabolites were identified as regulators, while BATF and TBX21 were key transcriptional regulators. BATF is a key regulator of myeloid cell differentiation, in line with the module being upregulated in mesenchymal-like cells and being primarily expressed in monocytes and TAMs.

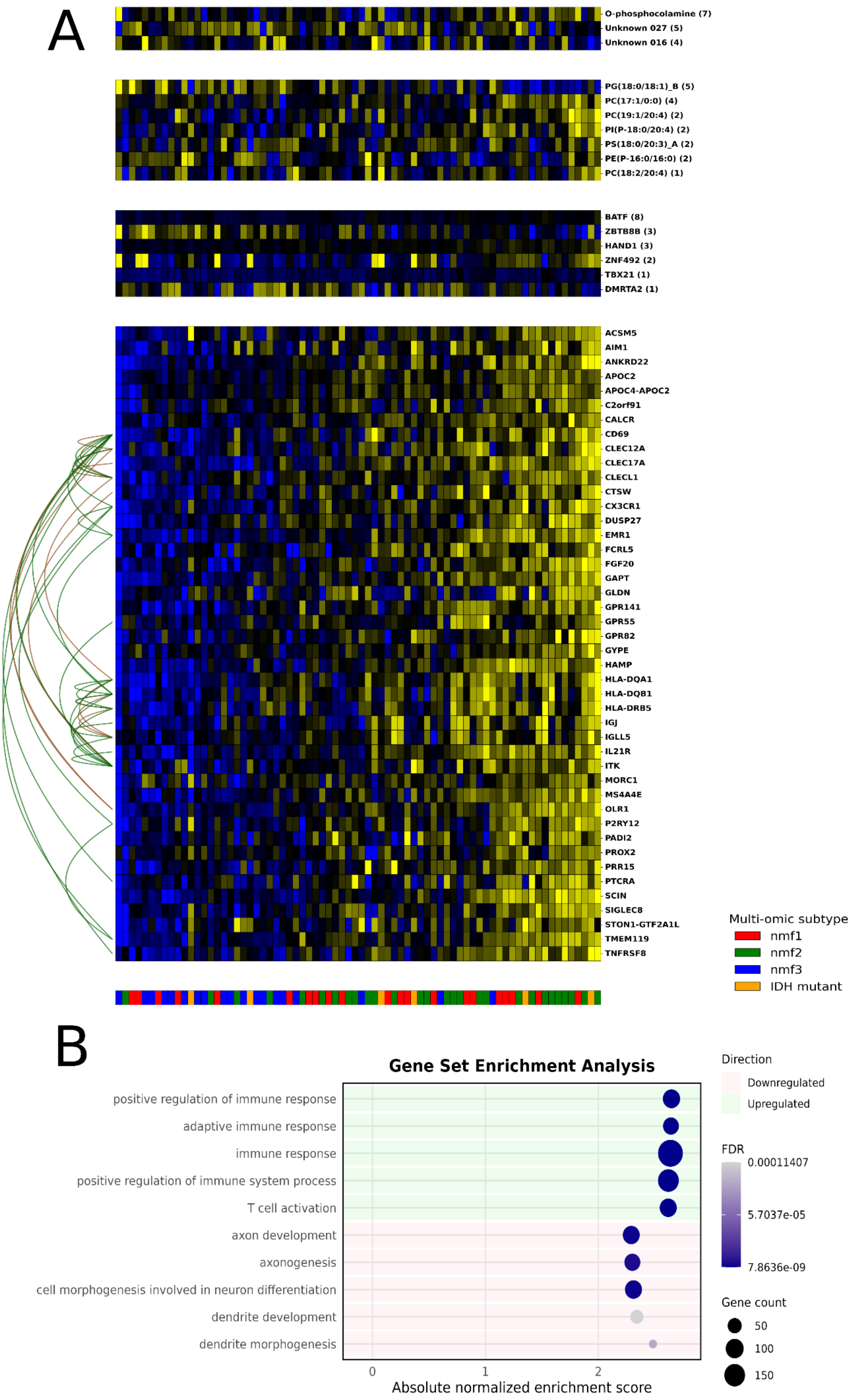
A
O-phosphocolamine (7)
Unknown 027 (5)
Unknown 016 (4)
PG(18:0/18:1)_B (5)
PC(17:1/0:0) (4)
PC(19:1/20:4) (2)
PI(P-18:0/20:4) (2)
PS(18:0/20:3)_A (2)
PE(P-16:0/16:0) (2)
PC(18:2/20:4) (1)
BATF (8)
ZBTB8B (3)
HAND1 (3)
ZNF492 (2)
TBX21 (1)
DMRTA2 (1)
ACSM5
AIM1
ANKRD22
APOC2
APOC4-APOC2
C2orf91
CALCR
CD69
CLEC12A
CLEC17A
CLECL1
CTSW
CX3CR1
DUSP27
EMR1
FCRL5
FGF20
GAPT
GLDN
GPR141
GPR55
GPR82
GYPE
HAMP
HLA-DQA1
HLA-DQB1
HLA-DRB5
IGJ
IGLL5
IL21R
ITK
MORC1
MS4A4E
OLR1
P2RY12
PADI2
PROX2
PRR15
PTCRA
SCIN
SIGLEC8
STON1-GTF2A1L
TMEM119
TNFRSF8
Multi-omic subtype
nmf1
nmf2
nmf3
IDH mutant
B
Gene Set Enrichment Analysis
positive regulation of immune response
adaptive immune response
immune response
positive regulation of immune system process
T cell activation
axon development
axonogenesis
cell morphogenesis involved in neuron differentiation
dendrite development
dendrite morphogenesis
0
1
2
Absolute normalized enrichment score
Direction
Downregulated
Upregulated
FDR
0.00011407
5.7037e-05
7.8636e-09
Gene count
50
100
150

**Supplementary Fig. 132. GBM module 23. A.** GBM module 23 colored by Z-scored expression (blue: -2; yellow: +2). TF, lipid and metabolite regulators with respective regulator scores above the module. Annotations at the bottom indicate multi-omic subtypes (proneural/nmf1: red; mesenchymal/nmf2: green; classical/nmf3: blue; IDH mutant: orange). **B.** Gene set enrichment analysis with GO biological process 2025 for module 23. **C.** Module 23 in the Lemonite KG. Metabolite regulators colored in red, TF regulators in green, module genes in orange. Nodes that connect metabolite regulators to TF regulators through a single PPI in the LemonIte KG colored in grey. Grey boxes represent genes involved in a protein-protein interaction that connects metabolite regulators to TF regulators in the Lemonite KG. Edges ending in a dot represent metabolic pathway interactions, normal lines represent PPIs and 'other' interactions.

GBM module 36 was significantly upregulated in mesenchymal-like (nmf2) samples (p-adj 3.30e-3) and was primarily expressed in monocytes and TAMs (main manuscript figure 4C). It was involved in immune response and defense response processes. Myo-inositol, creatinine and capric acid were identified as negative metabolic regulators, while FOXO6 and GFI1B were top transcriptional regulators. The module retrieved known interactions between metabolite regulators and module genes GPX3 and MGAM.

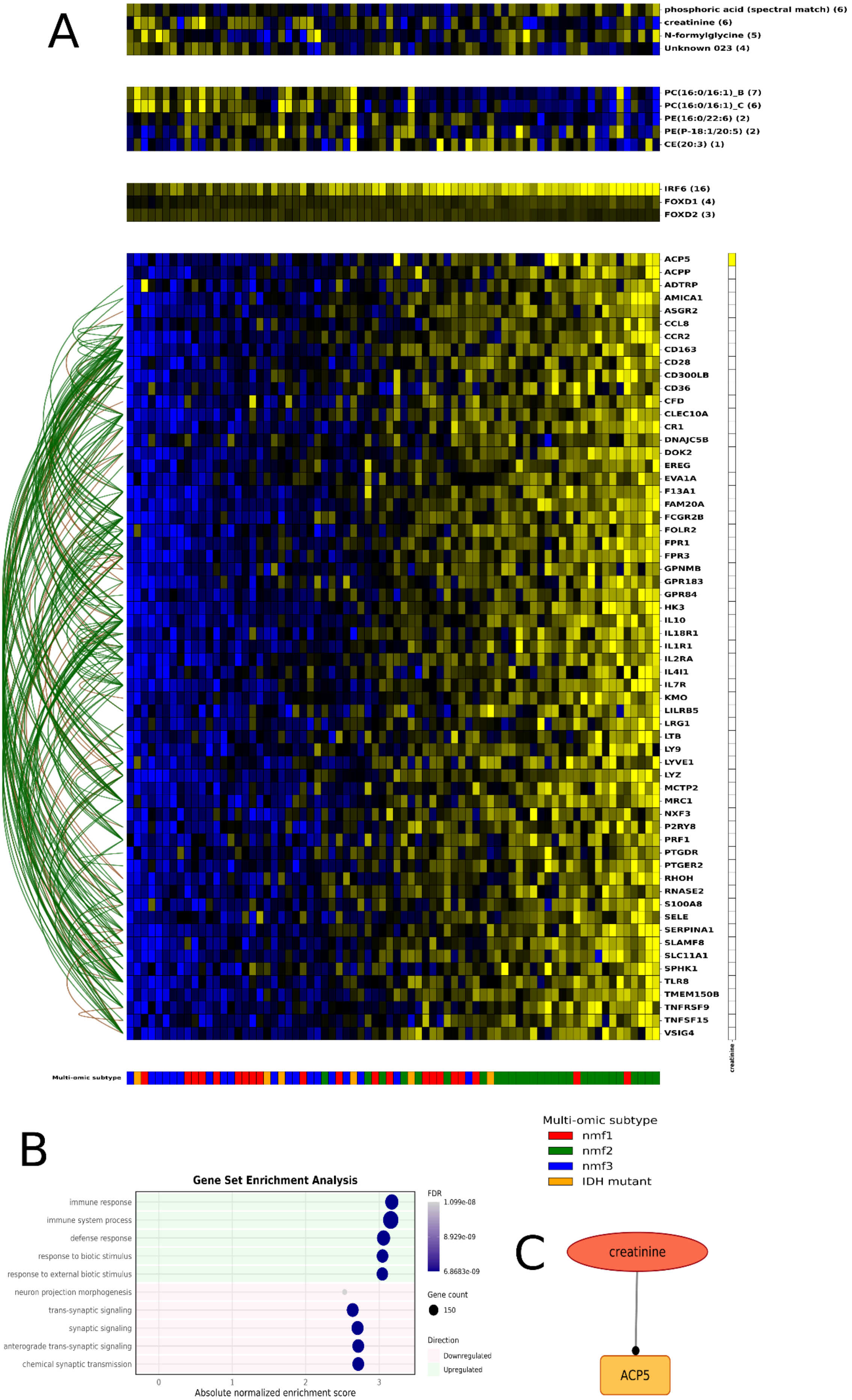
A
phosphoric acid (spectral match) (6)
creatinine (6)
N-formylglycine (5)
Unknown 023 (4)
PC(16:0/16:1)_B (7)
PC(16:0/16:1)_C (6)
PE(16:0/22:6) (2)
PE(P-18:1/20:5) (2)
CE(20:3) (1)
IRF6 (16)
FOXD1 (4)
FOXD2 (3)
ACP5
ACPP
ADTRP
AMICA1
ASGR2
CCL8
CCR2
CD163
CD28
CD300LB
CD36
CFD
CLEC10A
CR1
DNAJC5B
DOK2
EREG
EVA1A
F13A1
FAM20A
FCGR2B
FOLR2
FPR1
FPR3
GPNMB
GPR183
GPR84
HK3
IL10
IL18R1
IL1R1
IL2RA
IL4I1
IL7R
KMO
LILRB5
LRG1
LTB
LY9
LYVE1
LYZ
MCTP2
MRC1
NXF3
P2RY8
PRF1
PTGDR
PTGER2
RHOH
RNASE2
S100A8
SELE
SERPINA1
SLAMF8
SLC11A1
SPHK1
TLR8
TMEM150B
TNFRSF9
TNFSF15
VSIG4
creatinine
Multi-omic subtype
Multi-omic subtype
nmf1
nmf2
nmf3
IDH mutant
B
Gene Set Enrichment Analysis
immune response
immune system process
defense response
response to biotic stimulus
response to external biotic stimulus
neuron projection morphogenesis
trans-synaptic signaling
synaptic signaling
anterograde trans-synaptic signaling
chemical synaptic transmission
0
1
2
3
Absolute normalized enrichment score
FDR
1.099e-08
8.929e-09
6.8683e-09
Gene count
150
Direction
Downregulated
Upregulated
C
creatinine
ACP5

**3Supplementary Fig.14. GBM module 36**. **A.** GBM module 36 colored by Z-scored expression (blue: -2; yellow: +2). TF, lipid and metabolite regulators with respective regulator scores above the module. Annotations at the bottom indicate multi-omic subtypes (proneural/nmf1: red; mesenchymal/nmf2: green; classical/nmf3: blue; IDH mutant: orange). **B.** Gene set enrichment analysis with GO biological process 2025 for module 36. **C.** Module 36 in the Lemonite KG. Metabolite regulators colored in red, TF regulators in green, module genes in orange. Nodes that connect metabolite regulators to TF regulators through a single PPI in the LemonIte KG colored in grey. Grey boxes represent genes involved in a protein-protein interaction that connects metabolite regulators to TF regulators in the Lemonite KG. Edges ending in a dot represent metabolic pathway interactions, normal lines represent PPIs and 'other' interactions.

GBM module 50 was significantly upregulated in mesenchymal-like (nmf2) samples (p-adj 4.23e-5) ) and was primarily expressed in monocytes and TAMs (main manuscript Figure 4C). Accordingly, it was enriched for inflammatory response and immune system processes, including TNF signaling.

Myo-inositol was identified as a "negative" metabolic regulator, while IRF6 (with PEG3 and TFAP2A) was the top transcriptional regulator.

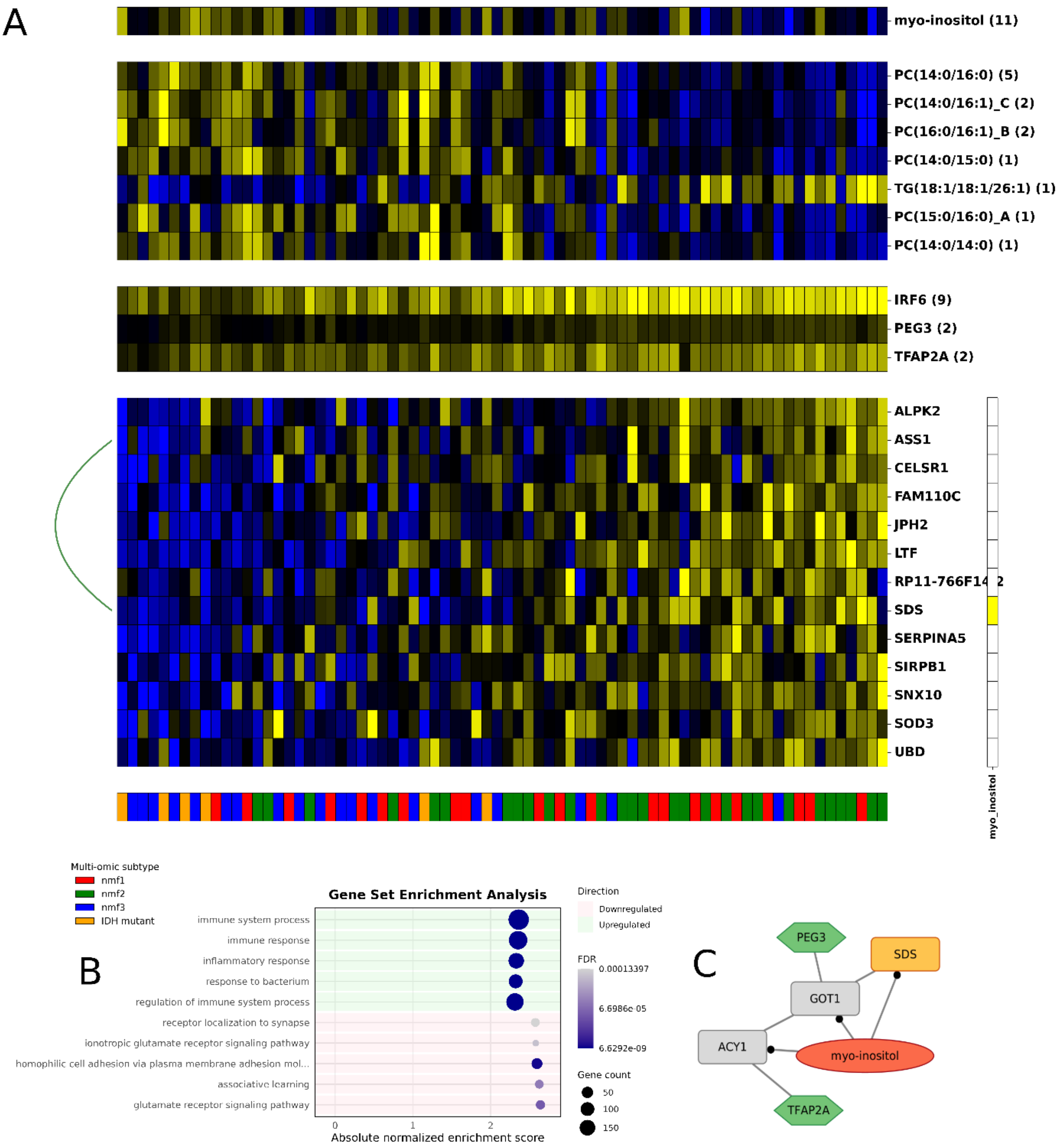

**4Supplementary Fig. 15. GBM module 50. A.** GBM module 50 colored by Z-scored expression (blue: -2; yellow: +2). TF, lipid and metabolite regulators with respective regulator scores above the module. Annotations at the bottom indicate multi-omic subtypes (proneural/nmf1: red; mesenchymal/nmf2: green; classical/nmf3: blue; IDH mutant: orange). **B.** Gene set enrichment analysis with GO biological process 2025 for module 50. **C.** Module 50 in the Lemonite KG. Metabolite regulators colored in red, TF regulators in green, module genes in orange. Nodes that connect metabolite regulators to TF regulators through a single PPI in the LemonIte KG colored in grey. Grey boxes represent genes involved in a protein-protein interaction that connects metabolite regulators to TF regulators in the Lemonite KG. Edges ending in a dot represent metabolic pathway interactions, normal lines represent PPIs and ‘other’ interactions.

GBM module 26 was significantly downregulated in mesenchymal-like (nmf2) samples (p-adj 2.27e-6) while showing a trend towards higher expression in proneural-like samples. Accordingly, this module was primarily expressed in oligodendrocyte progenitor cells (OPCs), OPC-like cells and NPC-like cells. Accordingly, genes in this module were involved in synaptic signaling and neuronal communication pathways, including ion transport (Supplementary Data 1). 2-hydroxyglutaric acid was the highest-scoring metabolite regulator, in line with the module showing high expression in the limited set of IDH-mutant samples.

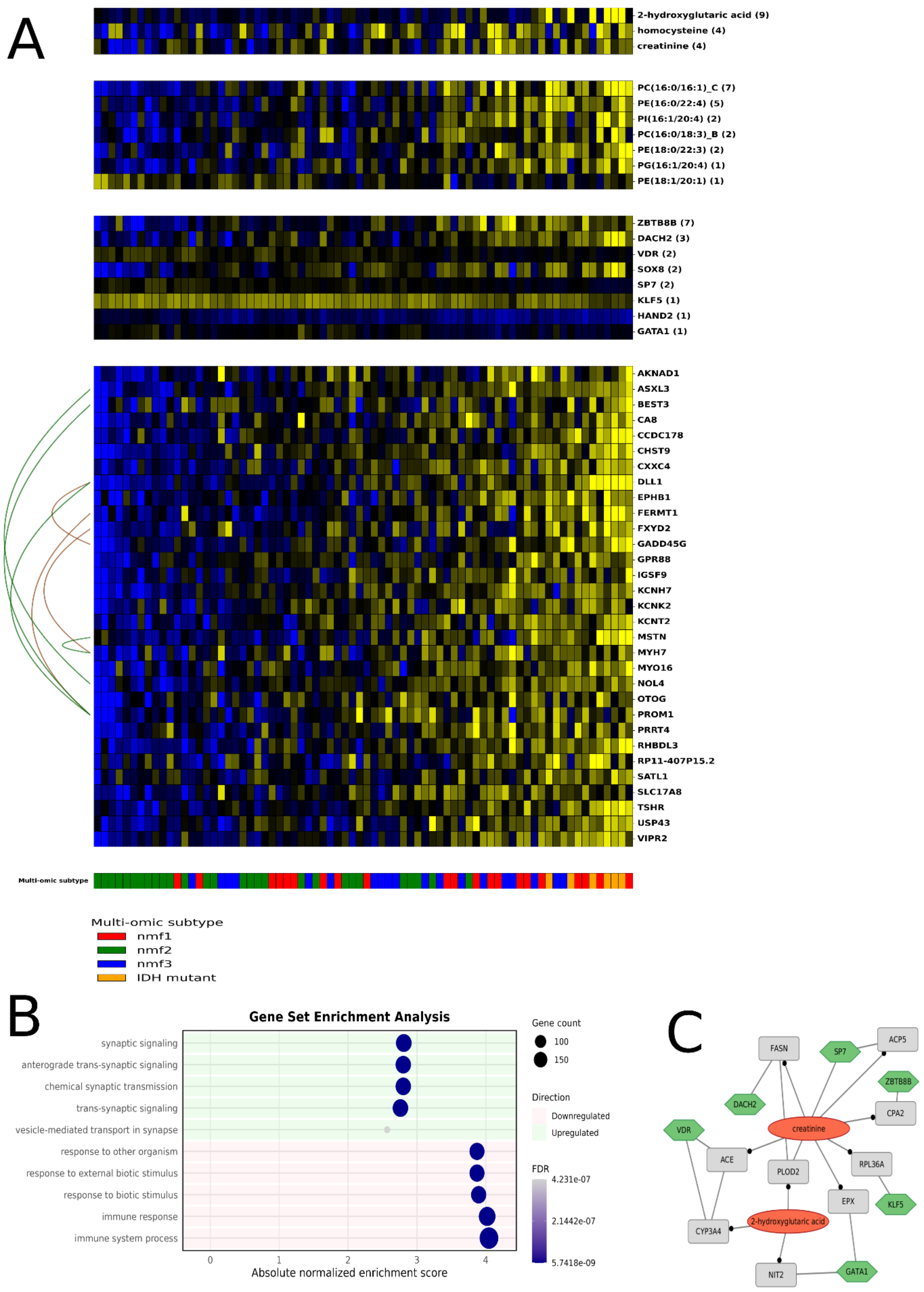

**5Supplementary Fig. 16. GBM module 26. A.** GBM module 26 colored by Z-scored expression (blue: -2; yellow: +2). TF, lipid and metabolite regulators with respective regulator scores above the module. Annotations at the bottom indicate multi-omic subtypes (proneural/nmf1: red; mesenchymal/nmf2: green; classical/nmf3: blue; IDH mutant: orange). **B.** Gene set enrichment analysis with GO biological process 2025 for module 26. **C.** Module 26 in the Lemonite KG. Metabolite regulators colored in red, TF regulators in green, module genes in orange. Nodes that connect metabolite regulators to TF regulators through a single PPI in the LemonIte KG colored in grey. Grey boxes represent genes involved in a protein-protein interaction that connects metabolite regulators to TF regulators in the Lemonite KG. Edges ending in a dot represent metabolic pathway interactions, normal lines represent PPIs and 'other' interactions.

GBM module 9 was significantly higher expressed in proneural (nmf1) and classical-like (nmf3) samples in comparison to mesenchymal-like samples (nmf2) (p-adj 4.24e-7, Kruskal–Wallis H test) and was significantly enriched for PPIs (p-adj 9.98e-16). Accordingly, genes in this module were primarily expressed in OPCs, OPC-like and NPC-like cells. Functionally, this module was associated with synaptic signaling and neuronal communication, including glutamatergic synapse activity and ion channel function (Supplementary Data 1). Homocysteine, D-malic acid and creatinine were identified as metabolic regulators, alongside multiple phosphatidylcholine and phosphatidylglycerol species, while SOX8, VAX2, GSX1 and ONECUT2 was predicted as negative transcriptional regulators and IRF6 as a positive transcriptional regulator. ONECUT2 is a well-known regulator of neuronal plasticity through remodeling of chromatin accessibility, while also homocysteine has been linked to OPC maturation through chromatin remodeling [4,5].

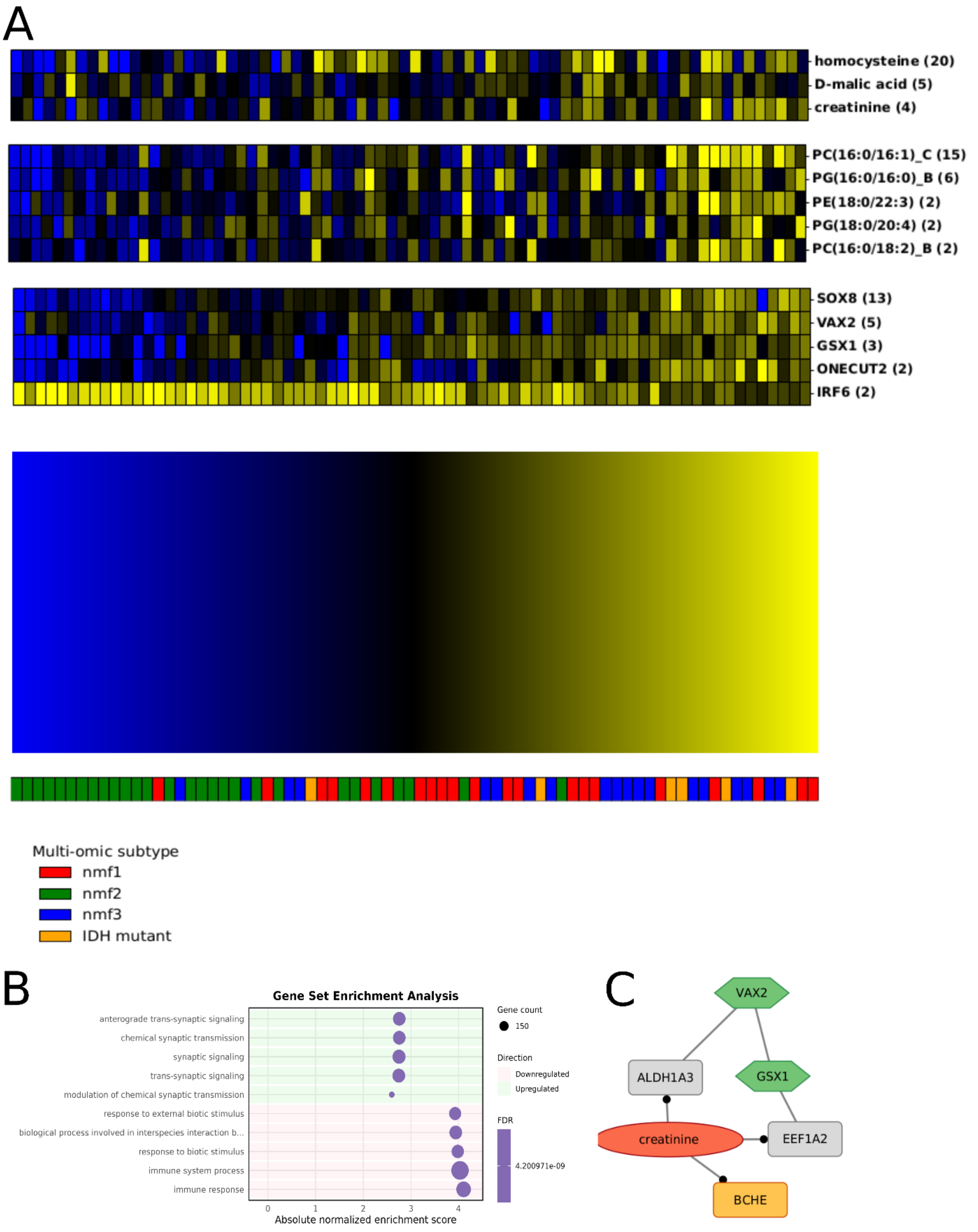

**Supplementary Fig. 17. GBM module 9. A.** GBM module 9 colored by Z-scored expression (blue: -2; yellow: +2). TF, lipid and metabolite regulators with respective regulator scores above the module. Annotations at the bottom indicate multi-omic subtypes (proneural/nmf1: red; mesenchymal/nmf2: green; classical/nmf3: blue; IDH mutant: orange). **B.** Gene set enrichment analysis with GO biological process 2025 for module 9. **C.** Module 9 in the Lemonite KG. Metabolite regulators colored in red, TF regulators in green, module genes in orange. Nodes that connect metabolite regulators to TF regulators through a single PPI in the LemonIte KG colored in grey. Grey boxes represent genes involved in a protein-protein interaction that connects metabolite regulators to TF regulators in the Lemonite KG. Edges ending in a dot represent metabolic pathway interactions, normal lines represent PPIs and ‘other’ interactions.

GBM module 2 was significantly upregulated in proneural-like (nmf1) samples (p-adj 1.23e-7) and showed strong PPI enrichment (p-adj 2.09e-38). Genes in module 2 were involved in synaptic signaling and neuronal system processes, in line with these genes primarily being expressed in OPC, OPC-like and NPC-like cells. Creatinine, L-cysteine and linoleic acid were identified as metabolite regulators, while CUX2 and MYTL1 both have established roles in glioblastoma [6,7].

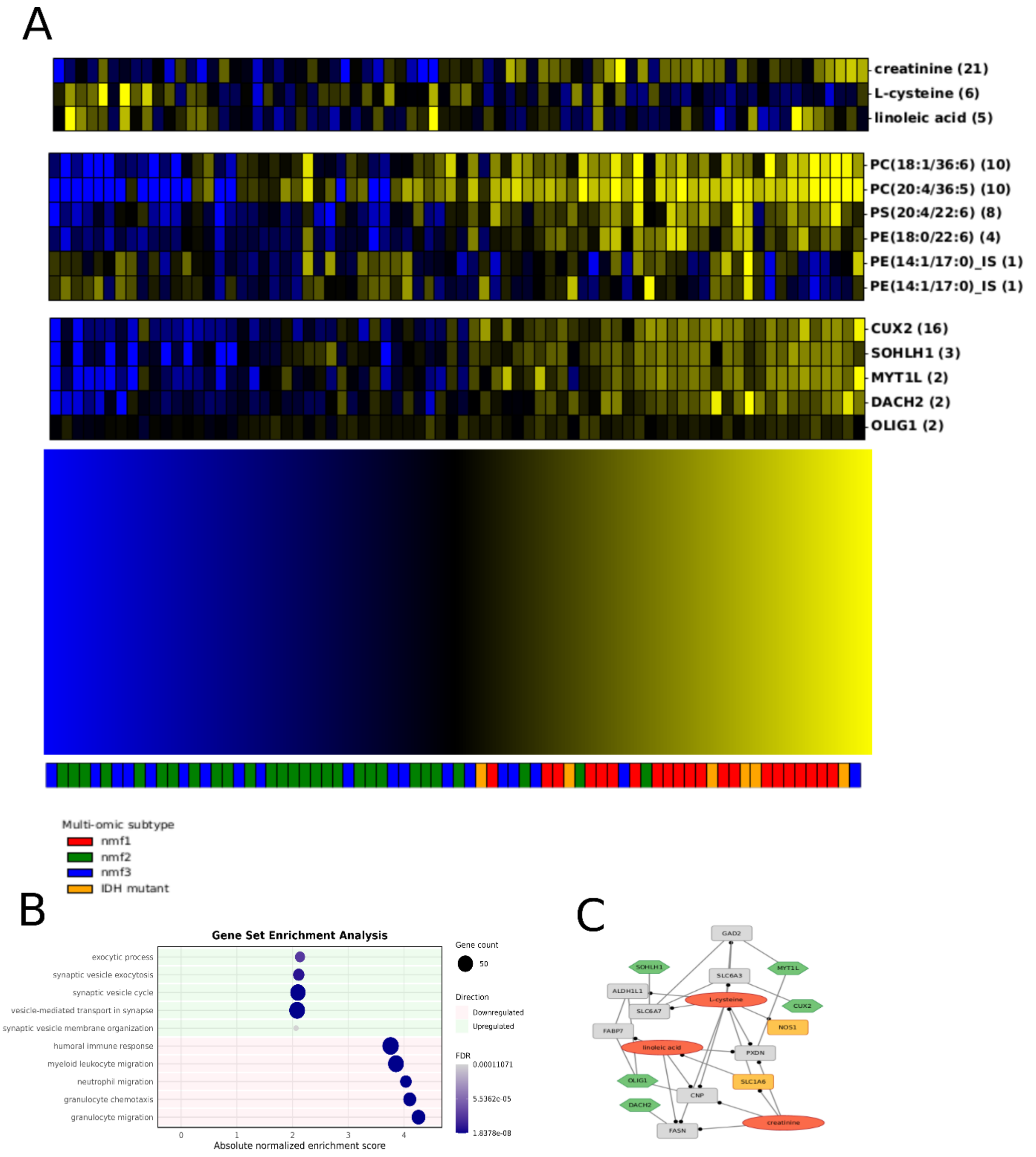

**Supplementary Fig. 18. 6 GBM module 2. A.** GBM module 2 colored by Z-scored expression (blue: -2; yellow: +2). TF, lipid and metabolite regulators with respective regulator scores above the module. Annotations at the bottom indicate multi-omic subtypes (proneural/nmf1: red; mesenchymal/nmf2: green; classical/nmf3: blue; IDH mutant: orange). **B.** Gene set enrichment analysis with GO biological process 2025 for module 2. **C.** Module 2 in the Lemonite KG. Metabolite regulators colored in red, TF regulators in green, module genes in orange. Nodes that connect metabolite regulators to TF regulators through a single PPI in the LemonIte KG colored in grey. Grey boxes represent genes involved in a protein-protein interaction that connects metabolite regulators to TF regulators in the Lemonite KG. Edges ending in a dot represent metabolic pathway interactions, normal lines represent PPIs and ‘other’ interactions.

GBM module 12 was significantly upregulated in proneural (nmf1) and IDH-mutant samples in comparison to other subtypes (p-adj 5.91e-7, Kruskal Wallis H test). Accordingly, module genes were mainly expressed in OPC, OPC-like and NPC-like cells, and functional enrichment analyses revealed this module to be involved in synaptic signaling and trans-synaptic communication.  L-cysteine, 2-hydroxyglutaric acid, glycerol-3-phosphate and maltose were identified as metabolite regulators, capturing known interactions with module genes FUT9, MACROD2, SLC16A7 and SLC25A21. TF regulators involved DACH2, ZNF727, GLIS1 and SOHLH1.

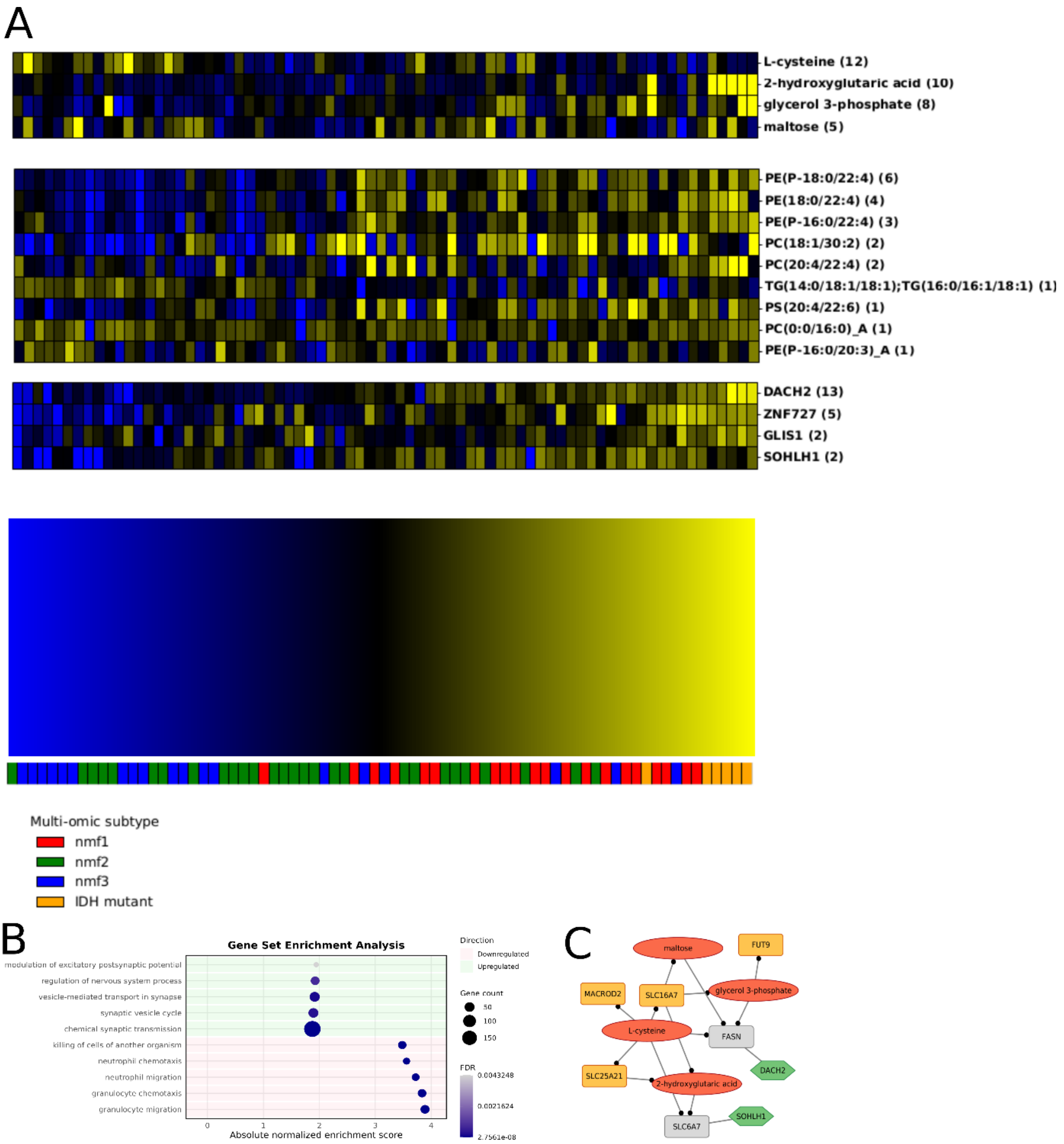

**Supplementary Fig. 19. GBM module 12. A.** GBM module 12 colored by Z-scored expression (blue: -2; yellow: +2). TF, lipid and metabolite regulators with respective regulator scores above the module. Annotations at the bottom indicate multi-omic subtypes (proneural/nmf1: red; mesenchymal/nmf2: green; classical/nmf3: blue; IDH mutant: orange). **B.** Gene set enrichment analysis with GO biological process 2025 for module 12. **C.** Module 12 in the Lemonite KG. Metabolite regulators colored in red, TF regulators in green, module genes in orange. Nodes that connect metabolite regulators to TF regulators through a single PPI in the LemonIte KG colored in grey. Grey boxes represent genes involved in a protein-protein interaction that connects metabolite regulators to TF regulators in the Lemonite KG. Edges ending in a dot represent metabolic pathway interactions, normal lines represent PPIs and 'other' interactions.

GBM module 62 was significantly differentially expressed between subtypes and showed higher expression in proneural (nmf1) samples (p-adj 1.21e-5, Kruskal-Wallis test). Accordingly, genes in module 62 were primarily expressed in OPC, OPC-like and NPC-like cells. Functional enrichment analyses revealed involvement in synaptic signaling processes (Supplementary Data 1). Creatinine and L-cysteine were identified as metabolic regulators, while SCRT1, SOX8, MYT1L, MYB and HAND2 were key transcriptional regulators. This module captured a known interaction between L-cysteine and GAD1.

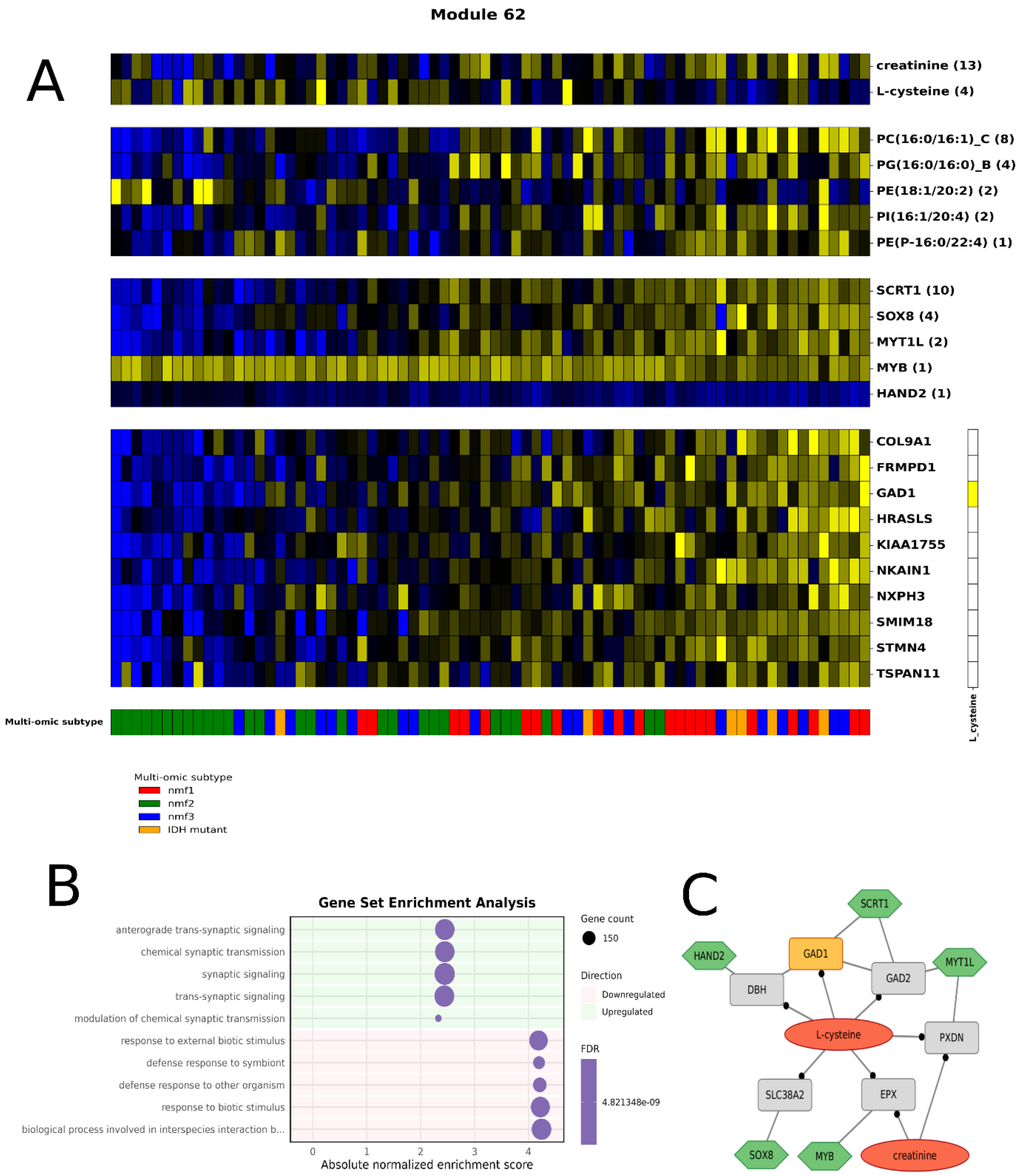

**7Supplementary Fig. 20. GBM module 62. A.** GBM module 62 colored by Z-scored expression (blue: -2; yellow: +2). TF, lipid and metabolite regulators with respective regulator scores above the module. Annotations at the bottom indicate multi-omic subtypes (proneural/nmf1: red; mesenchymal/nmf2: green; classical/nmf3: blue; IDH mutant: orange). **B.** Gene set enrichment analysis with GO biological process 2025 for module 62. **C.** Module 62 in the Lemonite KG. Metabolite regulators colored in red, TF regulators in green, module genes in orange. Nodes that connect metabolite regulators to TF regulators through a single PPI in the LemonIte KG colored in grey. Grey boxes represent genes involved in a protein-protein interaction that connects metabolite regulators to TF regulators in the Lemonite KG.

Edges ending in a dot represent metabolic pathway interactions, normal lines represent PPIs and ‘other’ interactions.

GBM module 17 showed differentially expressed (p-adj 9.20e-4, Kruskal-Wallis H test) between subtypes and showed higher expression in proneural samples. Accordingly, genes in this module were primarily expressed in OPC, OPC-like and NPC-like cells, and functional enrichment analyses revealed involvement in synaptic signaling and ion channel activity. 2-hydroxyglutaric acid and linoleic acid were identified as metabolite regulators, while TF regulators involved NRE2E1, DMRTA2, EGR4, SCRT1, SOX8, NKX2-2, MYTL1L and CUX2.

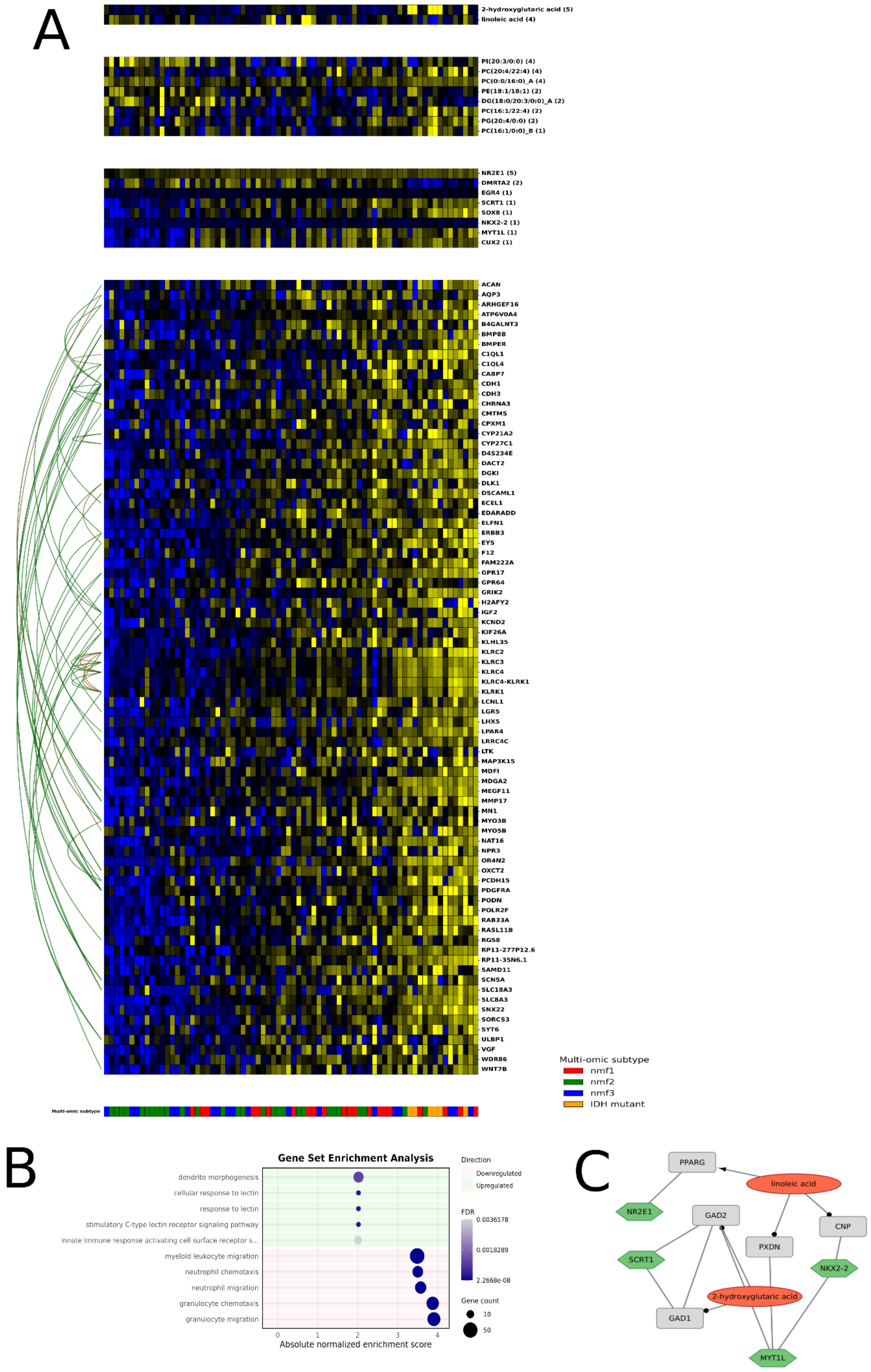
A
2-hydroxyglutaric acid (5)
linoleic acid (4)
PI(20:3/0:0) (4)
PC(20:4/22:4) (4)
PC(0:0/16:0)_A (4)
PE(18:1/18:1) (2)
DG(18:0/20:3/0:0)_A (2)
PC(16:1/22:4) (2)
PG(20:4/0:0) (2)
PC(16:1/0:0)_B (1)
NR2E1 (5)
DMRTA2 (2)
EGR4 (1)
SCRT1 (1)
SOX8 (1)
NKX2-2 (1)
MYT1L (1)
CUX2 (1)
ACAN
AQP3
ARHGEF16
ATP6V0A4
B4GALNT3
BMP8B
BMPER
C1QL1
C1QL4
CABP7
CDH1
CDH3
CHRNA3
CMTM5
CPXM1
CYP21A2
CYP27C1
D4S234E
DACT2
DGKI
DLK1
DSCAML1
ECEL1
EDARADD
ELFN1
ERBB3
EYS
F12
FAM222A
GPR17
GPR64
GRIK2
H2AFY2
IGF2
KCND2
KIF26A
KLHL35
KLRC2
KLRC3
KLRC4
KLRC4-KLRK1
KLRK1
LCNL1
LGR5
LHX5
LPAR4
LRRC4C
LTK
MAP3K15
MDFI
MDGA2
MEGF11
MMP17
MN1
MYO3B
MYO5B
NAT16
NPR3
OR4N2
OXCT2
PCDH15
PDGFRA
PODN
POLR2F
RAB33A
RASL11B
RGS8
RP11-277P12.6
RP11-35N6.1
SAMD11
SCN5A
SLC18A3
SLC8A3
SNX22
SORCS3
SYT6
ULBP1
VGF
WDR86
WNT7B
Multi-omic subtype
Multi-omic subtype
nmf1
nmf2
nmf3
IDH mutant
B
Gene Set Enrichment Analysis
dendrite morphogenesis
cellular response to lectin
response to lectin
stimulatory C-type lectin receptor signaling pathway
innate immune response activating cell surface receptor s...
myeloid leukocyte migration
neutrophil chemotaxis
neutrophil migration
granulocyte chemotaxis
granulocyte migration
0
1
2
3
4
Absolute normalized enrichment score
Direction
Downregulated
Upregulated
FDR
0.0036578
0.0018289
2.2668e-08
Gene count
10
50
C
PPARG
linoleic acid
NR2E1
GAD2
CNP
PXDN
SCRT1
NKX2-2
2-hydroxyglutaric acid
GAD1
MYT1L

**8Supplementary Fig. 21. GBM module 17. A.** GBM module 17 colored by Z-scored expression (blue: -2; yellow: +2). TF, lipid and metabolite regulators with respective regulator scores above the module. Annotations at the bottom indicate multi-omic subtypes (proneural/nmf1: red; mesenchymal/nmf2: green; classical/nmf3: blue; IDH mutant: orange). **B.** Gene set enrichment analysis with GO biological process 2025 for module 17. **C.** Module 17 in the Lemonite KG. Metabolite regulators colored in red, TF regulators in green, module genes in orange. Nodes that connect metabolite regulators to TF regulators through a single PPI in the LemonIte KG colored in grey. Grey boxes represent genes involved in a protein-protein interaction that connects metabolite regulators to TF regulators in the Lemonite KG. Edges ending in a dot represent metabolic pathway interactions, normal lines represent PPIs and ‘other’ interactions.

GBM module 53 was significantly differentially expressed between subtypes (p-adj 3.35e-4, Kruskal-Wallis H test) and module genes were primarily expressed in OPC, OPC-like and NPC-like cells. Accordingly, module genes were involved in synapse organization and chemical synaptic transmission (Supplementary Data 1). Metabolite regulators involved pyrophosphate, 2-hydroxyglutaric acid, homocysteine and heptadecanoic acid, while SOX8, CUX2 and ONECUT2, MYT1L, DACH2 and BARHL2 were assigned as TF regulators.

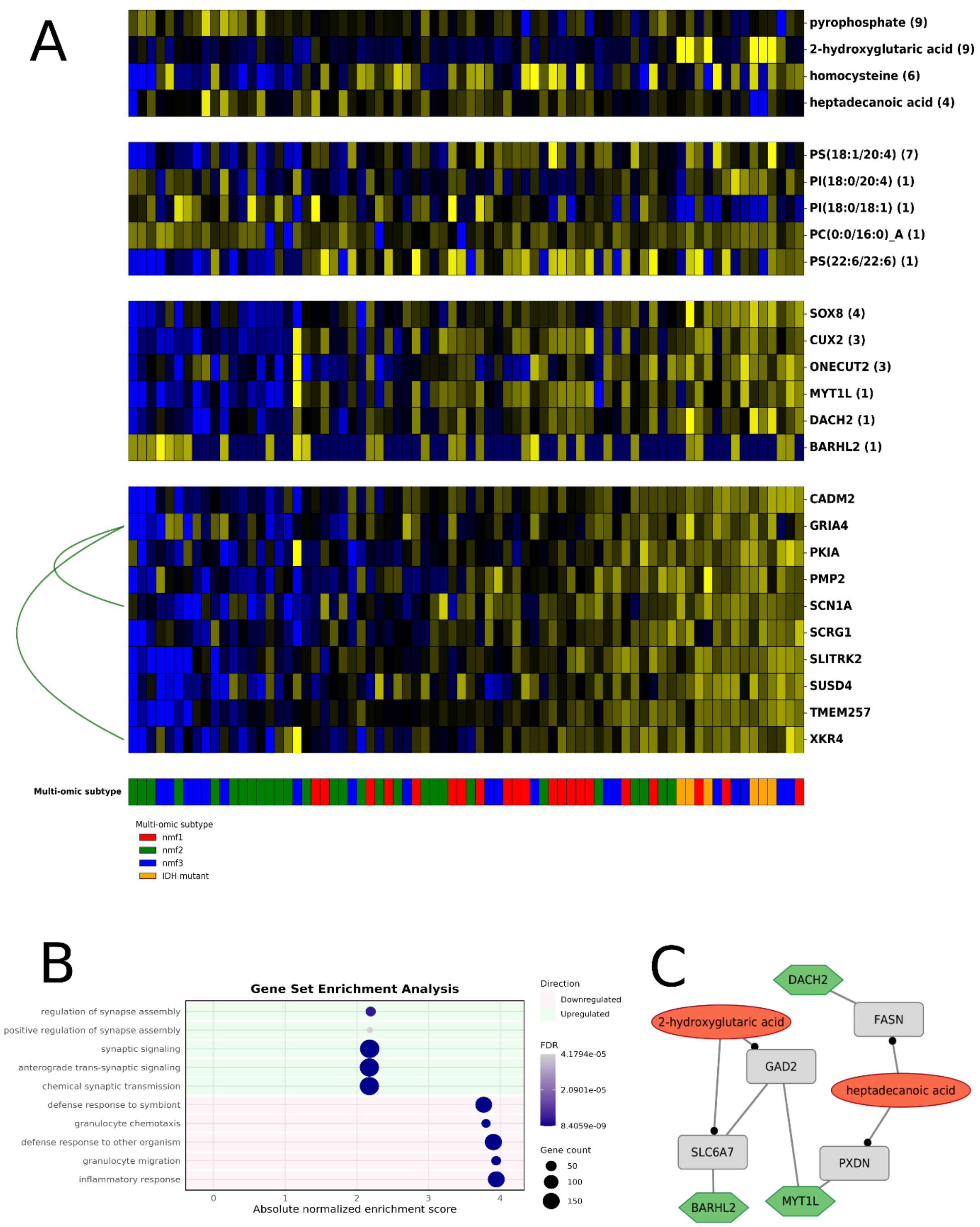

**Supplementary Fig. 22. 9GBM Module 53. A.** GBM module 53 colored by Z-scored expression (blue: -2; yellow: +2). TF, lipid and metabolite regulators with respective regulator scores above the module. Annotations at the bottom indicate multi-omic subtypes (proneural/nmf1: red; mesenchymal/nmf2: green; classical/nmf3: blue; IDH mutant: orange). **B.** Gene set enrichment analysis with GO biological process 2025 for module 53. **C.** Module 53 in the Lemonite KG. Metabolite regulators colored in red, TF regulators in green, module genes in orange. Nodes that connect metabolite regulators to TF regulators through a single

PPI in the LemonIte KG colored in grey. Grey boxes represent genes involved in a protein-protein interaction that connects metabolite regulators to TF regulators in the Lemonite KG. Edges ending in a dot represent metabolic pathway interactions, normal lines represent PPIs and ‘other’ interactions.

GBM Module 6 was significantly differentially upregulated in proneural (nmf1) samples (p-adj 3.25e-7, Kruskal-Wallis H test ) and showed strong PPI enrichment (p-adj 1.60e-22). Accordingly, genes in module 6 were primarily expressed in OPC, OPC-like and NPC-like cells. Functionally, module 6 was enriched for synaptic vesicle cycle and neurotransmitter signalling pathways (Supplementary Data 1). Creatinine, heptadecanoic acid and 2-hydroxyglutaric acid were identified as metabolite regulators, next to MYT1L and EMX1 as TF regulators. Furthermore, two phosphatidylcholine species were assigned as lipid regulators, as well as four phosphatidylethanolamines. The module retrieved known interactions between metabolite regulators and CMKT1A, CMKT1B, CYP4X1, ENTPD3, SLC27A2 and SLC4A10.

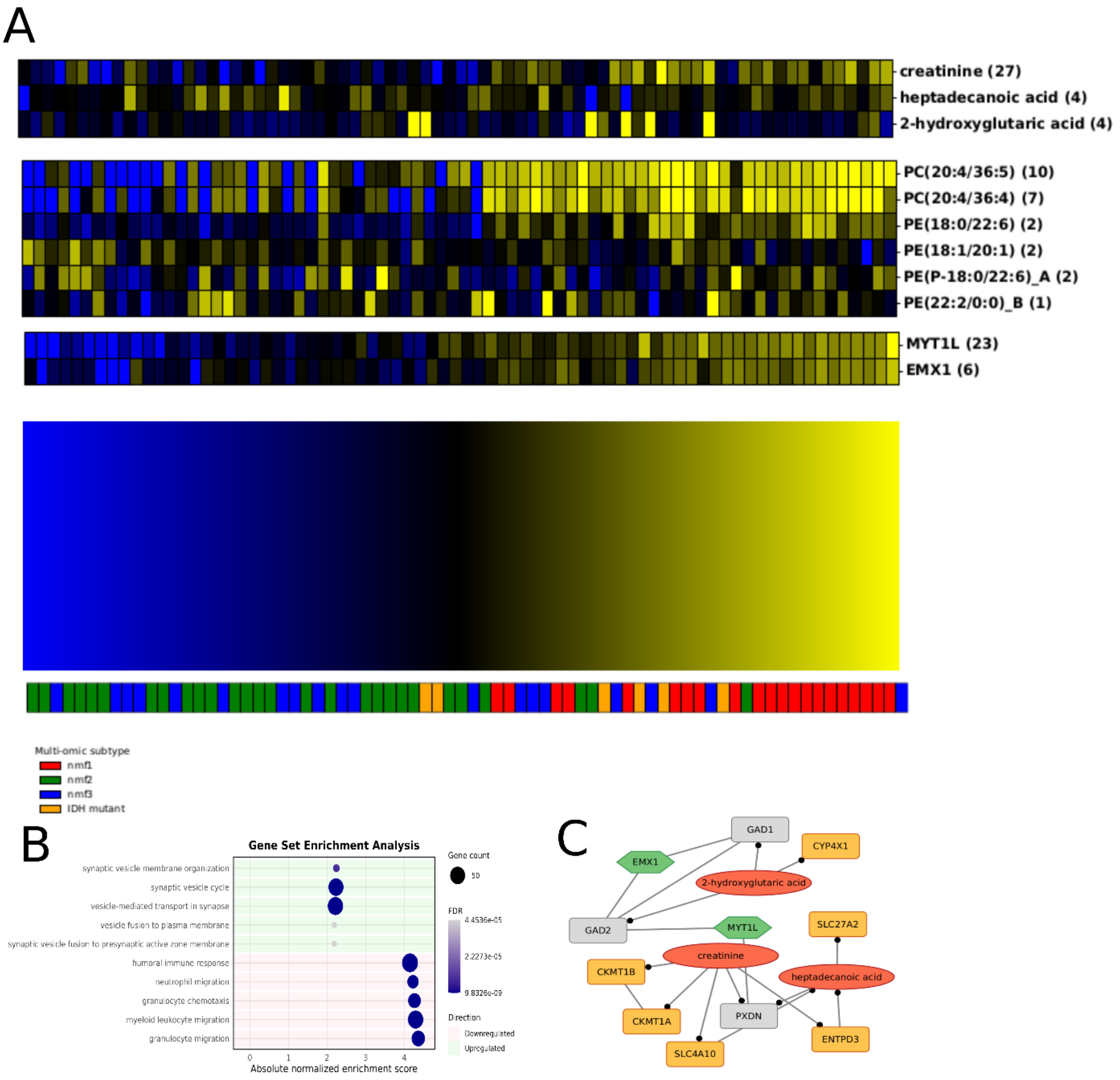

**10Supplementary Fig. 23. GBM Module 6. A.** GBM module 6 colored by Z-scored expression (blue: -2; yellow: +2). TF, lipid and metabolite regulators with respective regulator scores above the module. Annotations at the bottom indicate multi-omic subtypes (proneural/nmf1: red; mesenchymal/nmf2: green; classical/nmf3: blue; IDH mutant: orange). **B.** Gene set enrichment analysis with GO biological process 2025 for module 6. **C.** Module 6 in the Lemonite KG. Metabolite regulators colored in red, TF regulators in green, module genes in orange. Nodes that connect metabolite regulators to TF regulators through a single PPI in the LemonIte KG colored in grey. Grey boxes represent genes involved in a protein-protein interaction that connects metabolite regulators to TF regulators in the Lemonite KG. Edges ending in a dot represent metabolic pathway interactions, normal lines represent PPIs and ‘other’ interactions.

GBM Module 10 was significantly upregulated in mesenchymal-like (nmf2) samples, and primarily expressed in CD4/CD8 T cells. The module was significantly enriched for known PPIs (p-adj ~ 0, hypergeometric test) and was involved in adaptive immune response and T cell receptor signaling. Metabolite regulators involved DL-3-aminoisobutyric acid phosphoric acid, L-proline and an unknown metabolite, while TF regulators involved HAND1, NOTO, ZNF683 and ZBTB8B. ZNF683 is a well-established regulator of tissue-resident memory T cells, but has also been implicated in anti-tumor immunity following anti-PD1 therapy in leukemia [8].

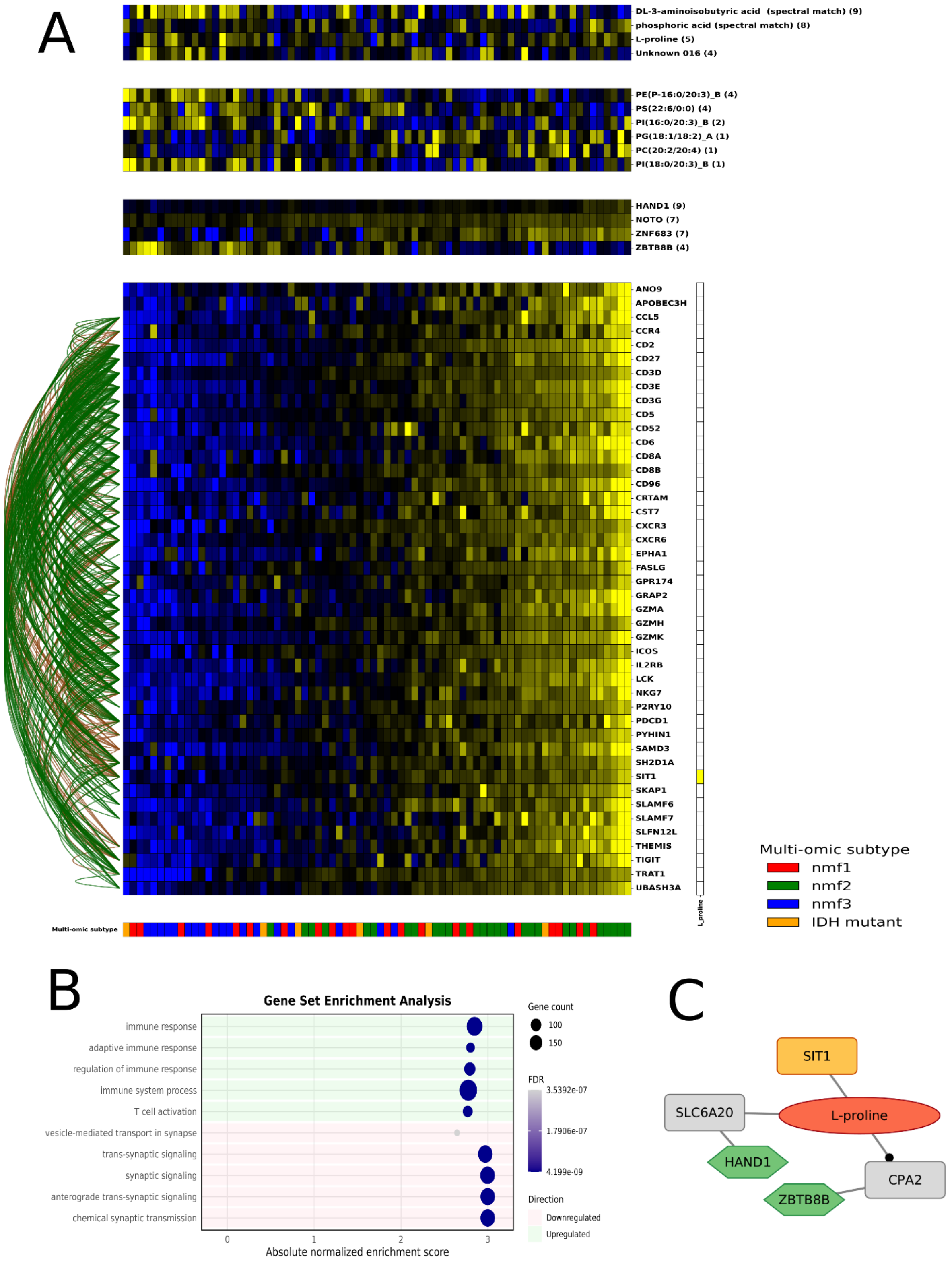

**11Supplementary Fig. 24. GBM Module 10. A.** GBM module 10 colored by Z-scored expression (blue: -2; yellow: +2). TF, lipid and metabolite regulators with respective regulator scores above the module. Annotations at the bottom indicate multi-omic subtypes (proneural/nmf1: red; mesenchymal/nmf2: green; classical/nmf3: blue; IDH mutant: orange). **B.** Gene set enrichment analysis with GO biological process 2025 for module 10. **C.** Module 10 in the Lemonite KG. Metabolite regulators colored in red, TF regulators in green, module genes in orange. Nodes that connect metabolite regulators to TF regulators through a single PPI in the LemonIte KG colored in grey. Grey boxes represent genes involved in a protein-protein interaction that connects metabolite regulators to TF regulators in the Lemonite KG. Edges ending in a dot represent metabolic pathway interactions, normal lines represent PPIs and 'other' interactions.

The modules that were selected for experimental validation in the IBD Lemonite network are described in more detail below.

IBD module 23 (Supplementary Fig. 6) comprised genes that were upregulated in UC relative to non-IBD controls (p-adj $2.28 \times 10^{-8}$, Kruskal-Wallis H test) and was significantly enriched for PPIs (p-adj $2.7 \times 10^{-14}$, hypergeometric test). STAT3 was the primary transcriptional regulator, followed by ERF2, ZFHX3 and ZNF704. Among the metabolite regulators, C18:1 LPC plasmalogen ranked first, alongside sphingomyelin and carnitine species.

IBD module 7 (Supplementary Fig. 7A) comprised genes that were upregulated in ulcerative colitis (UC) samples relative to non-IBD controls (p-adj $3.13 \times 10^{-8}$, Kruskal-Wallis H test; rank 6 of all modules), consistent with the strong PPI enrichment for this module (fold enrichment 6.9, p-adj $1.6 \times 10^{-255}$). The module was enriched for multiple inflammation-related pathways (Supplementary Fig. 7B, Supplementary Data 2). CEBPD was the top transcriptional regulator, followed by STAT3, IRF6, ZNF620 and ATF2. CEBPD is an IL-6/LPS-inducible transcription factor with a well-documented role in acute-phase and innate immune gene regulation, and CEBPD-deficient mice show enhanced susceptibility to DSS-induced colitis [9]. STAT3, the second regulator, is the canonical IL-6/IL-23–Th17 axis transcription factor central to IBD pathogenesis and a direct driver of several module genes [10]. The top metabolite regulators, C18:1 LPC plasmalogen, C2 carnitine, and TAG/DAG lipid species, point toward a lipid-remodeling and altered fatty-acid oxidation signature accompanying active inflammation.

IBD module 2 (Supplementary Fig. 8) comprised genes that were downregulated in UC relative to non-IBD controls (p-adj $2.23 \times 10^{-5}$, Kruskal-Wallis H test) and was significantly enriched for PPIs (p-adj $2.7 \times 10^{-14}$, hypergeometrical test). Module genes involved many genes encoding metabolic transporters (SLC26A3, AQP7, AQP8, ABCB1…) and accordingly shows enrichment for transmembrane transport-related pathways (Supplementary Data 2). ETV1 was the primary transcriptional regulator. Interestingly, ETV1 itself has recently been shown to be upregulated in IBD, to positively correlate with disease severity, and to drive CD4+ T cell activation and Th17 differentiation via the amino acid transporter SLC7A5 (member of module 23), in line with ETV1 activity but opposite to module expression [11]. The metabolite regulators (alpha-glycerophosphocholine, C2 carnitine, and several phosphatidylcholine, sphingomyelin and lysophospholipid species largely reflect general membrane phospholipid metabolism consistent with a mature, metabolically active absorptive epithelium.

IBD module 45 (Supplementary Fig. 9) comprised genes that were significantly downregulated in UC samples relative to non-IBD controls (p-adj $7.26 \times 10^{-7}$, Kruskal-Wallis H test). GSEA revealed that the module was involved in ‘estrogen metabolic process’ (upregulated), next to several downregulated inflammation-related pathways (p-adj $< 0.05$, GSEA, Supplementary Data 2). GSEA against KEGG database also highlighted several lower-ranked pathways that plausibly point toward a smoking-related signal, including chemical carcinogenesis, DNA adducts, taste transduction, and the NOD-like receptor signaling pathway (Supplementary Data 2). Trigonelline, an $NAD^+$ precursor and established marker of coffee consumption, was the top-ranked regulator, followed by hydroxycotinine, a nicotine metabolite and a well-validated biomarker of tobacco smoke exposure [12]. This is notable given the long-standing but mechanistically unexplained paradox in IBD whereby smoking exacerbates Crohn's disease yet is protective in ulcerative colitis [13]. Top TF regulators CREB3L2, MAFA, ZFHX3, BATF2 and DMRTA1 did not reveal any obvious relation to smoking behavior, although BATF2 expression correlates with active inflammation, mucosal immune cell infiltration and macrophage activation [14]. Interestingly, the 3th ranking metabolite regulator ornithine could be connected to TF regulator CREB3L2 through a PPI with SLC7A11 in the Lemonite KG, which has been linked to smoking behavior in GWAS studies [15].
